# A membrane-bound aromatic *O*-prenyltransferase catalyzes the last reaction step in citrus auraptene biosynthesis

**DOI:** 10.64898/2026.07.25.740671

**Authors:** Shuhei Matsushita, Ryosuke Munakata, Marwa Roumani, Alexandre Olry, Masaru Nakayasu, Alain Hehn, Tetsuya Matsukawa, Akifumi Sugiyama, Kazufumi Yazaki

## Abstract

Plants produce a variety of *O*-prenylated aromatics that exhibit biological activities beneficial to human health, and the presence of the *O*-prenyl moiety is often crucial to their functions. However, most aromatic *O*-prenylation genes remain unknown in plants. In this study, we report the molecular identification of an aromatic *O*-prenyltransferase (PT) involved in the biosynthesis of auraptene (7-geranyloxycoumarin), a citrus metabolite known for its preservative effect on human cognitive function. Based on *in silico* screening focusing on the membrane-bound PT family, *CpPT4* was isolated as a candidate from grapefruit (*Citrus* x *paradisi*), an auraptene-rich species. Enzymatic characterization demonstrated that recombinant CpPT4 specifically catalyzes umbelliferone 7-*O*-geranyltransferase activity to form auraptene, which differs from the enzymatic functions of known *O*-PTs. This enzyme also catalyzed aromatic *N*-prenylation to produce a new-to-nature auraptene analog. Regarding organ- and organellar-specific localization, it is strongly suggested that CpPT4 functions in the outer pericarp plastids, where auraptene is expected be formed. Furthermore, we found that *CpPT4* orthologs are widely distributed in citrus genomes. Intriguingly, mandarins and their descendant species possess dysfunctional orthologs, which is consistent with the low accumulation of auraptene and its downstream metabolites in these species. This study provides an example of the contribution of the UbiA superfamily to *O*-prenylated aromatic biosynthesis. Moreover, *CpPT4* can be useful as a tool in the synthetic biology-based production of auraptene and its analogs, as well as a molecular marker in the breeding of auraptene-rich citrus varieties.

## Introduction

Plants produce a remarkable variety of specialized metabolites that contribute to their adaptation to different environmental conditions. Among plant-specialized metabolites, over 600 molecules are classified as *O*-prenylated aromatic molecules that possess prenyl side chains attached to the aromatic core structures via C-O bonds (Munakata and Yazaki, 2024). Its chemodiversity is due to variations in the phenolic core structure (e.g., coumarins, flavonoids, xanthones, and aromatic alkaloids), prenyl chain length (e.g., dimethylallyl (C5), geranyl (C10), and farnesyl (C15) moieties), and prenylation position. Chemical modifications of prenyl side chains also enrich their chemical structures.

More than half of plant *O*-prenylated aromatics are classified as coumarins, with their main plant origins being Rutaceae, Apiaceae, and Asteraceae (Epifano et al., 2007; Murray, 2002). *O*-Prenylated aromatics are proposed to contribute to plant chemical defense against biotic threats, such as pathogens (Hamerski et al., 1990) and herbivores (Neal and Wu, 1994). In addition, various pharmacological and health-promoting effects have been reported for this metabolite group (Epifano et al., 2007). Studies on structure-activity relationships have demonstrated that the functions of *O*-prenylated aromatics are conferred or enhanced by the presence of a prenyl side chain (Adams et al., 2006; Murakami et al., 1997). The hydrophobic property enhances interactions with biomembranes (Hendrich et al., 2002) and target proteins (Sevrioukova, 2019) has been proposed as a mode of action of the *O*-prenylated aromatics.

In plant specialized metabolism, more than 100 genes belonging to the UbiA superfamily, a membrane-bound prenyltransferase (PT) gene family, have been identified as aromatic *C*-PTs that transfer prenyl side chains to aromatic molecules via C-C bonds. These genes have been isolated from various plant families (Munakata and Yazaki, 2024; Zhang et al., 2025), including Boraginaceae (Yazaki et al., 2002), Fabaceae (Sasaki et al., 2008), Cannabaceae (Tsurumaru et al., 2010), Rutaceae (Munakata et al., 2014), Apiaceae (Karamat et al., 2014; Munakata et al., 2016), and Hypericaceae (Ernst et al., 2024). These reported UbiA-type *C*-PTs also show diversity in enzymatic function, such as substrate specificity for prenyl acceptor and donor molecules and product specificity (regio-specificity). In contrast, there are few examples of the molecular identification of aromatic *O*-PT genes. Recently, two UbiA-type PT genes, *CpPT1* and *AkPT1,* were reported to encode a bergaptol 5-*O*-geranyltransferase (B5OGT) and a bergaptol 5,8-*O*-dimethylallyltransferase in grapefruit (*Citrus* × *paradisi*, Rutaceae) and *Angelica keiskei* (Apiaceae), respectively (Munakata et al., 2021). Later, aromatic *O*-PT activities were reported for UbiA-type PTs; however, these activities were detected as trace side reactions (Li et al., 2022) or in artificial substrate combinations (Tanaya et al., 2023). Currently, the poor understanding of aromatic *O*-PT genes has hindered the elucidation of the biosynthesis of plant *O*-prenylated aromatics.

Auraptene (7-geranyloxycoumarin) is a representative biosynthetically unresolved molecule. It was first isolated from the outer pericarp (flavedo) of *C. natsudaidai* in 1930 (Komatsu et al., 1930). Other citrus species also accumulate auraptene, with its content differing significantly by organ (Munakata et al., 2021) and species (Dugrand-Judek et al., 2015). This molecule is known for its bioactivities beneficial to human health, such as anti-inflammatory and anti-cancer effects (Tayarani-Najaran et al., 2021). A clinical study demonstrated that auraptene preserves cognitive function (Igase et al., 2018), leading to the commercialization of a citrus beverage with a high auraptene content (Fukuda et al., 2019). These bioactivities are also attractive in citrus breeding, as exemplified by the development of the auraptene-rich cultivar “Aurastar” (Omura and Shimada, 2016). In the biosynthesis of auraptene, its coumarin core structure, umbelliferone (7-hydroxycoumarin), is derived via the phenylpropanoid pathway. All enzymatic steps in the formation of umbelliferone have been characterized at the molecular level in Rutaceae, *e.g.* as cinnamic acid 4-hydroxylase (C4H) (Gravot et al., 2004) and *p-*coumaroyl CoA 2′-hydroxylase (C2′H) (Vialart et al., 2012). However, the gene encoding umbelliferone 7-*O*-GT (U7OGT or auraptene synthase) in the final step of the pathway remains unidentified (Fig. 1).

**Fig. 1.**
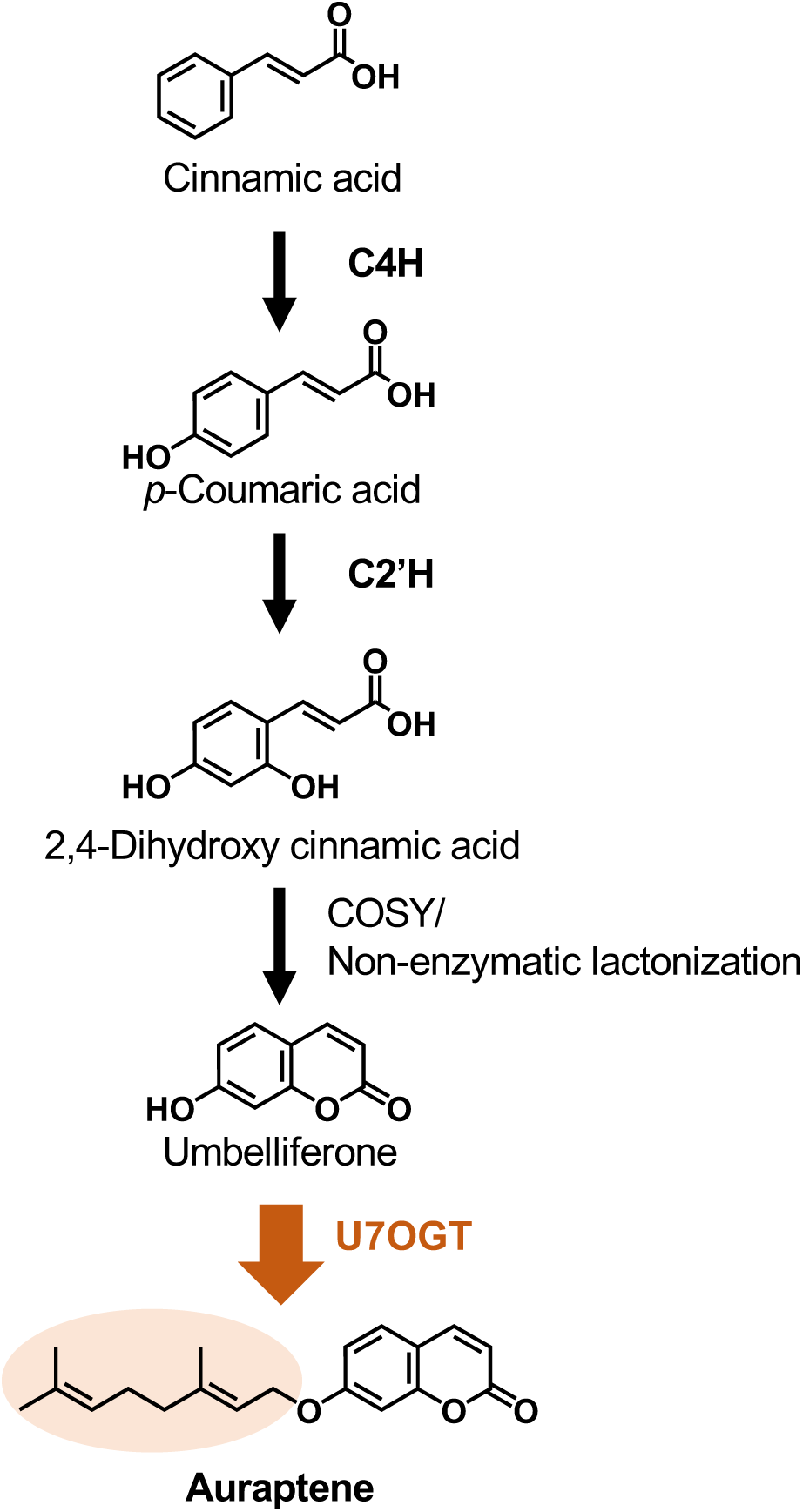
Proposed auraptene biosynthesis pathway in grapefruit. C4H, cinnamic acid 4-hydroxylase; C2’H, *p-* coumaroyl CoA 2′-hydroxylase; COSY, coumarin synthase; U7OGT, umbelliferone 7-*O*-geranyltransferase (auraptene synthase).

In this study, we attempted to identify an *auraptene synthase* from grapefruit, focusing on the UbiA superfamily, which has expanded in line with the high chemical diversity of prenylated aromatics in citrus (Munakata et al., 2021). A candidate gene was *in silico* screened based on transcriptomic data of grapefruit flavedo (Munakata et al., 2021) and the genome of pummelo (*C. grandis*), the female parent of grapefruit. This gene was functionally characterized using multifaceted approaches. In particular, the contribution of this gene to auraptene production in citrus was assessed by comparing the predicted intactness of the ortholog with the metabolite profiles of different species of this taxon.

## Results

### Isolation of a candidate gene encoding auraptene synthase from grapefruit

To identify the auraptene synthase gene, the transcriptome of the outer pericarp of grapefruit was utilized, in which this compound accumulates in a high concentration. Since biochemical analysis using native enzymes from the grapefruit pericarp suggested that the auraptene synthase is a UbiA-type protein, we selected candidate contigs that encode UbiA polypeptides to obtain 33 contigs from the RNA-seq data (Munakata et al., 2021). Subsequent to the exclusion of contigs associated with primary metabolism and the consideration of expression levels, four candidates were identified. As demonstrated in our previous study, the enzyme activities of three of them have already been evaluated (Munakata et al., 2021); however, none of these four clones exhibited auraptene synthesis activity. Among them, only contig c22985_g1_i1 was not characterized in detail because its gene product lacks the second aspartate-rich motif that is essential for the catalytic function of UbiA-type PT (Fig. S1). In this study, we began with a detailed evaluation of this candidate as an auraptene synthase using the public genome of pummelo (Wang et al., 2017). A manual exon-intron gene structure assessment of the genomic region corresponding to c22985_g1_i1 suggested that an intact coding sequence (CDS) could be transcribed by alternative splicing (Fig. S1). This assumption was confirmed by reverse transcription (RT)-PCR-based isolation of the intact CDS from grapefruit immature flavedo, where auraptene is highly accumulated (Munakata et al., 2021). This gene was named *C.* × *paradisi PT4* (*CpPT4*). In this experiment, we obtained seven amplicons from this tissue, and all of them maintained the nucleotide region encoding the second aspartate-rich motif. The splicing variant lacking this motif is probably minor in grapefruit immature flavedo.

*In silico* analysis of the polypeptide sequence of CpPT4 using the deepTMHMM (Hallgren et al., 2022) server predicted the presence of nine transmembrane regions, and TargetP 2.0 (Almagro Armenteros et al., 2019) and DeepLoc (Thumuluri et al., 2022) servers predicted 67 amino acids at the N-terminus as a transit peptide (Fig. S2). These polypeptide features are well conserved among plant UbiA-type PTs. In the phylogenetic tree, CpPT4 formed a clade with aromatic PTs involved in specialized metabolism in Rutaceae (Fig. 2 and Table S1), suggesting that CpPT4 is specific to this plant family. Among the reported aromatic PTs, CpPT4 showed the highest amino acid identity with CpPT3, the grapefruit U8GT (Fig. S2), which catalyzes C-C bond formation, but the identity value was moderate (58.3%), indicating the possibility that the enzymatic function of CpPT4 may differ from that of CpPT3.

**Fig. 2.**
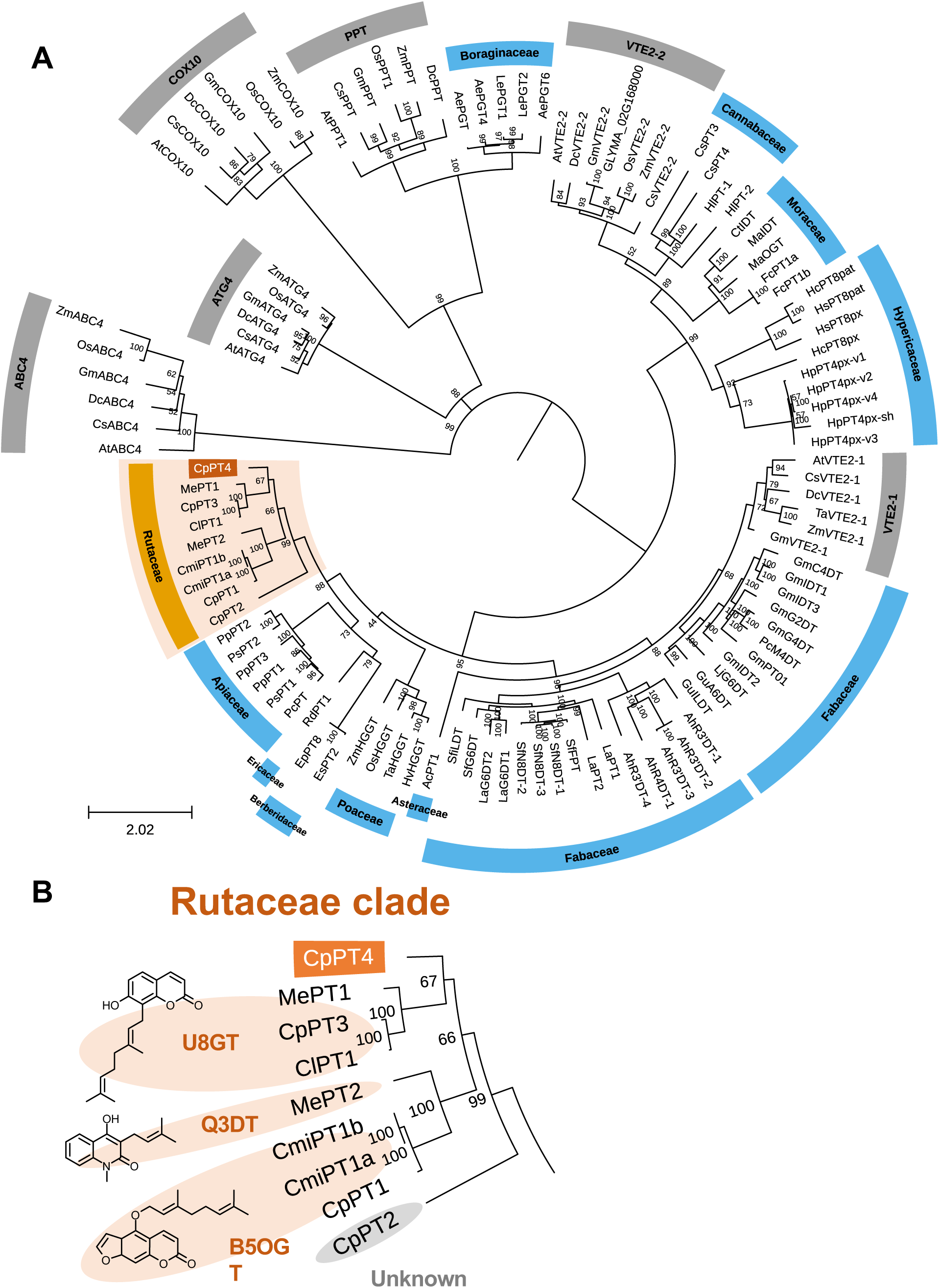
Phylogenetic analys is of CpPT4 within the UbiA superfamily. (A) A phylogenetic tree was constructed using the maximum likelihood method based on a MUSCLE multiple alignment of UbiA PTs, with 1,000 bootstrap replicates to assess branch support. Bootstrap values are indicated by the nodes separating the clades. Clades associated with primary metabolism are shaded in gray, whereas those related to specialized (secondary) metabolism are color-coded: orange for PTs from the Rutaceae family and light blue for PTs from other plant families. CpPT4 is indicated by an orange box. The accession numbers of the PTs included in the analysis are listed in Table S1. (B) Enlarged view of the Rutaceae clade. The main enzymatic activities and chemical structures of the products are shown. U8GT, umbelliferone 8-geranyltransferase; Q3DT, quinolone 3-dimethylallyltransferase; B5OGT, bergamottin 5-*O*-geranyltransferase.

### The U7OGT activity of recombinant CpPT4

Heterologous production of UbiA-type PTs in microbial systems often fails because of their membrane-bound properties (Karamat et al., 2014), whereas *Nicotiana benthamiana* has been used to functionally produce various UbiA-type PT proteins (Han et al., 2025; Munakata et al., 2021). Thus, *CpPT4* was transiently expressed in *N. benthamiana* leaves by agroinfiltration, and microsomes prepared from the leaves producing recombinant CpPT4 were used as crude enzyme sources. First, a preliminary *in vitro* screening was performed using 64 substrate pairs in the presence of Mg^2+^ as a cofactor (Table S2 and Fig. S3). When recombinant CpPT4 was incubated with umbelliferone and geranyl diphosphate (GPP) as prenyl acceptor and donor substrates, respectively, a single enzymatic product was detected by LC-PDA analysis (Fig. 3A). The enzymatic reaction product was identical to a standard specimen of auraptene in terms of retention time, UV spectrum, and mass spectrometry (MS) profiles (Fig. 3). In particular, MS^2^ analysis of both the reaction product and auraptene standard detected the neutral loss of 136 Da as the major fragmentation from the molecular ion peak of auraptene (m/z = 299) in the positive ion mode. This fragmentation corresponds to the total loss of a geranyl moiety and is probably specific to *O*-geranyl moiety attached to the aromatic ring because the neutral losses of 136 and 124 Da were observed in the MS^2^ analysis of other *O*-geranylated coumarin molecules (Munakata et al., 2021) and *C*-geranylated coumarins (Munakata et al., 2014), respectively.

**Fig. 3.**
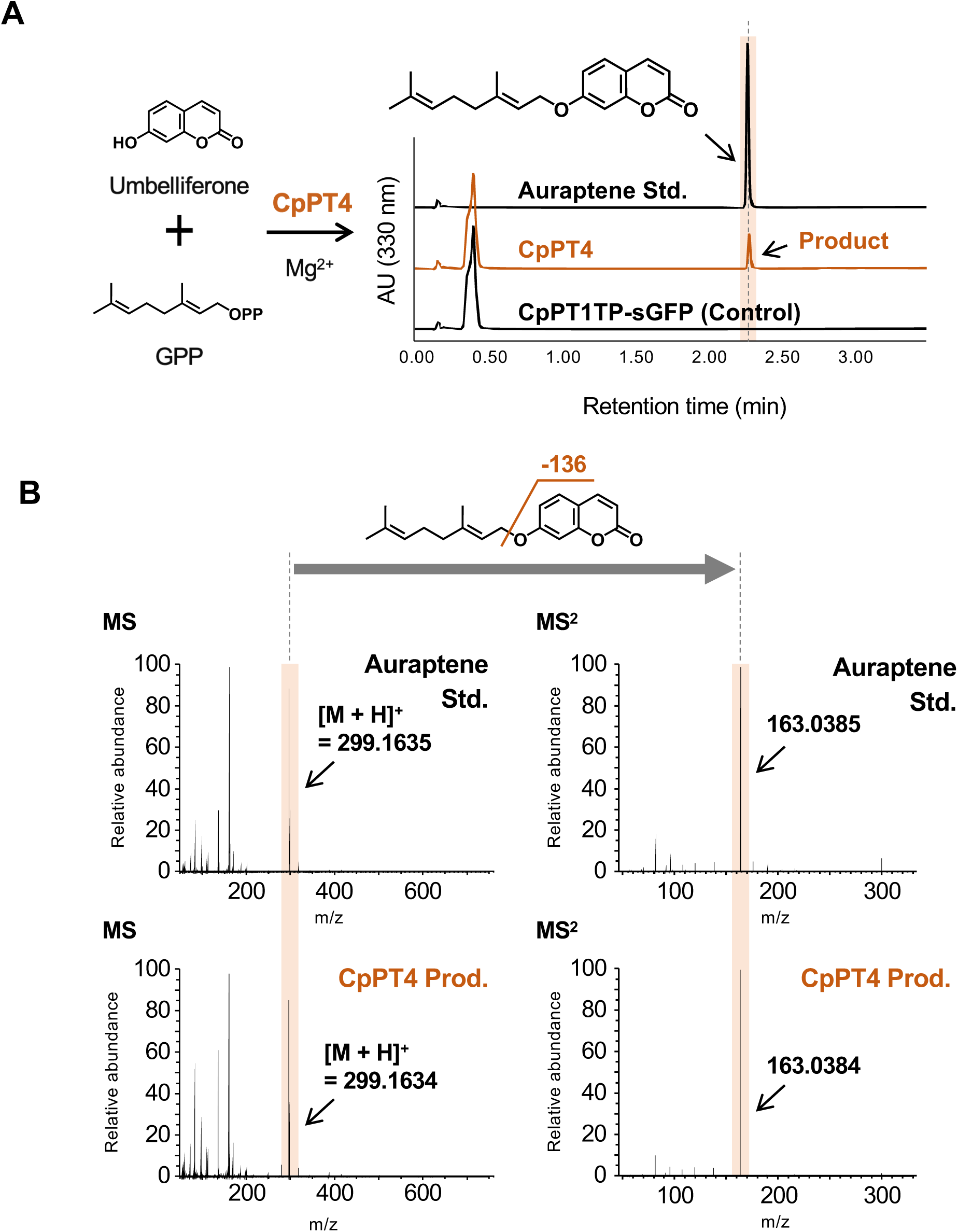
The U7OGT activity of recombinant CpPT4. (A) Ultraviolet chromatograms at 330 nm of the U7OGT reaction mixtures containing recombinant CpPT4. As a negative control, crude enzyme preparations expressing CpPT1TP-sGFP were used, and the corresponding chromatogram is displayed at the same scale as that of CpPT4 for comparison. (B) MS^2^ spectrum of the enzymatic reaction product acquired in the positive ion mode. The observed loss of 136 daltons was attributed to the cleavage of the *O*-geranyl moiety from the coumarin structure.

To confirm that the synthesis of auraptene was due to the enzymatic reaction of CpPT4, we tested several negative control assays, *i.e.*, incubation with microsomes containing synthetic green fluorescent protein (sGFP), heat-denatured microsomes, or EDTA, and incubation devoid of prenyl donor substrate, prenyl acceptor substrate, or microsomes. None of these conditions resulted in the production of enzymatic products (Fig. 3 and Fig. S4A). The reaction without MgCl_2_ yielded a trace amount of auraptene probably because metal ions were carried over from *N. benthamiana* during microsomal preparation (Fig. S4A). These results indicate that recombinant CpPT4 regio-specifically catalyzes 7-*O*-geranylation for umbelliferone. The U7OGT reaction with different divalent cations showed that Mg^2+^ was the best cofactor (Fig. S4B). The optimal pH was approximately 8.0, and the optimal temperature was around 28 °C (Fig. S4C and D). The apparent *K*_m_ values for umbelliferone and GPP were calculated to be 21 ± 1 and 12 ± 1 µM, respectively (Fig. S4E). These enzymatic properties resemble those of reported plant UbiA-type PTs (Karamat et al., 2014; Munakata et al., 2021, 2019).

### Substrate specificity of recombinant CpPT4

The prenyl acceptor specificity of CpPT4 was analyzed using GPP as a prenyl donor substrate in triplicate reactions (Fig. 4 and Fig. S3). When CpPT4 was incubated with coumarin molecules with different hydroxylation patterns, an enzymatic product was formed for 5,7-dihydroxycoumarin (no. 3 in Fig. 4) at a lower efficiency than that for umbelliferone (7-hydroxycoumarin, no. 1). Other molecules without hydroxy moieties at the C7 position were not accepted by CpPT4 (nos. 2, 7, 8, 9, 10, 12, and 13). This recognition pattern is consistent with the acceptance of scopoletin (6-methoxy-7-hydroxycoumarin, no. 6) and esculetin (6,7-dihydroxycoumarin, no. 4) as prenyl acceptors in the preliminary analysis (Table S2). The enzymatic products of 5,7-dihydroxycoumarin, scopoletin, and esculetin were all predicted to be *O*-geranylated molecules, based on the fragmentation pattern of the neutral loss of 136 Da in the MS^2^ analysis (Fig. S5).

**Fig. 4.**
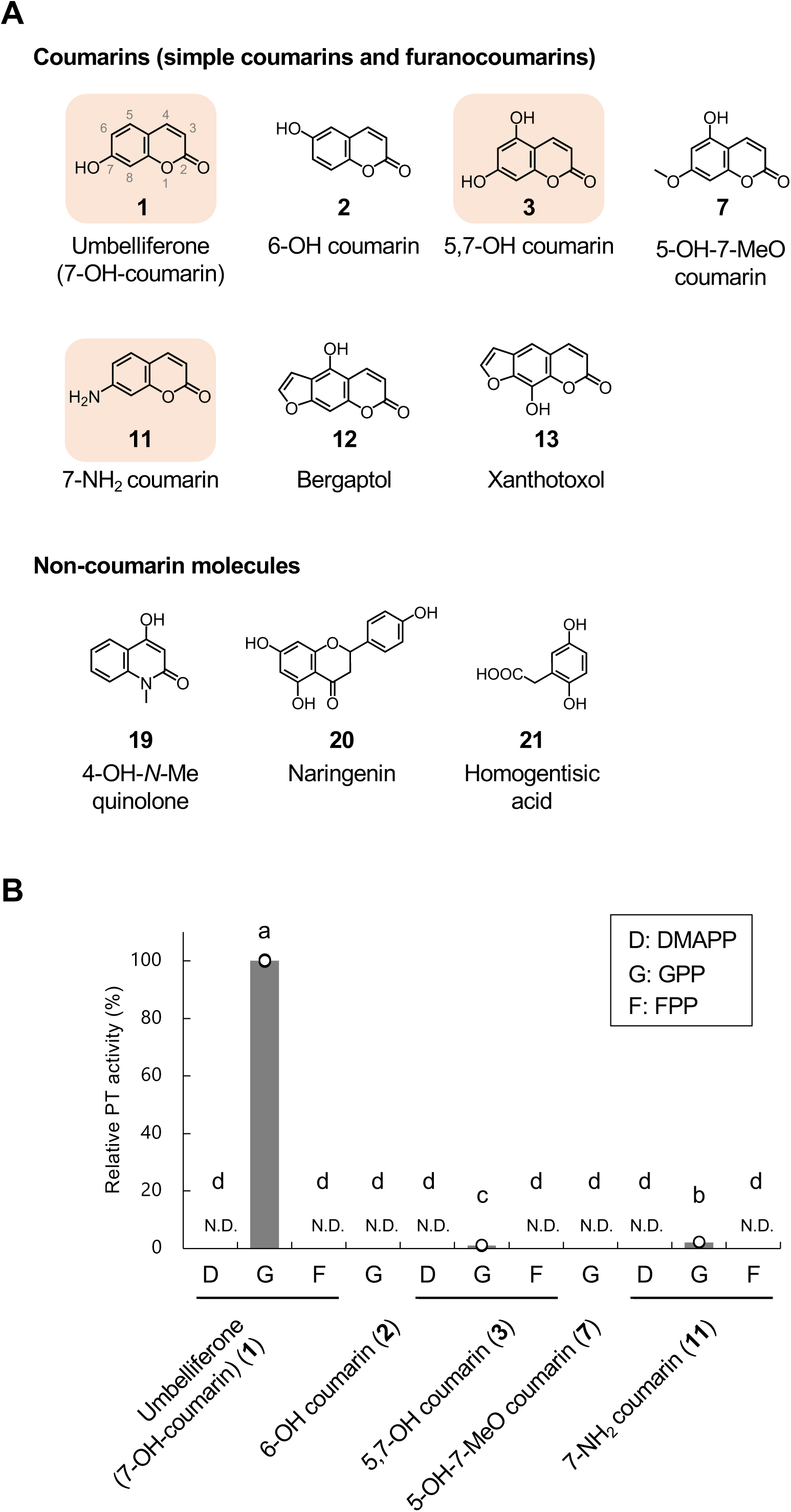
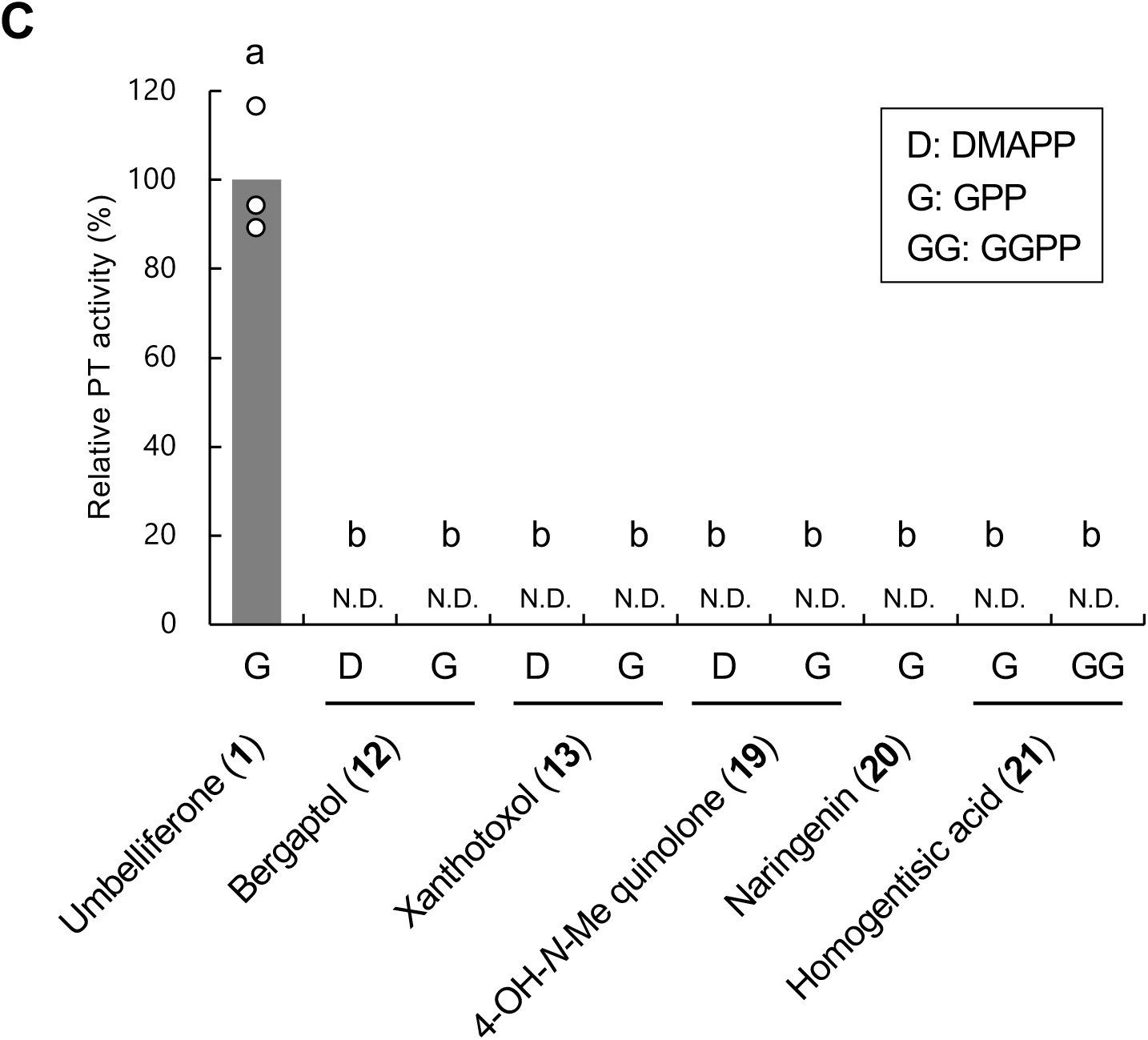
Substrate specificity of recombinant CpPT4. (A) Chemical structures of the selected prenyl acceptor substrates. Substrates that accept prenyl moieties are highlighted in orange. (B) Enzymatic activity toward simple (bicyclic) coumarin molecules. (C) Enzymatic activities toward various other phenolic groups. Activities are shown relative to the average U7OGT activity of the corresponding group. All the reactions were performed in triplicate. N.D., not detected. Significant differences among groups were evaluated using Tukey’s test (p < 0.05), and distinct letters indicate significant differences.

In most aromatic prenylations by UbiA superfamily proteins, prenyl carbocations generated from prenyl diphosphates electrophilically attack aromatic molecules that possess an electron-donating hydroxyl moiety attached to the aromatic ring (Cheng and Li, 2014), This led us to test another electron-donating group, the amino moiety. When 7-aminocoumarin (no. 11) was used as the prenyl acceptor substrate, CpPT4 produced the unnatural 7-geranylaminocoumarin (Fig. 4 and Fig. S5). To the best of our knowledge, this is the first example of aromatic *N*-prenylation activity in the UbiA superfamily. We also tested aromatic molecules other than coumarins, *i.e.* as naringenin (a flavanone molecule), quinolones, and homogentisic acid, considering the prenylated aromatic molecules present in Rutaceae species (Chun et al., 2006; Ito et al., 1988; Li et al., 2022). However, no reaction products were detected for these non-coumarin molecules (Fig. 4 and Table S2).

The prenyl donor specificity of CpPT4 was also investigated using prenyl diphosphates of different chain lengths, dimethylallyl diphosphate (DMAPP), GPP, and farnesyl diphosphate (FPP), in the presence of the aromatic molecules accepted in the GT assays. No reaction products were obtained in any combination except for GPP (Fig. 4 and Table S1). Our substrate specificity analysis demonstrated that CpPT4 accepts GPP and coumarins with electrophilic moieties, such as the hydroxyl and amino moieties, at the C7 position as prenyl donor and acceptor substrates, respectively.

### In planta localization of CpPT4

The expression levels of *CpPT4* were quantified in different grapefruit organs using reverse transcription-quantitative polymerase chain reaction (RT-qPCR) (Fig. 5 and Fig. S6). In both immature and mature fruits, the expression of *CpPT4* was clearly observed in the flavedo but not in the inner white peel (albedo). Buds and leaves showed similar or lower expression levels than those in flavedo tissues. This expression pattern is similar to that of auraptene and its downstream metabolite epoxyauraptene (Munakata et al., 2021). Hereafter, auraptene and auraptene-derived metabolites are collectively referred to as auraptenes. We also observed higher expression levels of this gene in mature flavedo than in immature flavedo, which differs from the accumulation pattern of auraptenes in these two flavedo samples.

**Fig. 5.**
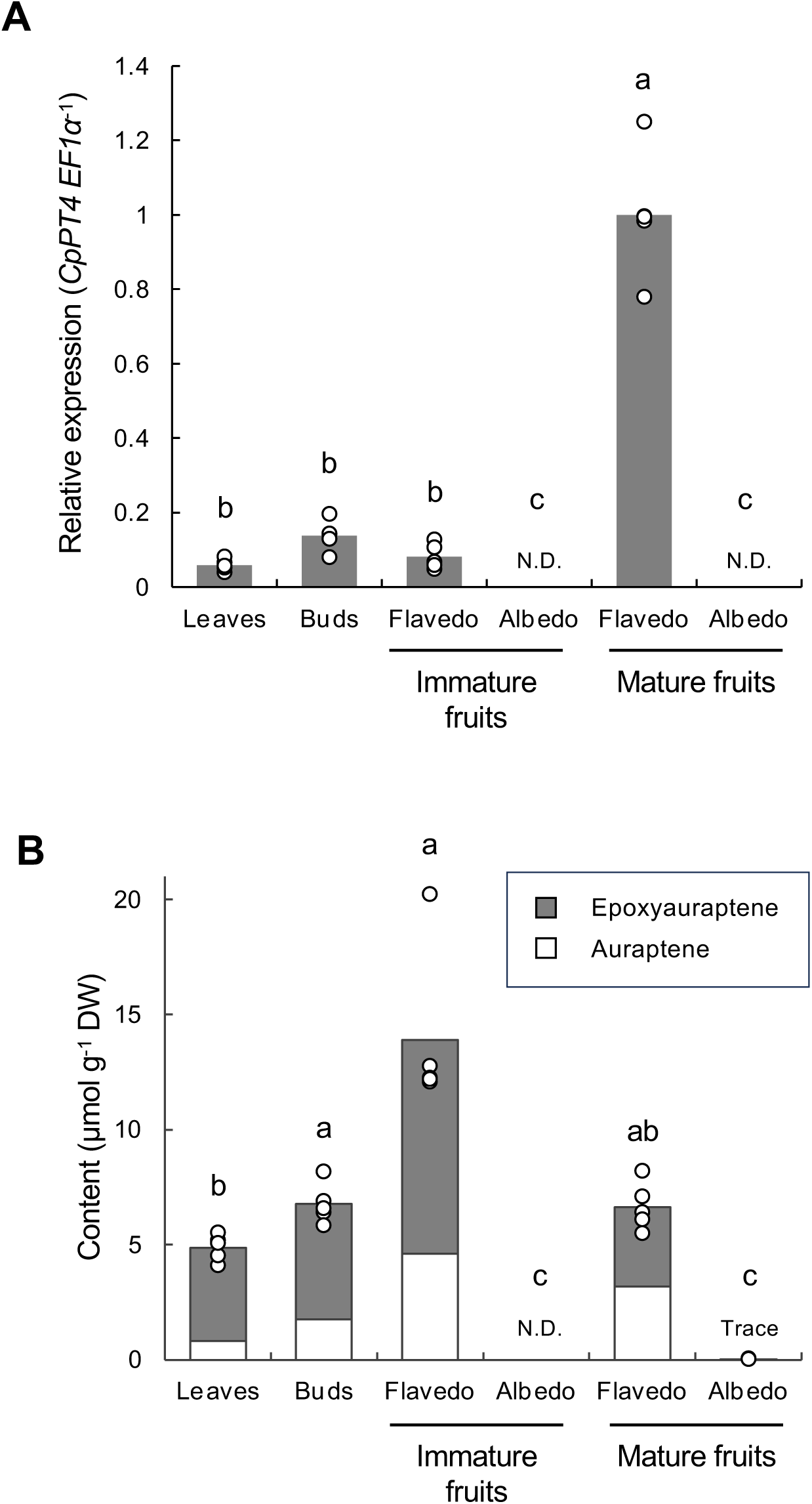
Expression levels of *CpPT4* and accumulation of auraptenes in different grapefruit organs. (A) Relative expression levels of *CpPT4* normalized to *CpEF1α* in grapefruit leaves, buds, immature fruit flavedo, immature fruit albedo, mature fruit flavedo, and mature fruit albedo. Expression levels are shown relative to the average *CpPT4* /*CpEF1α* ratio in mature fruit flavedo. Data represent five biological replicates; bars indicate mean values. (B) Auraptene (white bars) and epoxyauraptene (gray bars) levels in the same grapefruit organs analyzed in (A). Quantification data were obtained from our previous study (Munakata et al., 2021). Value represents the mean of five biological replicates. Significant differences among organ types were evaluated using the Games–Howell test (p < 0.05). Different letters indicate statistically significant differences between the groups. N.D., not detected.

We also assessed the subcellular localization of CpPT4 *in planta* using a heterologous expression system. synthetic GFP (sGFP) was fused to the C terminus of the first 70 amino acids containing the predicted TP of CpPT4 (CpPT4TP-sGFP) and transiently expressed in the epidermal cells of *N. benthamiana* by agroinfiltration. Confocal microscopy of epidermal cells expressing CpPT4TP-sGFP revealed that the chimeric protein localized to chloroplasts (Fig. 6). The same localization pattern was observed for the chimeric protein between the full-length CpPT4 and sGFP (CpPT4-sGFP) (Fig. 6), although the GFP signal was weaker than that of CpPT4TP-sGFP. Our microscopic observations strongly suggest that CpPT4 functions in plastids, where GPP, the preny donor substrate of CpPT4, is supplied via the methylerythritol phosphate (MEP) pathway.

**Fig. 6.**
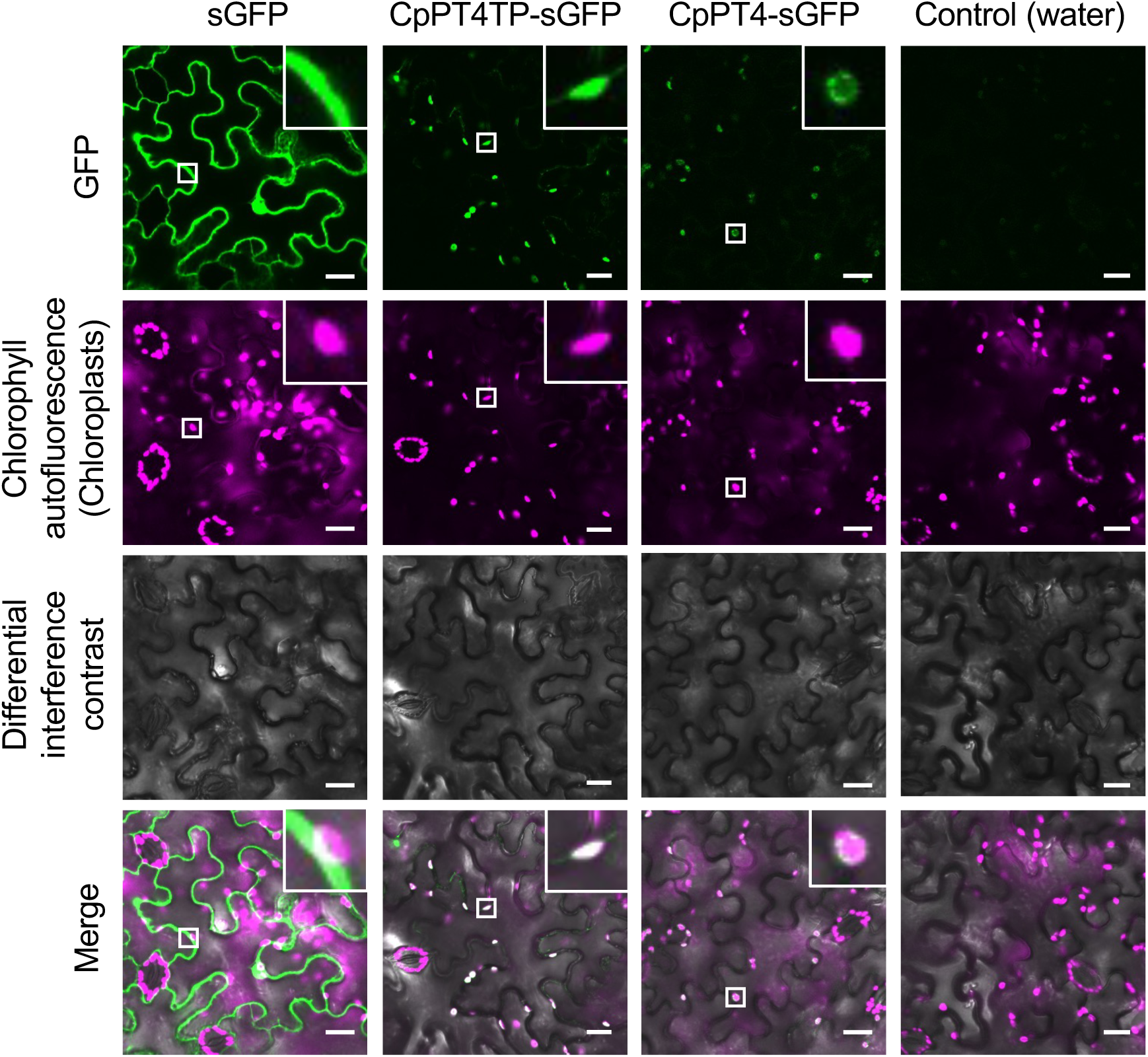
Subcellular localization of CpPT4. CpPT4TP-sGFP and full-length CpPT4-sGFP fusion proteins were transiently expressed in *N. benthamiana* leaves via agroinfiltration. Confocal microscopy was used to image the epidermal cells expressing the fusion proteins. Free sGFP and water infiltrations were used as controls. Chloroplasts were identified based on chlorophyll autofluorescence and are shown in magenta (pseudocolor). The image brightness and contrast were uniformly adjusted across all samples. Inserts are magnified views of the chloroplast regions. Scale bars, 20 µm.

### Polymorphism of the CpPT4 locus in citrus species

To gain insight into the conservation of *CpPT4* orthologs in citrus, a blastn search was performed for the public genomes of various citrus species belonging to the *Citrus*, *Fortunella*, *Poncirus*, and *Atalantia* genera (a primitive citrus group) (Liu et al., 2022), using the *CpPT4* CDS as a query. This analysis identified single loci showing high nucleotide identities with *CpPT4* in all examined genomes (Fig. 7, Table S3, and Fig. S7), and we presumed these sequences to be the orthologs of *CpPT4*. This conservation is consistent with the distribution of auraptenes in these genera (Ogawa et al., 2000; Pailee et al., 2020). However, regarding the quantitative trait, the concentration of auraptenes largely varies among citrus species (Dugrand-Judek et al., 2015), implying interspecies diversity in the functionality of this locus. To confirm this assumption, we conducted an in-depth search for *CpPT4* orthologs with mutations that lead to loss of function. Furthermore, the presence of these mutations was confirmed by PCR-based genotyping of representative commercial citrus species, *i.e.,* pummelo, mandarin, sweet orange, and clementine.

**Fig. 7.**
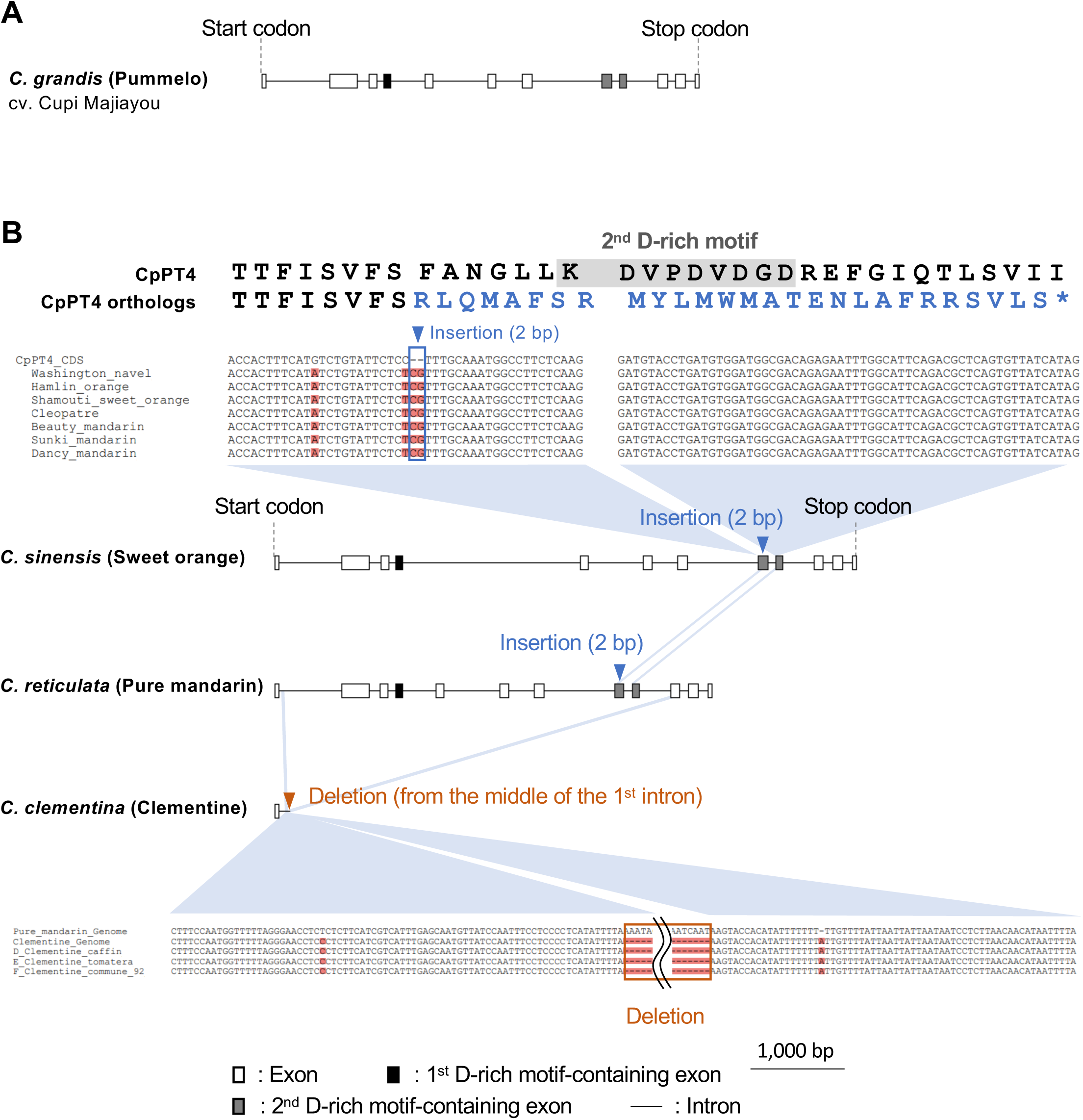
The gene structures of *CpPT4* orthologs in citrus species. (A) Gene structure of the *CpPT4* ortholog in pummelo (cv. Cupi Majiayou). Exons and introns are represented by boxes and lines, respectively, in the schematic. The exons containing the first and second aspartate-rich motifs are highlighted in black and gray, respectively. (B) Gene structures of *CpPT4* orthologs in sweet orange, pure (ancestral) mandarin (cv. Mangshan), and clementine (cv. Clemenules), along with the corresponding genomic sequences, were validated using PCR. Blue and orange triangles indicate insertions and deletions predicted to cause loss of function, respectively. PCR validation confirmed the insertions in four pure mandarin and three sweet orange varieties and the deletion in three clementine varieties. Alignment of genomic sequences from pure mandarins and clementines (Pure_mandarin_Genome and Clementine_Genome) illustrated the exon–intron structure near the deletion site.

First, we analyzed pummelo, an auraptenes-rich species (Dugrand-Judek et al., 2015), for which the genomes of two varieties, Cupi Mjiayou and Wanbaiyou, are available (Liu et al., 2022). In the Cupi Mjiayou genome, we found a sequence that enables the expression of the CDS identical to *CpPT4*, in line with the metabolite profile of pummelo and the relationship that pummelo is a female parent of grapefruit (Wu et al., 2018). In contrast, the Wanbaiyou genome possesses a *CpPT4* ortholog that contains a 2-bp deletion in the 6^th^ exon, which causes an in-frame stop codon in the 9^th^ exon (Fig. S7A). The resulting enzyme is probably not functional because of the lack of the last quarter of its polypeptide, which contains the 2^nd^ aspartate-rich motif crucial to the catalytic function of the UbiA-type PT (Cheng and Li, 2014). This result was unexpected, as Wanbaiyou contains auraptene at a level comparable to that of other pummelo varieties (Ogawa et al., 2000). Thus, we assessed this deletion using genomic PCR and found it in none of the tested 35 pummelo varieties, including our Wanbaiyou individual. The 2-bp deletion leading to loss-of-function of this gene is unlikely to be prevalent in pummelo, and the variation at this site is within Wanbaiyou.

The genomes of pure mandarins (an ancestral species of sweet orange and clementine), sweet orange, and clementine were also investigated as representatives of citrus species that accumulate few auraptenes. Amplicon sequencing showed that the pure mandarin and sweet orange genomes commonly contain a 2 bp insertion before the 2^nd^ aspartate-rich motif. This insertion causes a frameshift that leads to an in-frame stop codon at the 9^th^ exon, and thereby, the gene product is predicted to be dysfunctional as an aromatic PT. The clementine genome has a distinct mutation, with most of its *CpPT4* ortholog sequences lost (Fig. 7). These insertions and deletions were confirmed by genomic PCR of several varieties of sweet orange, commercial mandarin, and clementine (Fig. 7).

Other citrus species were also searched using public genome databases. In a phytochemical investigation of citrus flavedo, *F. hindsii* and *P. trifoliata* produced auraptene at levels similar to those in pummelo and grapefruit varieties, whereas this molecule was undetectable in *C. ichangensis* (Ogawa et al., 2000). In accordance with this result, the *CpPT4* orthologs of *F. hindsii* and *P. trifoliata* do not possess any clear mutations that lead to loss-of-function, and that of *C. ichangensis* has a frame-shift mutation that produces a truncated polypeptide lacking approximately the last quarter of its full length (Fig. S7). However, for some species, we were unable to draw a conclusion about the link between the predicted intactness of the *CpPT4* ortholog and the accumulation level of auraptenes. Citron (*C. medica*) is a progenitor species and the male parent of lime (*C. aurantiifolia*) and lemons (*C. limon*). Although citron accumulates trace amounts of auraptenes (Dugrand-Judek et al., 2015), its *CpPT4* ortholog is likely functional in its primary sequence (Fig. S7).

In summary, our investigation of the *CpPT4* orthologs in diverse citrus species revealed mutations that are possibly related to the undetectable or trace accumulation of auraptenes in mandarin, sweet orange, clementine, and *C. ichangensis*. Further analysis is required to fully explain the molecular mechanisms underlying the diversity in the auraptene content in citrus.

## Discussion

Aromatic *O*-prenylation is a biosynthetic reaction that contributes to the chemo-diversity and biological function of aromatic molecules in plant specialized metabolism (Epifano et al., 2007; Munakata and Yazaki, 2024), whereas most aromatic *O*-PT genes remain unexplored in plants. Regarding auraptene biosynthesis, a study on *Murraya exotica* (Rutaceae) demonstrated that a UbiA-type PT, MePT1, catalyzes the U7OGT reaction to yield auraptene; however, this activity is much less efficient than its main function, U8GT (Li et al., 2022). Moreover, its possible orthologs in lemon (ClPT1) and grapefruit (CpPT3) also show U8GT activity as their main function (Munakata et al., 2021, 2014), suggesting the presence of a genuine aromatic PT gene dedicated to auraptene synthesis in Rutaceae. In this study, we identified *CpPT4*, a UbiA superfamily gene, as a *U7OGT*, completing the final step in the auraptene biosynthetic pathway.

### Molecular evolution of the UbiA superfamily proteins to aromatic O-PTs in Rutaceae

The integration of CpPT4 into the phylogenetic tree sheds light on the evolutionary relationship between aromatic *O*-PTs and *C*-PTs in Rutaceae. Our previous report suggested that aromatic *O*-PTs arose independently in Rutaceae and Apiaceae (Munakata et al., 2021). Consistent with this evolutionary process, CpPT4 was included in the Rutaceae clade of the phylogenetic tree. Unexpectedly, CpPT4 was located next to the *C*-PT subclade of U8GTs (Cl-PT1, CpPT3, and MePT1) and distant from the *O-*PT subclade of bergaptol 5,8-*O*-GTs (CpPT1 and CmiPT1a/b). This *O*-PT subclade is adjacent to MePT2, another *C*-PT (Li et al., 2022). This mixed distribution between *C*-PTs and *O*-PTs indicates the possibility that functional diversification between aromatic *C*-PTs and *O*-PTs occurred multiple times within Rutaceae. Many specialized UbiA-type PT genes remain uncharacterized in the Rutaceae genomes. Elucidating their functions will help capture the molecular evolution process of the UbiA superfamily in this plant taxon at a higher resolution.

An open question related to the acquisition of aromatic *O*-PTs is how aromatic *O*-prenylation is more beneficial to the environmental adaptation of rutaceous plants than aromatic *C*-prenylation. Aromatic *O*-prenylation significantly increases the hydrophobicity of aromatics by masking hydroxyl moieties directly with hydrophobic prenyl chains, which may lead to defensive activities unique to *O*-prenylated derivatives. In addition, this moiety may be relevant to the accumulation of aromatics. In citrus, *O*-prenylated coumarins, including auraptene, mainly accumulate in oil cavities, which are extracellular spaces primarily filled with hydrophobic volatile terpenes (Voo et al., 2012). In general, specialized metabolites stored in oil cavities do not possess hydrophilic moieties. For example, major terpenes, such as limonene and myrcene, are hydrocarbons, and the hydroxyl moieties of flavonoids are fully methoxylated. This suggests that the hydrophobicity conferred by *O*-prenylation may facilitate the accumulation of certain aromatic molecules in the hydrophobic extracellular space. Some specialized metabolites are toxic to the producing cells, and more specifically, citrus oil cavities accumulate furanocoumarins, which are toxic to plant cells (Murray et al., 1982), in their *O*-prenylated and/or *O-*methylated forms. The extracellular accumulation strategy in citrus enables the sequestration of toxic specialized metabolites from organelles, thereby avoiding the risk of self-intoxication.

### Unique biochemical properties of CpPT4 and its application to synthetic biology

A common feature of plant UbiA PTs is their ability to catalyze aromatic prenylation with high reaction specificity for both substrate recognition and prenylation position (Munakata and Yazaki, 2024). Similarly, recombinant CpPT4 exhibited high substrate and product specificities toward U7OGT activity, which is advantageous in metabolic engineering, such as auraptene production in microorganisms by synthetic biology. Microbial strains with high productivity of CpPT4 substrates, such as umbelliferone (Wang et al., 2023) or GPP (Zhao et al., 2016), have been established. Moreover, modifications of membrane-bound UbiA-type PTs for functional expression have been reported in different microbial host species, including bacteria (Bamba et al., 2024; Carruthers et al., 2025; Liu et al., 2025; Munakata et al., 2019). The integration of these engineering approaches would enable the establishment of a *de novo* microbial production system for auraptene. Although auraptene can be obtained by extraction from citrus flavedo, many auraptene-rich citrus species also accumulate furanocoumarins that are toxic to humans in this tissue (Dugrand-Judek et al., 2015). Synthetic biology-based systems using CpPT4 will enable intrinsic auraptene production without the risk of toxic molecule contamination.

An important finding of our biochemical analysis was the detection of aromatic *N*-prenylation activity of the UbiA-type PT. In most aromatic prenylations by this enzyme family, the prenyl moiety is directly transferred to the electron-donating hydroxyl moiety on the benzene ring (*O*-prenylation) or the carbon at the *ortho*-position to this hydroxyl moiety (*C*-prenylation) (Munakata and Yazaki, 2024). The acceptance of 7-aminocoumarin by CpPT4 indicates that the amino moiety functions as an electron-donating group, which is the first example in the UbiA superfamily. Recently, the catalytic mechanisms underlying the interaction with aromatic substrates have been intensively studied in plant UbiA-type PTs (Ernst et al., 2024; Han et al., 2025). These advances will enable the engineering of UbiA-type PTs that efficiently transfer prenyl chains directly to amino groups or their *ortho*-positions on aromatic substrates. Our biochemical data opened a way to reach the untapped chemical space of prenylated aromatics using UbiA-type PTs. Considering the bioactivities of prenylated aromatics, their analogs, including 7-geranylaminocoumarin synthesized in this study, would be promising targets for screening new bioactive molecules.

### Application of CpPT4 to citrus breeding

In agriculture, specialized metabolites such as polymethoxyflavones, furanocoumarins, volatile terpenes, carotenoids, and limonoids have a substantial impact on the quality of citrus fruits and related products (Omura and Shimada, 2016). Thus, genomic loci related to the contents of these metabolites have been intensively investigated (Matsumoto et al., 2022; Munakata et al., 2021; Yu et al., 2017), and polymorphisms that potentially serve as practical molecular markers have been identified. Regarding the contribution to citrus breeding, our genotyping analysis found polymorphisms in *CpPT4* orthologs that are possibly relevant to the auraptene content in different commercial species. In particular, mandarin, sweet orange, and clementine have dysfunctional mutations in this gene, in line with their low auraptene accumulation. These mutations may serve as molecular markers to eliminate progenies with low auraptene content in fruits. Similarly, the intact *CpPT4* sequences of auraptene-rich species, such as pummelo, may be used as indicators of high auraptene content.

In contrast, we also found complex auraptene metabolism in citrus. In the expression analysis, *CpPT4* was highly expressed in flavedo and much less in albedo in immature and mature fruits, consistent with the accumulation pattern of auraptenes (Munakata et al., 2021). However, although mature flavedo accumulates less auraptenes than immature flavedo, the relationship between the expression levels of this gene is the opposite in these organs. One possible explanation for this mismatch lies in substrate availability; the supply of umbelliferone or GPP, the substrates for CpPT4, would be the rate-limiting step in the biosynthesis of auraptenes in mature flavedo. Moreover, umbelliferone and GPP are intermediates of other coumarin molecules and monoterpenes, respectively, in citrus flavedo, indicating that competition between substrates among multiple pathways should also be considered as well (Voo et al., 2012). These factors may explain why citron (*C. medica*) accumulates undetectable levels of auraptenes, but its *CpPT4* ortholog sequence is seemingly intact (Dugrand-Judek et al., 2015). To find a rational approach to create citrus varieties with higher auraptene contents than existing varieties, it is necessary to consider the related metabolic network, which varies by the developmental stage of the fruit and by citrus varieties/species.

## Conclusion

This study provides a new example of the UbiA superfamily’s contribution to the biosynthesis of plant *O*-prenylated aromatics, which include various pharmacologically active molecules. We also demonstrated that the amino moiety can serve as an electron-donating group in aromatic prenylation by this enzyme family, potentially expanding the chemical diversity of prenylated aromatics. CpPT4 can be useful for the synthetic biology-based production of auraptene and its analogs. In citrus breeding, the polymorphism of this gene among citrus species would lead to the development of a molecular marker for the auraptene content.

## Materials and methods

### Plant materials and reagents

Immature fruits of Ruby grapefruit (*C.* × *paradisi* cv. Ruby) were collected from the Jardins Botaniques de Nancy (Nancy, France). Marsh grapefruit (*C.* × *paradisi* cv. Marsh) organs were collected from Regional Revitalization and Agricultural Research Institute of Kindai University. Leaves of various citrus species were obtained from Corsica, Regional Revitalization and Agricultural Research Institute of Kindai University, and the Research Center of Genetic Resources, National Agriculture and Food Research Organization (NARO), Japan (NARO Genebank: https://www.gene.affrc.go.jp/index_en.php) (Table S4). These plant samples were frozen in liquid nitrogen and stored at -80 ℃. DMAPP and GPP used in the preliminary tests were kindly provided by Dr. Hirobumi Yamamoto (Toyo University), Dr. Tomohisa Kuzuyama (The University of Tokyo), and Dr. Takashi Kawasaki (Kyoto University). DMAPP and GPP in the other experiments and other prenyl diphosphates were purchased from Sigma-Aldrich (St. Louis, MO, US). Limettin was kindly provided by Dr. Frederic Bourgaud (Université de Lorraine, Nancy, France). The other aromatic molecules were purchased from Alfa Aesar (Thermo Fisher Scientific, Waltham, US), BioDuro (Irvine, US), Extrasynthase (Lyon, France), FUJIFILM Wako Pure Chemicals (Osaka, Japan), Herboreal (Edinburgh, UK), Indofine Chemical Company (Hillsborough, NJ, US), Sigma-Aldrich, Tokyo Chemical Industry (Tokyo, Japan), and Toronto Research Chemicals (Vaughan, Canada).

### Isolation of CpPT4 gene from grapefruit and construction of its plant expression plasmid for in vitro characterization

Frozen immature flavedo of the Ruby grapefruit cultivar was ground into powder using a mortar and pestle. Total RNA was extracted from the powder using the E.Z.N.A Plant RNA Kit (Omega Bio-tek, Norcross, USA) and DNaseI (Sigma-Aldrich, St. Louis, USA) and subjected to first-strand cDNA synthesis using the SuperScript IV First-Strand Synthesis System (Thermo Fisher Scientific). The cDNA pool was used as a template in PCR using PrimeSTAR Max DNA Polymerase (Takara Bio, Kusatsu, Japan) and the primer pair CpPT4_5’UTR_Fw and CpPT4_3’UTR_Rv (Table S5). The amplicon was inserted into the vector pCR8/GW/TOPO by TOPO reaction-based TA cloning and sequenced. The full-length CDS of CpPT4 was amplified by PCR using PrimeSTAR Max DNA Polymerase (Takara Bio) and the primer pair CpPT4_TOPO_Fw and CpPT4_TOPO_Rv (Table S5). The resulting amplicon was inserted into the vector pENTR/D-TOPO (Thermo Fisher Scientific) by directional TOPO reaction and finally into the vector pGWB502 by LR recombination with the Gateway LR Clonase II Enzyme mix (Thermo Fisher Scientific) to construct a plasmid containing *P35S-CpPT4-TNos*. The sequence identical to *CpPT4* was isolated from the immature flavedo of the grapefruit cultivar Marsh using essentially the same procedure, except for total RNA extraction with the RNeasy Plant Mini kit (Qiagen, Hilden, Germany) and cloning with pGEM-T Easy Vector Systems (Promega, Madison, USA).

### Enzymatic characterization of CpPT4

To express CpPT4 in *N. benthamiana*, the plasmid pGWB502-*CpPT4* was introduced into *Agrobacterium tumefaciens* strain LBA4404. This strain was infiltrated into *N. benthamiana* leaves along with the *A. tumefaciens* strain C58C1 carrying pBIN61-*P19* (Voinnet et al., 2003). Microsomal fractions were extracted from infiltrated leaves following the protocol of Karamat et al. (2014), suspended in 100 mM Tris-HCl buffer (pH 8.0), and stored at −80 °C. For *in vitro* enzymatic assays, the microsomes were diluted in the same buffer and used immediately. The standard reaction mixture (200 μL total volume) included 200 μM of each prenyl acceptor and donor substrate, 10 mM MgCl₂, and the microsomal preparation of CpPT4 in 100 mM Tris-HCl (pH 8.0) supplemented with 1 mM dithiothreitol. reactions were incubated at 28 °C for 2 hours. As a negative control, we used microsomes obtained from *N. benthamiana* leaves expressing a fusion protein of the transit peptide of CpPT1 (N-terminal 70 amino acids) and sGFP (CpPT1TP-sGFP) (Munakata et al., 2021).

### Extraction of reaction products

The substrates and products were extracted using 300 μl of ethyl acetate. Prior to extraction, 100 μl of water was added to adjust the volume for easy collection of the ethyl acetate fraction in our experimental setup. The mixtures were vortexed for 5 min and centrifuged at 4,400 × *g* for 5 min, and 200 μl of the ethyl acetate fraction ( upper phase) was collected. The extract was vacuum evaporated to dryness and dissolved in 100 μl of methanol by vortexing for 5 min. After centrifugation for 5 min at 20,400 × *g*, the supernatant was subjected to LC/MS analysis.

### LC/MS analysis of extracts of CpPT4 reaction mixtures

The enzyme products were analyzed using a NEXERA UHPLC system (Shimadzu, Kyoto, Japan) equipped with a photodiode array detector (PDA, SPD-M40, Shimadzu). Separation was performed on a C18 reverse-phase column (ACQUITY UPLC BEH C18 1.7 μm, 2.1 × 50 mm, Waters) using a linear gradient of solvent A (water containing 0.1 (v/v) formic acid) and solvent B (acetonitrile containing 0.1% (v/v) formic acid). The gradient program was as follows: 20-95% solvent B over 2.50 min; maintained at 95% B for 1.00 min, followed by re-equilibration to the initial conditions. The flow rate was 1.0 ml min^-1^, and the column temperature was maintained at 40°C. The absorbance was monitored in the range of 250– 370 nm.

For LC/MS^2^ analysis, a Vanquish UHPLC System (Thermo Fisher Scientific) coupled to a Q Exactive Focus mass spectrometer (Thermo Fisher Scientific) was used. Chromatographic separation was performed on a Shim-pack Velox Biphenyl column (1.8 μm, 2.1 × 100 mm^2^, Shimadzu) using the same solvent system as above. The gradient was: 5-95% solvent B over 20.00 min, held at 95% B for 5.00 min, and returned to the initial conditions. The flow rate was 0.3 ml min^-1^ at 40°C. Pierce LTQ Velos ESI Positive Ion Calibration Solution (Thermo Fisher Scientific) was used as the calibration standard. The additional LC/MS^2^ parameters are listed in Table S6.

### RT-qPCR

We used cDNA samples synthesized from total RNA of different citrus organs, as described in our previous study (Munakata et al., 2021). In this study, each citrus sample was ground to a powder using a mortar and pestle, and the resulting powder pool was used for both total RNA extraction for RT-qPCR and coumarin quantification (Fig. 5B).

RT-qPCR was performed using a CFX96 Touch Deep Well system (Bio Rad, Hercules, CA, USA) with Thunderbird SYBR qPCR Mix (Toyobo), following the manufacturer’s protocol. Each 25 μl reaction contained cDNA template, 12.5 μl of SYBR qPCR Mix, and either 5.0 pmol of each *CpPT4*-specific primer (CpPT4_qRTPCR_Fw and CpPT4_qRTPCR_Rv) or 7.5 pmol of *CpEF1α* reference gene primer (CpEF1α_Fw and CpEF1α_Rv) (Table S5). The thermal cycling conditions were as follows: initial denaturation at 95°C for 2 min, followed by 35 cycles of denaturation at 95°C for 15 s and annealing/extension at 62°C for 30 s.

### Construction of plasmids for expression of GFP-fusion proteins and microscopic observation

For subcellular localization analysis, two *CpPT4* constructs were generated: *CpPT4TP,* which encodes the N-terminal 70 amino acids of CpPT4, and *CpPT4 (-stop),* which encodes the full-length coding sequence without the stop codon. These fragments were PCR-amplified using PrimeSTAR Max DNA Polymerase (Takara Bio) and the primer pairs CpPT4_TOPO_Fw/CpPT4_TP210_Rv and CpPT4_TOPO_Fw/CpPT4_woStop_Rv (Table S5).

The PCR products were first cloned into the pENTR/D-TOPO vector, followed by LR recombination into the pGWB505 destination vector (Nakagawa et al., 2007), resulting in constructs expressing *P35S-CpPT4TP-sGFP-Tnos* and *P35S-CpPT4(-stop)-sGFP-Tnos*. GFP-fusion proteins were transiently expressed in *N. benthamiana* leaves via agroinfiltration. A construct expressing free sGFP (*P35S-sGFP-Tnos* in pHKN29 (Kumagai and Kouchi, 2003)) was used as the control.

Confocal laser scanning microscopy was performed using an FV3000 microscope (Olympus, Tokyo, Japan) equipped with a UPLFLN 40× objective lens. GFP fluorescence was excited at 488 nm and detected at 500–540 nm, whereas chloroplast autofluorescence was excited at 640 nm and detected at 650–750 nm. Differential interference contrast (DIC) images were simultaneously acquired. All the imaging parameters were kept constant across all samples.

### Genotyping of the CpPT4 locus in various citrus species

Frozen leaf tissues from multiple citrus species were ground into fine powder using mortar and pestle. Genomic DNA was extracted using either the E.Z.N.A. Plant DNA Kit (Omega Bio-tek) or DNeasy Plant Pro Kit (Qiagen). Partial genomic sequences of *CpPT4* orthologs were amplified by PCR using extracted DNA, PrimeSTAR Max DNA Polymerase (Takara Bio), and species-specific primer pairs (Csinensis_Fw/Csinensis_Rv for *C. sinensis* and *C. reticulata*, Cclementina_Fw/Cclementina_Rv for *C. clementina*, and Cgrandis_Fw/Cgrandis_Rv for *C. grandis*) (Table S5).

PCR products were separated by agarose gel electrophoresis and purified from gel slices using the Wizard SV Gel and PCR Clean-Up System (Promega) prior to sequencing. The number of biological replicates (trees), technical replicates (leaves), and sequenced amplicons differed among the citrus varieties (Table S4).

### In silico analysis

PT sequences were obtained from the NCBI (https://www.ncbi.nlm.nih.gov/), Phytozome (https://phytozome.jgi.doe.gov/pz/portal.html#), Citrus sinensis Annotation Project (currently updated as Citrus Pan-genome to Breeding Database, http://citrus.hzau.edu.cn/) (Liu et al., 2022), and OneKP (https://www.onekp.com/) databases. The genome datasets used for the ortholog identification of *CpPT4* are listed in Table S7. Prediction of transit peptides was performed using TargetP 2.0 (Almagro Armenteros et al., 2019) and DeepLoc (Thumuluri et al., 2022), whereas multiple transmembrane domain predictions were conducted using DeepTMHMM (Hallgren et al., 2022). Local blast searches against the grapefruit flavedo transcriptome were conducted using BioEdit (http://www.mbio.ncsu.edu/BioEdit/bioedit.htmlhttp://www.mbio.ncsu.edu/BioEdit/bioedit.html), which was also used to calculate amino acid identities among PTs. For phylogenetic analysis, PT sequences were aligned using MUSCLE and MEGA 11 (http://www.megasoftware.net/). A maximum-likelihood phylogenetic tree was constructed using IQ-TREE v2.2.2.6 (Minh et al., 2020), using the JTT+F+R6 model selected via ModelFinder (Kalyaanamoorthy et al., 2017). Branch support was assessed using UFBoot (Hoang et al., 2018). During BLAST-based gene or protein mining, the terminal boundaries of the hit regions were manually adjusted to prevent the ambiguous mapping of query bases to multiple sites.

### Statistics and reproducibility

Relative apparent *K*_m_ values were determined by non-linear fitting using Sigmaplot version 14.5. Substrate specificity, organ-specific expression of *CpPT4*, and auraptene content were statistically analyzed using the Tukey and Games-Howell tests, as appropriate, with R software version 4.2.1 (R Core Team, 2022).

For organ-specific analyses, five buds, five leaves, flavedo, and albedo tissues from five immature fruits and flavedo and albedo tissues from five mature fruits were used. *In vitro* enzymatic activity assays were performed in three independent replicates for each sample.

## Data availability

The nucleotide sequence of *CpPT4* is available in NCBI under the accession number LC887959.

## Acknowledgments

We thank Dr. Yann Froelicher (International Center for Agronomic Research and Development, San Giuliano), Dr. Patrick Ollitrault (Centro de Protección Vegetal y Biotecnología, Valencia), the Genebank of NARO, and Jardins Botaniques de Nancy for providing citrus samples. We also thank Dr. Tomohisa Kuzuyama (The University of Tokyo) and Dr. Takashi Kawasaki (Niigata University) for GPP, Dr. Hirobumi Yamamoto (Toyo University) for DMAPP, Dr. Frédéric Bourgaud (Université de Lorraine) for coumarin derivatives, Dr. David Baulcombe for the pBIN61-P19 plasmid, Dr. Tsuyoshi Nakagawa (Shimane University) for pGWB vectors, and Dr. Hiroshi Kouchi (International Christian University) for the pHKN29 plasmid. We are grateful to Ms. Kaori Kanazawa (Kyoto University), Mr. Yoko Kamata (Kyoto University), Mr. Clément Charles (Université de Lorraine), Ms. Auriane Drack (Université de Lorraine), Ms. Chloé Perrier (Université de Lorraine), Mr. Maxime Guffroy (Université de Lorraine), and Ms. Rose Bulteau (Université de Lorraine) for technical assistance. Microscopic analysis was performed with the technical support of Dr. Takuji Ichino (Kobe Pharmaceutical University, Kobe, Japan). The Development and Assessment of Sustainable Humanosphere system of the Research Institute for Sustainable Humanosphere (Kyoto University) provided the plant growth environment.

## Author Contributions

R.M., A.H., K.Y. conceived research. R.M., A.H., A.S., K.Y. supervised research. S.M., R.M., M.R., A.O., N.M. performed the experiments. T.M. maintained and provided research materials. S.M., R.M., K.Y. wrote the manuscript with contribution of all the authors.

## Supplemental Tables and Figures

**Table S1. PT sequences used for in silico analysis.**

UbiA PTs are involved in primary and specialized metabolism (VTE2-1-, VTE2-2-, and PPT-related PTs) in plants.

**Table S2. *In vitro* functional screening of CpPT4.**

The enzymatic function of recombinant CpPT4 was screened using simple (bicyclic) coumarins, furanocoumarins, and other aromatic molecules as prenyl acceptors and DMAPP, GPP, FPP, and GGPP as prenyl donors (n = 1). Substrate pairs resulting in enzymatic products are marked with pluses. N.D., not detected. N.T., not tested.

**Table S3. Summary of BLAST analysis for identifying orthologs in citrus genomes.**

BLASTN searches were performed using *CpPT4* CDS against 10 citrus genomes with -outfmt 6, max_target_seqs=5000, and max_hsps=5000. Multiple high-scoring segment pairs for the same query– subject pair were merged on the query side to calculate coverage, and subject intervals within 10 kb were considered to belong to the same locus. For each query–hit locus pair, the maximum identity (Max Identity), total bitscore, and best e-value were calculated. Top hits that showed the highest Bitscore and lowest E-values were designated as orthologs, except for *C. clementina* whose top hit showed a much lower bioscore and a much higher E-value than the top hits of the other tested species. For this species, the schaffold1_locus1 was assigned as an ortholog because its full hit sequence showed 100% identity with the corresponding region of CpPT4. Moreover, a large deletion was detected directly downstream of this locus, which greatly affected the bitscore and E-value of this locus.

**Table S4. Polymorphism in CpPT4 ortholog locus of C. sinensis, C. clementina, C. reticulata, and C. grandis.**

Varieties with mutations leading to loss of function confirmed by PCR sequencing are marked with a plus and varieties without the mutations are marked with a minus. The number of iterations of tree, leaf, and PCR amplicons are noted.

**Table S5. Primers used in this work.**

**Table S6. LC/MS^2^ conditions for analysis.**

**Table S7. Genome assemblies used for *CpPT4* ortholog identification.**

**Fig. S1. Multiple alignment of the pummelo, c22985_g1_i1, and *CpPT4*.**

The second aspartate-rich motif is highlighted by a blue square. The upper ruler indicates the base number from the start codon on the pummelo (cv. Wanbaiyou).

**Fig. S2*. In silico* analysis of the CpPT4 polypeptide.**

**(A)** Multiple transmembrane regions of CpPT4 predicted by DeepTMHMM. **(B)** Probability of cleavage site of CpPT4 predicted by TargetP-2.0. **(C)** Transit peptide of CpPT4 predicted by DeepLoc. **(D)** MUCSLE multiple alignment of CpPT4 and Rutaceae PTs. The first and second aspartate-rich motifs are highlighted by the boxes. **(E)** Amino acid sequence identity among the PTs of Rutaceae.

**Fig. S3. The chemical structure of prenyl acceptor substrates tested in functional screening of CpPT4.**

Substrates that accept prenyl moieties are highlighted in orange. The number of carbons in umbelliferone (compound 1) is shown in grey.

**Fig. S4. Characterization of the U7OGT activity of CpPT4.**

**(A)** Negative control assays for U7OGT activity. From left to right: incubations performed in the presence of EDTA, with microsomes after heat denaturation (95 °C for 25 min), in the absence of MgCl_2_, umbelliferone, GPP, or microsomes, and the full assay. U7OGT activities are shown relative to the mean U7OGT activity in the full assay. The bars represent the means of three independent experiments (n = 3). N.D., not detected. **(B)** Divalent cation dependence of U7OGT activity. Incubation was performed in the presence of MgCl_2_, MnCl_2_, CaCl_2_, CoCl_2_, ZnCl_2_, or water. U7OGT activities are shown relative to the mean U7OGT activity with MgCl_2_. The bars represent the means of three independent experiments (n = 3). N.D., not detected. **(C)** pH dependency. U7OGT activity was measured at pH 6.0 to 9.0 using KPi buffer (pH 6.0 to 7.5) and Tris-HCl buffer (pH 7.5 to 9.0). The results of three independent experiments (n = 3) are shown relative to the mean U7OGT activity at pH 8.0. The lines are drawn with the means. **(D)** Temperature dependency of the data. Incubations were performed at temperatures ranging from 10 to 40 °C. The results of three independent experiments are shown relative to the mean U7OGT activity at 28 °C. The line is drawn with the means. **(E)** Kinetic analysis. The apparent *K*_m_ values for GPP (left) and umbelliferone (right) were calculated using the nonlinear least squares method. The concentrations of substrates ranged from 2–250 µM for GPP and 2–250 µM for umbelliferone in the presence of 250 µM co-substrate. For each condition, three independent experiments were performed (n = 3).

**Fig. S5. Enzymatic reactions catalyzed by recombinant CpPT4.**

**(A-H)** *O/N-*GT activities of CpPT4 toward coumarin derivatives, excluding umbelliferone. **(A, C, E, G)** UV chromatograms of reaction mixtures containing CpPT4 and individual coumarins— **(A)** 5,7-dihydroxycoumarin (no. 3), **(C)** esculetin (no. 4), **(E)** scopoletin (no. 6), and **(G)** 7-aminocoumarin (no. 11) —in the presence of GPP as the prenyl donor. CpPT1TP-sGFP served as the negative control, and its chromatograms are shown at scales comparable to those of the corresponding CpPT4 reactions. **(B, D, F, H)** MS and MS^2^ spectra of the products formed in **(A, C, E, G)**, respectively, acquired in the positive ion mode. Based on the MS^2^ fragmentation patterns and substrate specificity of CpPT4 (Table S2), the products of 5,7-dihydroxycoumarin (no. 3), esculetin (no. 4), and scopoletin (no. 6) were predicted to be 7-*O*-geranylated.

**Fig. S6. Grapefruit organs used in this study.**

Leaves, buds, and immature and mature fruits of grapefruit (cv. Marsh) were harvested from the Regional Revitalization and Agricultural Research Institute farm of Kindai University. The outer peel flavedo and inner white peel albedo were collected from each fruit.

**Fig. S7. The nucleotide sequences of *CpPT4* orthologs in *Citrus*.**

**(A)** MAFFT multiple alignment of *CpPT4* CDS and its orthologs from pummelo (cv. Cupi Majiyaou and Wanbaiyou), sweet orange, pure mandarin, and clementine, respectively. Mismatches with *CpPT4* are highlighted in red. The deletion of the *CpPT4* ortholog in Wanbaiyou is shown by the red box, and the insertion in sweet orange and pure mandarin is shown by the blue box. **(B)** Gene structure model of *CpPT4* ortholog in pummelo based on (A). The deletion of the *CpPT4* ortholog was confirmed by PCR in one pummelo variety, Pomplemousse indien 1 (Pomplemousse_indien_1_PCR_del), which had another sequence without the deletion (Pomplemousse_indien_1_PCR_wodel). Alignment of the related pummelo varieties Cupi Majiayou and Wanbaiyou genomic sequences (Cupi_Majiayou_Genome and Wanbaiyou_Genome) indicated an exon-intron structure proximate to the deletion. **(C)** MAFFT multiple alignment of *CpPT4* CDS and its orthologs from *Atalantia buxifolia*, *Fortunella hindsii*, *Poncirus trifoliata*, *C. ichangensis*, and *C. medica*. Mismatches with *CpPT4* are highlighted in red. The deletion of the *CpPT4* ortholog in *C. ichangensis* is shown in the red box. **(D)** Gene structures of *CpPT4* orthologs of *Atalantia buxifolia*, *Fortunella hindsii, Poncirus trifoliata, C. ichangensis* and *C. medica* (Citron) based on their genome sequences.

## References

1. Adams, M., Ettl, S., Kunert, O., Wube, A., Haslinger, E., Bucar, F., et al. (2006) Antimycobacterial activity of geranylated furocoumarins from *Tetradium daniellii*. Planta Med. 72: 1132–1135.

2. Almagro Armenteros, J.J., Salvatore, M., Emanuelsson, O., Winther, O., von Heijne, G., Elofsson, A., et al. (2019) Detecting sequence signals in targeting peptides using deep learning. Life Sci Alliance. 2: e201900429.

3. Bamba, T., Munakata, R., Ushiro, Y., Kumokita, R., Tanaka, S., Hori, Y., et al. (2024) De novo production of the bioactive phenylpropanoid artepillin C using membrane-bound prenyltransferase in *Komagataella phaffii*. ACS Synth Biol.

4. Carruthers, D.N., Donnell, I., Sundstrom, E., Keasling, J.D., and Lee, T.S. (2025) Prenol production in a microbial host via the “repass” pathways. Metab Eng. 88: 261–274.

5. Cheng, W., and Li, W. (2014) Structural insights into ubiquinone biosynthesis in membranes. Science. 343: 878–881.

6. Chun, J., Lee, J., Ye, L., Exler, J., and Eitenmiller, R.R. (2006) Tocopherol and tocotrienol contents of raw and processed fruits and vegetables in the United States diet. J Food Compos Anal. 19: 196–204.

7. Dugrand-Judek, A., Olry, A., Hehn, A., Costantino, G., Ollitrault, P., Froelicher, Y., et al. (2015) The distribution of coumarins and furanocoumarins in *Citrus* species closely matches *Citrus* phylogeny and reflects the organization of biosynthetic pathways. PLoS ONE. 10: e0142757.

8. Epifano, F., Genovese, S., Menghini, L., and Curini, M. (2007) Chemistry and pharmacology of oxyprenylated secondary plant metabolites. Phytochemistry. 68: 939–953.

9. Ernst, L., Lyu, H., Liu, P., Paetz, C., Sayed, H.M.B., Meents, T., et al. (2024) Regiodivergent biosynthesis of bridged bicyclononanes. Nat Commun. 15: 4525.

10. Fukuda N., Tamai Y., Furukawa Y., Amakura Y., Okuyama S., Yoshimura M., et al. (2019). JP2019033681A.

11. Gravot, A., Larbat, R., Hehn, A., Lièvre, K., Gontier, E., Goergen, J.-L., et al. (2004) Cinnamic acid 4-hydroxylase mechanism-based inactivation by psoralen derivatives: cloning and characterization of a C4H from a psoralen producing plant—*Ruta graveolens*—exhibiting low sensitivity to psoralen inactivation. Arch Biochem Biophys. 422: 71–80.

12. Hallgren, J., Tsirigos, K.D., Pedersen, M.D., Armenteros, J.J.A., Marcatili, P., Nielsen, H., et al. (2022).

13. Hamerski, D., Schmitt, D., and Matern, U. (1990) Induction of two prenyltransferases for the accumulation of coumarin phytoalexins in elicitor-treated Ammi majus cell suspension cultures. Phytochemistry. 29: 1131–1135.

14. Han, J., Munakata, R., Takahashi, H., Koeduka, T., Kubota, M., Moriyoshi, E., et al. (2025) Catalytic mechanism underlying the regiospecificity of coumarin-substrate transmembrane prenyltransferases in Apiaceae. Plant Cell Physiol. 66: 1–14.

15. Hanley, M.J., Cancalon, P., Widmer, W.W., and Greenblatt, D.J. (2011) The effect of grapefruit juice on drug disposition. Expert Opin Drug Metab Toxicol. 7: 267–286.

16. Hendrich, A.B., Malon, R., Pola, A., Shirataki, Y., Motohashi, N., and Michalak, K. (2002) Differential interaction of Sophora isoflavonoids with lipid bilayers. Eur J Pharm Sci. 16: 201–208.

17. Hoang, D.T., Chernomor, O., von Haeseler, A., Minh, B.Q., and Vinh, L.S. (2018) UFBoot2: Improving the ultrafast bootstrap approximation. Mol Biol Evol. 35: 518–522.

18. Igase, M., Okada, Y., Ochi, M., Igase, K., Ochi, H., Okuyama, S., et al. (2018) . J Prev Alzheimers Dis.

19. Ito, C., Mizuno, T., Matsuoka, M., Kimura, Y., Sato, K., Kajiura, I., et al. (1988) A new flavonoid and other new components from *Citrus* plants. Chem Pharm Bull. 36: 3292–3295.

20. Kalyaanamoorthy, S., Minh, B.Q., Wong, T.K.F., von Haeseler, A., and Jermiin, L.S. (2017) ModelFinder: fast model selection for accurate phylogenetic estimates. Nat Methods. 14: 587–589.

21. Karamat, F., Olry, A., Munakata, R., Koeduka, T., Sugiyama, A., Paris, C., et al. (2014) A coumarin-specific prenyltransferase catalyzes the crucial biosynthetic reaction for furanocoumarin formation in parsley. Plant J. 77: 627–638.

22. Komatsu, S., Tanaka, S., Ozawa, S., Kubo, R., Ono, Y., and Matsuda, M. (1930) Biochemical studies on natsumikan (Natsumikan no seikagakuteki kenkyu). J Chem Soc Jpn. 51: 478–498.

23. Kumagai, H., and Kouchi, H. (2003) Gene silencing by expression of hairpin RNA in *Lotus japonicus* roots and root nodules. Mol Plant-Microbe Interactions. 16: 663–668.

24. Li, N., Liu, X., Zhang, M., Zhang, Z., Zhang, B., Wang, X., et al. (2022) Characterization of a coumarin *C*-/*O*-prenyltransferase and a quinolone *C*-prenyltransferase from *Murraya exotica*. Org Biomol Chem. 20: 5535–5542.

25. Liu, Hanmingzi, Wang, X., Liu, S., Huang, Y., Guo, Y.-X., Xie, W.-Z., et al. (2022) Citrus Pan-Genome to Breeding Database (CPBD): A comprehensive genome database for citrus breeding. Mol Plant. 15: 1503–1505.

26. Liu, J., Zhu, Y., Zhang, J., Sun, L., Sheng, J.-Z., Tan, Z., et al. (2025) Metabolic engineering and strain mating of *Yarrowia lipolytica* for sustainable production of prenylated aromatic compounds. ACS Sustain Chem Eng. 13: 3149–3159.

27. Matsumoto, Y., Kubo, T., Itami, Y., Islam, Md.Z., Watanabe, S., and Kotoda, N. (2022) QTL mapping of polymethoxyflavone (PMF) accumulation in citrus. Tree Genet Genomes. 18: 7.

28. Minh, B.Q., Schmidt, H.A., Chernomor, O., Schrempf, D., Woodhams, M.D., von Haeseler, A., et al. (2020) IQ-TREE 2: New models and efficient methods for phylogenetic inference in the genomic era. Mol Biol Evol. 37: 1530–1534.

29. Mousavi, S.H., Davari, A.-S., Iranshahi, M., Sabouri-Rad, S., and Tayarani Najaran, Z. (2015) Comparative analysis of the cytotoxic effect of 7-prenyloxycoumarin compounds and herniarin on MCF-7 cell line. Avicenna J Phytomedicine. 5: 520–530.

30. Munakata, R., Inoue, T., Koeduka, T., Karamat, F., Olry, A., Sugiyama, A., et al. (2014) Molecular cloning and characterization of a geranyl diphosphate-specific aromatic prenyltransferase from lemon. Plant Physiol. 166: 80–90.

31. Munakata, R., Olry, A., Karamat, F., Courdavault, V., Sugiyama, A., Date, Y., et al. (2016) Molecular evolution of parsnip (*Pastinaca sativa*) membrane-bound prenyltransferases for linear and/or angular furanocoumarin biosynthesis. New Phytol. 211: 332–344.

32. Munakata, R., Olry, A., Takemura, T., Tatsumi, K., Ichino, T., Villard, C., et al. (2021) Parallel evolution of UbiA superfamily proteins into aromatic *O*-prenyltransferases in plants. Proc Natl Acad Sci. 118: e2022294118.

33. Munakata, R., Takemura, T., Tatsumi, K., Moriyoshi, E., Yanagihara, K., Sugiyama, A., et al. (2019) Isolation of *Artemisia capillaris* membrane-bound di-prenyltransferase for phenylpropanoids and redesign of artepillin C in yeast. Commun Biol. 2: 384.

34. Munakata, R., and Yazaki, K. (2024) How did plants evolve the prenylation of specialized phenolic metabolites by means of UbiA prenyltransferases? Curr Opin Plant Biol. 81: 102601.

35. Murakami, A., Kuki, W., Takahashi, Y., Yonei, H., Nakamura, Y., Ohto, Y., et al. (1997) Auraptene, a citrus coumarin, inhibits 12-0-tetradecanoylphorbol-13-acetate-induced tumor promotion in ICR mouse skin, possibly through suppression of superoxide generation in leukocytes. Jpn J Cancer Res Gann. 88: 443–452.

36. Murray, R.D.H. (2002) The naturally occurring coumarins. Fortschritte Chem Org Naturstoffe Prog Chem Org Nat Prod Progres Dans Chim Subst Org Nat. 83: 1–619.

37. Murray, R.D.H., Méndez, J., and Brown, S.A. (1982) The natural coumarins: occurrence, chemistry, and biochemistry. Wiley.

38. Nakagawa, T., Suzuki, T., Murata, S., Nakamura, S., Hino, T., MAEO, K., et al. (2007) Improved gateway binary vectors: high-performance vectors for creation of fusion constructs in transgenic analysis of plants. Biosci Biotechnol Biochem. 71: 2095–2100.

39. Neal, J.J., and Wu, D. (1994) Inhibition of Insect Cytochromes P450 by Furanocoumarins. Pestic Biochem Physiol. 50: 43–50.

40. Ogawa, K., Kawasaki, A., Yoshida, T., Nesumi, H., Nakano, M., Ikoma, Y., et al. (2000) Evaluation of Auraptene Content in Citrus Fruits and Their Products. J Agric Food Chem. 48: 1763–1769.

41. Omura, M., and Shimada, T. (2016) Citrus breeding, genetics and genomics in Japan. Breed Sci. 66: 3–17.

42. Pailee, P., Prawat, H., Ploypradith, P., Mahidol, C., Ruchirawat, S., and Prachyawarakorn, V. (2020) Atalantiaphyllines A-G, prenylated acridones from *Atalantia monophylla* DC. and their aromatase inhibition and cytotoxic activities. Phytochemistry. 180: 112525.

43. R Core Team (2022) R: A language and environment for statistical computing. R Foundation for Statistical Computing, Vienna, Austria [WWW Document]. URL https://www.R-project.org/

44. Sasaki, K., Mito, K., Ohara, K., Yamamoto, H., and Yazaki, K. (2008) Cloning and Characterization of Naringenin 8-Prenyltransferase, a Flavonoid-Specific Prenyltransferase of Sophora flavescens. Plant Physiol. 146: 1075–1084.

45. Sevrioukova, I.F. (2019) Structural insights into the interaction of cytochrome P450 3A4 with suicide substrates: mibefradil, azamulin and 6′,7′-dihydroxybergamottin. Int J Mol Sci. 20: 4245.

46. Tanaya, R., Kodama, T., Lee, Y.-E., Yasuno, Y., Shinada, T., Takahashi, H., et al. (2023) Catalytic Potential of *Cannabis* Prenyltransferase to Expand Cannabinoid Scaffold Diversity. Org Lett. 25: 8601–8605.

47. Tayarani-Najaran, Z., Tayarani-Najaran, N., and Eghbali, S. (2021) A review of auraptene as an anticancer agent. Front Pharmacol. 12.

48. Thumuluri, V., Almagro Armenteros, J.J., Johansen, A.R., Nielsen, H., and Winther, O. (2022) DeepLoc 2.0: multi-label subcellular localization prediction using protein language models. Nucleic Acids Res. 50: W228–W234.

49. Tsurumaru, Y., Sasaki, K., Miyawaki, T., Momma, T., Umemoto, N., and Yazaki, K. (2010) An aromatic prenyltransferase-like gene *HlPT-1* preferentially expressed in lupulin glands of hop. Plant Biotechnol. 27: 199–204.

50. Vialart, G., Hehn, A., Olry, A., Ito, K., Krieger, C., Larbat, R., et al. (2012) A 2-oxoglutarate-dependent dioxygenase from Ruta graveolens L. exhibits p-coumaroyl CoA 2′-hydroxylase activity (C2′H): a missing step in the synthesis of umbelliferone in plants. Plant J. 70: 460–470.

51. Voinnet, O., Rivas, S., Mestre, P., and Baulcombe, D. (2003) An enhanced transient expression system in plants based on suppression of gene silencing by the p19 protein of tomato bushy stunt virus. Plant J. 33: 949–956.

52. Voo, S.S., Grimes, H.D., and Lange, B.M. (2012) Assessing the biosynthetic capabilities of secretory glands in *citrus* peel. Plant Physiol. 159: 81–94.

53. Wang, P., Fan, Z., Wei, W., Yang, C., Wang, Y., Shen, X., et al. (2023) Biosynthesis of the plant coumarin osthole by engineered *Saccharomyces cerevisiae*. ACS Synth Biol. 12: 2455–2462.

54. Wang, X., Xu, Y., Zhang, S., Cao, L., Huang, Y., Cheng, J., et al. (2017) Genomic analyses of primitive, wild and cultivated citrus provide insights into asexual reproduction. Nat Genet. 49: 765–772.

55. Wu, G.A., Terol, J., Ibanez, V., López-García, A., Pérez-Román, E., Borredá, C., et al. (2018) Genomics of the origin and evolution of Citrus. Nature. 554: 311–316.

56. Yazaki, K., Kunihisa, M., Fujisaki, T., and Sato, F. (2002) Geranyl diphosphate:4-hydroxybenzoate geranyltransferase from *Lithospermum erythrorhizon*. Cloning and characterization of a key enzyme in shikonin biosynthesis. J Biol Chem. 277: 6240–6246.

57. Yu, Y., Bai, J., Chen, C., Plotto, A., Yu, Q., Baldwin, E.A., et al. (2017) Identification of QTLs controlling aroma volatiles using a ‘Fortune’ x ‘Murcott’ (*Citrus reticulata*) population. BMC Genomics. 18: 646.

58. Zhang, Y., Jiao, D., Shen, C., Zhou, J., Guo, J., Yang, J., et al. (2025) Plant prenyltransferases: Diversity, catalytic activities, mechanisms, and application in heterologous production of prenylated natural products. J Integr Plant Biol. n/a.

59. Zhao, J., Bao, X., Li, C., Shen, Y., and Hou, J. (2016) Improving monoterpene geraniol production through geranyl diphosphate synthesis regulation in *Saccharomyces cerevisiae*. Appl Microbiol Biotechnol. 100: 4561–4571.

