## Supplementary Tables and Figures for "A membrane-bound aromatic *O*-prenyltransferase catalyzes the last reaction step in citrus auraptene biosynthesis"

**Table S1.** PT sequences used for *in silico* analysis.

| PT in primary metabolism | Plant species | Accession No. |
|---|---|---|
| **ABC4s in Phylloquinone biosynthesis** | | |
| AtABC4 | *Arabidopsis thaliana* | NP_001117518.1 |
| CsABC4 | *Citrus sinensis* | XP_024948946.1 |
| DcABC4 | *Daucus carota* | XP_017230268.1 |
| GmABC4 | *Glycine max* | XP_003532605.1 |
| OsABC4 | *Oryza sativa* | NP_001049226.1 |
| ZmABC4 | *Zea mays* | NP_001152170.1 |
| **ATGs in Chrolophyll biosynthesis** | | |
| AtATG4 | *Arabidopsis thaliana* | NP_190750.1 |
| CsATG4 | *Citrus sinensis* | XP_006481064.1 |
| DcATG4 | *Daucus carota* | XP_017243789.1 |
| GmATG4 | *Glycine max* | NP_001239633.1 |
| OsATG4 | *Oryza sativa* | ABO31092.1 |
| ZmATG4 | *Zea mays* | NP_001142204.1 |
| **COX10s in Heam *a* biosynthesis** | | |
| AtCOX10 | *Arabidopsis thaliana* | NP_566019.1 |
| CsCOX10 | *Citrus sinensis* | XP_006464403.1 |
| DcCOX10 | *Daucus carota* | XP_017227264.1 |
| GmCOX10 | *Glycine max* | XP_003556552.1 |
| OsCOX10 | *Oryza sativa* | EEC70799.1 |
| ZmCOX10 | *Zea mays* | AFW89544.1 |
| **PPTs in Ubiquinone biosynthesis** | | |
| AtPPT1 | *Arabidopsis thaliana* | NP_567688.1 |
| CsPPT | *Citrus sinensis* | XP_015387502.1 |
| DcPPT | *Daucus carota* | XP_017222930.1 |
| GmPPT | *Glycine max* | XP_006602724.1 |
| OsPPT1 | *Oryza sativa* | BAE96574.1 |
| ZmPPT | *Zea mays* | NP_001148558.1 |
| **VTE2-1s in Tocopherol biosynthesis** | | |
| AtVTE2-1 | *Arabidopsis thaliana* | NP_849984.1 |
| CsVTE2-1 | *Citrus sinensis* | XP_006474206.1 |
| DcVTE2-1 | *Daucus carota* | XP_017253952.1 |
| GmVTE2-1 | *Glycine max* | NP_001241496.1 |
| TaVTE2-1 | *Triticum aestivum* | ABB70123.1 |
| ZmVTE2-1 | *Zea mays* | ACG45339.1 |
| **VTE2-2s in Plastoquinone biosynthesis** | | |
| AtVTE2-2 | *Arabidopsis thaliana* | NP_001154609.1 |
| CsVTE2-2 | *Citrus sinensis* | XP_006481679.1 |
| DcVTE2-2 | *Daucus carota* | XP_017246707.1 |
| GmVTE2-2 | *Glycine max* | NP_001237900.1 |
| OsVTE2-2 | *Oryza sativa* | XP_015646905.1 |
| ZmVTE2-2 | *Zea mays* | NP_001146703.1 |

**Table S1.** PT sequences used for *in silico* analysis. *-continued*

| VTE2-1-related PTs | Plant species | Accession No. |
|---|---|---|
| **Apiaceae** | | |
| PcPT | *Petroselinum crispum* | BAO31627.1 |
| PsPT1 | *Pastinaca sativa* | AJW31563.1 |
| PsPT2 | *Pastinaca sativa* | AJW31564.1 |
| PpPT1 | *Peucedanum praeruptorum* | WIL06374.1 |
| PpPT2 | *Peucedanum praeruptorum* | WIL06375.1 |
| PpPT3 | *Peucedanum praeruptorum* | WIL06376.1 |
| **Asteraceae** | | |
| AcPT1 | *Artemisia capillaris* | BBG56301.1 |
| **Ericaceae** | | |
| RdPT1 | *Rhododendron dauricum* | BBD96134.1 |
| **Fabaceae** | | |
| AhR3'DT-1 | *Arachis hypogaea* | AQM74173.1 |
| AhR3'DT-2 | *Arachis hypogaea* | AQM74174.1 |
| AhR3'DT-3 | *Arachis hypogaea* | AQM74175.1 |
| AhR3'DT-4 | *Arachis hypogaea* | AQM74176.1 |
| AhR4DT-1 | *Arachis hypogaea* | AQM74172.1 |
| GmC4DT | *Glycine max* | BAW32575.1 |
| GmG2DT | *Glycine max* | BAW32578.1 |
| GmG4DT | *Glycine max* | NP_001235990.1 |
| GmIDT1 | *Glycine max* | BAW32576.1 |
| GmIDT2 | *Glycine max* | BAW32577.1 |
| GmIDT3 | *Glycine max* | XP_014618511.1 |
| GmPT01 | *Glycine max* | KRH76147.1 |
| GuA6DT | *Glycyrrhiza uralensis* | AIT11912.1 |
| GuILDT | *Glycyrrhiza uralensis* | AMR58303.1 |
| LaPT1 | *Lupinus albus* | AER35706.1 |
| LaPT2 | *Lupinus albus* | AWK21939.1 |
| LaG6DT1 | *Lupinus albus* | QYF10880.1 |
| LaG6DT2 | *Lupinus albus* | QYF10881.1 |
| LjG6DT | *Lotus japonicus* | ARV85585.1 |
| PcM4DT | *Psoralea corylifolia* | AYV64464.1 |
| SfFPT | *Sophora flavescens* | AHA36633.1 |
| SfG6DT | *Sophora flavescens* | BAK52291.1 |
| SfiLDT | *Sophora flavescens* | BAK52290.1 |
| SfN8DT-1 | *Sophora flavescens* | BAG12671.1 |
| SfN8DT-2 | *Sophora flavescens* | BAG12673.1 |
| SfN8DT-3 | *Sophora flavescens* | BAK52289.1 |
| **Poaceae** | | |
| HvHGGT | *Hordeum vulgare* | AAP43911.1 |
| OsHGGT | *Oryza sativa* | AAP43913.1 |
| TaHGGT | *Triticum aestivum* | AAP43912.1 |
| ZmHGGT | *Zea mays* | XP_008659772.1 |

**Table S1.** PT sequences used for *in silico* analysis. *-continued*

| VTE2-1-related PTs | Plant species | Accession No. |
|---|---|---|
| **Rutaceae** | | |
| ClPT1 | *Citrus limon* | BAP27988.1 |
| CpPT1 | *Citrus* × *paradisi* | BCH36128.1 |
| CpPT2 | *Citrus* × *paradisi* | BCH36129.1 |
| CpPT3 | *Citrus* × *paradisi* | BCH36130.1 |
| CmiPT1a | *Citrus micrantha* | BCH36131.1 |
| CmiPT1b | *Citrus micrantha* | BCH36132.1 |
| MePT1 | *Murraya exotica* | WJW73955.1 |
| MePT2 | *Murraya exotica* | WJW73956.1 |
| **Berberidaceae** | | |
| EpPT8 | *Epimedium pubescens* | UYI00165.1 |
| EsPT2 | *Epimedium sagittatum* | OENC2.p (BMDC) |

**Table S1.** PT sequences used for *in silico* analysis. *-continued*

| VTE2-2- and PPT-related PTs | Plant species | Accession No. |
|---|---|---|
| **Cannabaceae** | | |
| CsPT3 | *Cannabis sativa* | DAC76713.1 |
| CsPT4 | *Cannabis sativa* | DAC76710.1 |
| HlPT-1 | *Humulus lupulus* | BAJ61049.1 |
| HlPT-2 | *Humulus lupulus* | AJD80255.1 |
| **Hypericaceae** | | |
| HcPT8pat | *Hypericum calycinum* | AZK16227.1 |
| HcPT8px | *Hypericum calycinum* | AZK16226.1 |
| HsPT8pat | *Hypericum sampsonii* | AZK16225.1 |
| HsPT8px | *Hypericum sampsonii* | AZK16224.1 |
| HpPT4px-v1 | *Hypericum perforatum* | WIL60111.1 |
| HpPT4px-v2 | *Hypericum perforatum* | WIL60112.1 |
| HpPT4px-v3 | *Hypericum perforatum* | WIL60113.1 |
| HpPT4px-v4 | *Hypericum perforatum* | WIL60114.1 |
| HpPT4px-sh | *Hypericum perforatum* | WIL60115.1 |
| **Moraceae** | | |
| CtIDT | *Cudrania tricuspidata* | AJD80983.1 |
| FcPT1a | *Ficus carica* | BBC82715.1 |
| FcPT1b | *Ficus carica* | BBC82716.1 |
| MaIDT | *Morus alba* | AJD80982.1 |
| MaOGT | *Morus alba* | AXN57307.1 |
| **Boraginaceae** | | |
| AePGT | *Arnebia euchroma* | ABD59796.2 |
| AePGT4 | *Arnebia euchroma* | ANC67957.1 |
| AePGT6 | *Arnebia euchroma* | ANC67959.1 |
| LePGT1 | *Lithospermum erythrorhizon* | BAB84122.1 |
| LePGT2 | *Lithospermum erythrorhizon* | BAB84123.1 |
| **Fabaceae** | | |
| GLYMA_02G168000 | *Glycine max* | KRH71769.1 |

UbiA PTs are involved in primary and specialized metabolism (VTE2-1-, VTE2-2-, and PPT-related PTs) in plants.

**Table S2.** *In vitro* functional screening of CpPT4

| No. | Prenyl acceptor | Prenyl donor | | | |
|---|---|---|---|---|---|
| | | DMAPP (C5) | GPP (C10) | FPP (C15) | GGPP (C20) |
| **Coumarins (simple coumarins and furanocoumarins)** | | | | | |
| 1 | Umbelliferone (7-Hydroxycoumarin) | N.D. | + | N.D. | N.T. |
| 2 | 6-Hydroxycoumarin | N.D. | N.D. | N.D. | N.T. |
| 3 | 5,7-Dihydroxycoumarin | N.D. | + | N.D. | N.T. |
| 4 | Esculetin (6,7-Dihydroxycomarin) | N.D. | + | N.D. | N.T. |
| 5 | 5-Methoxy-7-hydroxycoumarin | N.D. | N.D. | N.D. | N.T. |
| 6 | Scopoletin (7-Hydroxy-6-Methoxycoumarin) | N.D. | + | N.D. | N.T. |
| 7 | 5-Hydroxy-7-methoxycoumarin | N.D. | N.D. | N.D. | N.T. |
| 8 | Isoscopoletin (6-Hydroxy-7-methoxycoumarin) | N.D. | N.D. | N.D. | N.T. |
| 9 | 7-Methoxy-8-hydroxycoumarin | N.D. | N.D. | N.D. | N.T. |
| 10 | Limettin (5,7-Dimethoxycoumarin) | N.D. | N.D. | N.D. | N.T. |
| 11 | 7-Aminocoumarin | N.D. | + | N.D. | N.T. |
| 12 | Bergaptol (5-Hydroxypsoralen) | N.D. | N.D. | N.D. | N.T. |
| 13 | Xanthotoxol (8-Hydroxypsoralen) | N.D. | N.D. | N.D. | N.T. |
| **Non-coumarin molecules** | | | | | |
| 14 | *p*-Coumaryl alcohol | N.D. | N.D. | N.D. | N.T. |
| 15 | *p*-Coumaraldehyde | N.D. | N.D. | N.D. | N.T. |
| 16 | *p*-Coumaric acid | N.D. | N.D. | N.D. | N.T. |
| 17 | Ferulic acid | N.D. | N.D. | N.D. | N.T. |
| 18 | 4-Hydroxyquinolin-2(1H)-one | N.D. | N.D. | N.D. | N.T. |
| 19 | 4-Hydroxy-*N*-methylquinolone | N.D. | N.D. | N.D. | N.T. |
| 20 | Naringenin | N.D. | N.D. | N.D. | N.T. |
| 21 | Homogentisic acid | N.D. | N.D. | N.D. | N.D. |

The enzymatic function of recombinant CpPT4 was screened using simple (bicyclic) coumarins, furanocoumarins, and other aromatic molecules as prenyl acceptors and DMAPP, GPP, FPP, and GGPP as prenyl donors (n = 1). Substrate pairs resulting in enzymatic products are marked with pluses. N.D., not detected. N.T., not tested.

**Table S3.** Summary of BLAST analysis for identifying orthologs in citrus genomes.

| Genome | Hit locs | Coverage | Max Identity | Bitscore | E-value | Subject Start | Subject End | Strand | Ortholog |
|---|---|---|---|---|---|---|---|---|---|
| Atalantia_buxfoliata.v1.0.genome | scaffold9675_locus1 | 0.959 | 98.85 | 1943.8 | 2.95E-131 | 2087866 | 2091455 | + | ortholog |
| | scaffold23061_locus1 | 0.355 | 100 | 788.8 | 1.57E-34 | 49980 | 58382 | - | non-ortholog |
| | scaffold26253_locus1 | 0.089 | 83.02 | 95.3 | 7.46E-18 | 853062 | 853166 | + | non-ortholog |
| | scaffold9675_locus2 | 0.085 | 91.09 | 137 | 1.22E-30 | 2102676 | 2102776 | + | non-ortholog |
| Citrus_clementina_v1 | scaffold_7_locus1 | 0.438 | 95 | 675.9 | 8.91E-42 | 14108710 | 14115418 | - | non-ortholog |
| | scaffold_2_locus1 | 0.089 | 83.96 | 100 | 1.53E-19 | 12651152 | 12651256 | + | non-ortholog |
| | scaffold_1_locus2 | 0.076 | 93.33 | 134 | 1.51E-29 | 1347123 | 1347212 | + | non-ortholog |
| | scaffold_1_locus1 | 0.043 | 100 | 95.3 | 7.13E-18 | 1327901 | 1327951 | + | ortholog |
| Citrus_grandis_Cupi_Majiayou.v1.0.genome | chr4_locus2 | 1 | 100 | 2239.7 | 1.56E-149 | 37330058 | 37334632 | - | ortholog |
| | chr1_locus1 | 0.438 | 95.24 | 669.1 | 1.82E-39 | 21495492 | 21502205 | + | non-ortholog |
| | chr2_locus1 | 0.084 | 84.85 | 100 | 1.87E-19 | 28758605 | 28758703 | - | non-ortholog |
| | chr2_locus2 | 0.084 | 84.85 | 100 | 1.87E-19 | 28783799 | 28783897 | - | non-ortholog |
| Citrus_grandis_Wanbaiyou.v1.0.genome | chr4_locus1 | 0.926 | 100 | 2015.7 | 1.47E-144 | 1824926 | 1829510 | + | ortholog |
| | chr1_locus1 | 0.296 | 94.87 | 505.1 | 7.96E-38 | 17250041 | 17256795 | + | non-ortholog |
| | chrUn_locus2 | 0.296 | 94.87 | 505.1 | 7.96E-38 | 38114562 | 38121334 | + | non-ortholog |
| | chr1_locus2 | 0.291 | 100 | 505.1 | 7.96E-38 | 18841191 | 18847960 | - | non-ortholog |
| Citrus_ichangensis.v1.0.genome | scaffold_2_locus2 | 1 | 100 | 2133.7 | 1.54E-134 | 228197 | 232709 | - | ortholog |
| | scaffold_891_locus1 | 0.434 | 100 | 938.4 | 1.75E-44 | 46258 | 48081 | - | non-ortholog |
| | scaffold_873_locus1 | 0.404 | 95.24 | 607 | 4.91E-40 | 33671 | 36518 | + | non-ortholog |
| | scaffold_120_locus1 | 0.259 | 95.24 | 445 | 1.77E-39 | 40005 | 41478 | - | non-ortholog |
| Citrus_medica.v1.0.genome | scaffold_295_locus2 | 1 | 100 | 2084.2 | 8.21E-128 | 40121 | 44686 | - | ortholog |
| | scaffold_194_locus1 | 0.432 | 100 | 661.2 | 2.59E-38 | 384112 | 391217 | + | non-ortholog |
| | scaffold_192_locus1 | 0.089 | 82.86 | 95.3 | 9.59E-18 | 485085 | 485189 | + | non-ortholog |
| | scaffold_192_locus2 | 0.089 | 82.86 | 95.3 | 9.59E-18 | 506574 | 506678 | + | non-ortholog |
| Citrus_reticulata_scaf | scaffold497_cov94_locus1 | 0.964 | 100 | 2066.3 | 3.21E-136 | 1069862 | 1074297 | + | ortholog |
| | scaffold85922_cov81_locus1 | 0.263 | 95.24 | 455 | 1.71E-39 | 776405 | 777886 | - | non-ortholog |
| | scaffold1186_cov86_locus1 | 0.089 | 84.91 | 106 | 3.77E-21 | 180112 | 180216 | + | non-ortholog |
| | scaffold1186_cov86_locus2 | 0.089 | 83.81 | 100 | 1.75E-19 | 202313 | 202417 | + | non-ortholog |
| Citrus_sinensis.v3.0.genome | chr4_locus1 | 1 | 100 | 2131.1 | 3.13E-136 | 3493206 | 3499285 | + | ortholog |
| | chr1_locus1 | 0.296 | 94.87 | 505.1 | 7.75E-38 | 15995795 | 16002579 | + | non-ortholog |
| | chr1_locus2 | 0.296 | 94.87 | 505.1 | 7.75E-38 | 16036189 | 16042967 | + | non-ortholog |
| | chr1_locus3 | 0.296 | 94.87 | 505.1 | 7.75E-38 | 16076582 | 16083358 | + | non-ortholog |
| Fortunella_hindsii.v1.0.genome | tig00001125_arrow_pilon_locus2 | 1 | 100 | 2151.7 | 1.62E-134 | 2687942 | 2692315 | - | ortholog |
| | tig00008652_arrow_pilon_locus1 | 0.405 | 95.24 | 612 | 5.14E-40 | 49003 | 52122 | - | non-ortholog |
| | tig00040300_arrow_pilon_locus1 | 0.405 | 95.24 | 612 | 5.14E-40 | 717356 | 720479 | - | non-ortholog |
| | tig00000053_arrow_pilon_locus1 | 0.084 | 84.85 | 100 | 1.90E-19 | 4520036 | 4520134 | - | non-ortholog |
| Poncirus_trifoliata_v1.3 | scaffold_1_locus1 | 0.926 | 100 | 2122.2 | 2.45E-141 | 1349274 | 1359532 | + | ortholog |
| | scaffold_7_locus1 | 0.263 | 95.24 | 444 | 1.31E-39 | 12521147 | 12522632 | - | non-ortholog |
| | scaffold_2_locus4 | 0.089 | 85.85 | 111 | 6.23E-23 | 8688246 | 8688350 | + | non-ortholog |
| | scaffold_2_locus1 | 0.065 | 90.91 | 104 | 1.04E-20 | 8609435 | 8609511 | + | non-ortholog |

BLASTN searches were performed using *CpPT4* CDS against 10 citrus genomes with -outfmt 6, max_target_seqs=5000, and max_hsps=5000. Multiple high-scoring segment pairs for the same query–subject pair were merged on the query side to calculate coverage, and subject intervals within 10 kb were considered to belong to the same locus. For each query–hit locus pair, the maximum identity (Max Identity), total bitscore, and best e-value were calculated. Top hits that showed the highest Bitscore and lowest E-values were designated as orthologs, except for *C. clementina* whose top hit showed a much lower bioscore and a much higher E-value than the top hits of the other tested species. For this species, the schaffold1_locus1 was assigned as an ortholog because its full hit sequence showed 100% identity with the corresponding region of CpPT4. Moreover, a large deletion was detected directly downstream of this locus, which greatly affected the bitscore and E-value of this locus.

**Table S4.** Polymorphism in *CpPT4* ortholog locus of *C. sinensis*, *C. clementina*, *C. reticulata* and *C. grandis.*

| Species | Varieties | Mutation in CpPT4 ortholog locus | Tree replicates | Leaf replicates | Amplicons | Sampling | DNA extraction kit | JP number |
|---|---|---|---|---|---|---|---|---|
| *C. sinensis* | Washington navel | + | 3 | 1 | 1 | Corsica | E.Z.N.A. Plant DNA Kit (Omega Bio-tek) | - |
| *C. sinensis* | Hamlin orange | + | 3 | 1 | 1 | Corsica | E.Z.N.A. Plant DNA Kit (Omega Bio-tek) | - |
| *C. sinensis* | Shamouti sweet orange | + | 3 | 1 | 1 | Corsica | E.Z.N.A. Plant DNA Kit (Omega Bio-tek) | - |
| *C.clementina* | Clementine caffin | + | 3 | 3 | 1 | Corsica | E.Z.N.A. Plant DNA Kit (Omega Bio-tek) | - |
| *C.clementina* | Clementine tomatera | + | 3 | 3 | 1 | Corsica | E.Z.N.A. Plant DNA Kit (Omega Bio-tek) | - |
| *C.clementina* | Clementine commune | + | 3 | 3 | 1 | Corsica | E.Z.N.A. Plant DNA Kit (Omega Bio-tek) | - |
| *C. reticulata* | Cleopatre | + | 2 | 3 | 1 | Corsica | E.Z.N.A. Plant DNA Kit (Omega Bio-tek) | - |
| *C. reticulata* | Beauty mandarin | + | 3 | 3 | 1 | Corsica | E.Z.N.A. Plant DNA Kit (Omega Bio-tek) | - |
| *C. reticulata* | Sunki mandarin | + | 3 | 1 | 1 | Corsica | E.Z.N.A. Plant DNA Kit (Omega Bio-tek) | - |
| *C. reticulata* | Dancy mandarin | + | 3 | 1 | 1 | Corsica | E.Z.N.A. Plant DNA Kit (Omega Bio-tek) | - |
| *C. grandis* | Chandler pummelo | - | 3 | 1 | 1 | Corsica | E.Z.N.A. Plant DNA Kit (Omega Bio-tek) | - |
| *C. grandis* | Timor pummelo | - | 3 | 1 | 1 | Corsica | E.Z.N.A. Plant DNA Kit (Omega Bio-tek) | - |
| *C. grandis* | Pink pummelo | - | 3 | 1 | 1 | Corsica | E.Z.N.A. Plant DNA Kit (Omega Bio-tek) | - |
| *C. grandis* | Tahiti pummelo | - | 1 | 2 | 1 | Corsica | E.Z.N.A. Plant DNA Kit (Omega Bio-tek) | - |
| *C. grandis* | Eingerdi pummelo | - | 1 | 2 | 1 | Corsica | E.Z.N.A. Plant DNA Kit (Omega Bio-tek) | - |
| *C. grandis* | Sunshine pummelo | - | 1 | 2 | 1 | Corsica | E.Z.N.A. Plant DNA Kit (Omega Bio-tek) | - |
| *C. grandis* | Eilot pummelo | - | 1 | 2 | 1 | Corsica | E.Z.N.A. Plant DNA Kit (Omega Bio-tek) | - |
| *C. grandis* | Reinking pummelo | - | 1 | 2 | 1 | Corsica | E.Z.N.A. Plant DNA Kit (Omega Bio-tek) | - |
| *C. grandis* | Sans pépin pummelo | - | 1 | 2 | 1 | Corsica | E.Z.N.A. Plant DNA Kit (Omega Bio-tek) | - |
| *C. grandis* | Kao Pan pummelo | - | 1 | 2 | 1 | Corsica | E.Z.N.A. Plant DNA Kit (Omega Bio-tek) | - |
| *C. grandis* | Kao pan | - | 1 | 2 | 1 | NARO | DNeasy Plant Pro Kit (Qiagen) | 117437 |
| *C. grandis* | Tanikawa buntan | - | 2 | 1 | 1 | Kindai | DNeasy Plant Pro Kit (Qiagen) | - |
| *C. grandis* | Tanikawa buntan | - | 1 | 2 | 1 | NARO | DNeasy Plant Pro Kit (Qiagen) | 117433 |
| *C. grandis* | Mei pummelo | - | 2 | 1 | 1 | Kindai | DNeasy Plant Pro Kit (Qiagen) | - |
| *C. grandis* | Yellow pummelo | - | 2 | 1 | 1 | Kindai | DNeasy Plant Pro Kit (Qiagen) | - |
| *C. grandis* | Otachibana | - | 2 | 1 | 1 | Kindai | DNeasy Plant Pro Kit (Qiagen) | - |
| *C. grandis* | Anseikan | - | 2 | 1 | 1 | Kindai | DNeasy Plant Pro Kit (Qiagen) | - |
| *C. grandis* | Anseikan | - | 1 | 2 | 1 | NARO | DNeasy Plant Pro Kit (Qiagen) | 117431 |
| *C. grandis* | Kuchinotsu No. 2 | - | 2 | 1 | 1 | Kindai | DNeasy Plant Pro Kit (Qiagen) | - |
| *C. grandis* | Hirado buntan | - | 2 | 1 | 1 | Kindai | DNeasy Plant Pro Kit (Qiagen) | - |
| *C. grandis* | Hirado buntan | - | 1 | 2 | 1 | NARO | DNeasy Plant Pro Kit (Qiagen) | 171507 |
| *C. grandis* | Banpeiyu (Wanbaiyou) | - | 1 | 2 | 1 | NARO | DNeasy Plant Pro Kit (Qiagen) | 171506 |
| *C. grandis* | Kouchi peiyu | - | 1 | 2 | 1 | NARO | DNeasy Plant Pro Kit (Qiagen) | 113248 |
| *C. grandis* | Morrison | - | 1 | 2 | 1 | NARO | DNeasy Plant Pro Kit (Qiagen) | 113376 |
| *C. grandis* | Sha tian you | - | 1 | 2 | 1 | NARO | DNeasy Plant Pro Kit (Qiagen) | 115504 |
| *C. grandis* | Andoukan | - | 1 | 2 | 1 | NARO | DNeasy Plant Pro Kit (Qiagen) | 115520 |
| *C. grandis* | Hougen buntan | - | 1 | 2 | 1 | NARO | DNeasy Plant Pro Kit (Qiagen) | 117282 |
| *C. grandis* | Matou buntan | - | 1 | 2 | 1 | NARO | DNeasy Plant Pro Kit (Qiagen) | 117283 |
| *C. grandis* | Oogonkitsu | - | 1 | 2 | 1 | NARO | DNeasy Plant Pro Kit (Qiagen) | 117410 |
| *C. grandis* | Banoukan | - | 1 | 2 | 1 | NARO | DNeasy Plant Pro Kit (Qiagen) | 117432 |
| *C. grandis* | Bangkok buntan-hirakei | - | 1 | 2 | 1 | NARO | DNeasy Plant Pro Kit (Qiagen) | 168894 |
| *C. grandis* | Vietnum buntan 2 | - | 1 | 2 | 1 | NARO | DNeasy Plant Pro Kit (Qiagen) | 168896 |
| *C. grandis* | Vietnum buntan 22 | - | 1 | 2 | 1 | NARO | DNeasy Plant Pro Kit (Qiagen) | 168897 |
| *C. grandis* | Nagasaki zabon 3 daime | - | 1 | 2 | 1 | NARO | DNeasy Plant Pro Kit (Qiagen) | 244576 |
| *C. grandis* | Tosa buntan chimera | - | 1 | 2 | 1 | NARO | DNeasy Plant Pro Kit (Qiagen) | 245243 |
| *C. grandis* | Buntan B kei chimera | - | 1 | 2 | 1 | NARO | DNeasy Plant Pro Kit (Qiagen) | 245244 |
| *C. grandis* | CRC2240 chimera | - | 1 | 2 | 1 | NARO | DNeasy Plant Pro Kit (Qiagen) | 245245 |

Varieties with mutations leading to loss of function confirmed by PCR sequencing are marked with a plus and varieties without the mutations are marked with a minus. The number of iterations of tree, leaf, and PCR amplicons are noted.

**Table S5.** Primers used in this work.

| Primer name | Sequence (5'-3') |
|---|---|
| CpPT4_5'UTR_Fw | GCTAAGGATCGGAGAGAGAGG |
| CpPT4_3'UTR_Rv | GTTGCCATTGCTATCTGTCG |
| CpPT4_TOPO_Fw | CACCATGGTTCATATGCATTCATG |
| CpPT4_TOPO_Rv | TCAACGTACAAAATGAATAAGGAGG |
| CpPT4_qRTPCR_Fw | TTTACGTGGTGGCCCTAAAC |
| CpPT4_qRTPCR_Rv | CTCCAGTTCCCGTTGAGAAG |
| CpEF1α_Fw | TAAGGATGGGCAGACCAGAG |
| CpEF1α_Rv | TGGTGGCATCCATCTTGTTA |
| CpPT4_TP210_Rv | GGTAGAAAGACAACGATACT |
| CpPT4_woStop_Rv | ACGTACAAAATGAATAAGGAGG |
| Csinensis_Fw | CAGTGGAAGCAAGTGGTCAA |
| Csinensis_Rv | GCGGTTAATCGCAGAACAAT |
| Cclementina_Fw | TTCCCAATCTTGAAGCTCGT |
| Cclementina_Rv | CAAAATCTTGAAGGGGGAGA |
| Cgrandis_Fw | TAACAAGCCCGATCTTCCAC |
| Cgrandis_Rv | GGGCACTTTACTTGGATGGA |

**Table S6.** LC/$MS^2$ conditions for analysis.

| Properties of Full MS | |
|---|---|
| **General** | |
| Polarity | positive |
| dd-$MS^2$ | Discovery |
| **Full MS** | |
| Resolution | 70,000 |
| Scan range | 50 to 750 m/z |
| **dd-$MS^2$ Discovery** | |
| Resolution | 17,500 |
| Isolation window | 1.6 m/z |
| (N)CE / stepped (N)CE | nce: 15, 30, 45 |
| AGC target | 1e5 |
| Maximum IT | 60 ms |

**Table S7.** Genome assemblies used for *CpPT4* ortholog identification.

| Species (Scientific Name) | Common Name | Cultivar/Strain | Genome Version | Database | Reference |
|---|---|---|---|---|---|
| *Citrus grandis* | Pummelo | Cupi Majiayou | v1.0 | Citrus Pan-genome to Breeding Database | (Lu *et al.*, 2022) |
| *Citrus grandis* | Pummelo | Wanbaiyou | v1.0 | Citrus Pan-genome to Breeding Database | (Wang *et al.*, 2017) |
| *Citrus sinensis* | Sweet orange | HZAU_DHSO_2021 | v3.0 | Citrus Pan-genome to Breeding Database | (Wang *et al.*, 2021) |
| *Citrus reticulata* | Mandarin | Mangshan | v1.0 | Citrus Pan-genome to Breeding Database | (Wang *et al.*, 2018) |
| *Citrus clementina* | Clementine | Clemenules | v1.0 | Phytozome | (Wu *et al.*, 2014) |
| *Atalantia buxifolia* | Chinese box orange | HKC | v1.0 | Citrus Pan-genome to Breeding Database | (Wang *et al.*, 2017) |
| *Fortunella hindsii* | Hongkong kumquat | S3y-45 | v1.0 | Citrus Pan-genome to Breeding Database | (Zhu *et al.*, 2019) |
| *Poncirus trifoliata* | Hardy orange | DPI 50-7 | v1.3.1 | Phytozome | (Peng *et al.*, 2020) |
| *Citrus ichangensis* | Papeda | XJC | v1.0 | Citrus Pan-genome to Breeding Database | (Wang *et al.*, 2017) |
| *Citrus medica* | Citron | XZ | v1.0 | Citrus Pan-genome to Breeding Database | (Wang *et al.*, 2017) |

### References


Lu, Z., Huang, Y., Mao, S., Wu, F., Liu, Y., Mao, X., et al. (2022) The high-quality genome of pummelo provides insights into the tissue-specific regulation of citric acid and anthocyanin during domestication. *Horticulture Research*. 9: uhac175.

Peng, Z., Bredeson, J.V., Wu, G.A., Shu, S., Rawat, N., Du, D., et al. (2020) A chromosome-scale reference genome of trifoliate orange (*Poncirus trifoliata*) provides insights into disease resistance, cold tolerance and genome evolution in Citrus. *The Plant Journal*. 104: 1215–1232.

Wang, X., Xu, Y., Zhang, S., Cao, L., Huang, Y., Cheng, J., et al. (2017) Genomic analyses of primitive, wild and cultivated citrus provide insights into asexual reproduction. *Nat Genet*. 49: 765–772.

Wang, L., He, F., Huang, Y., He, J., Yang, S., Zeng, J., et al. (2018) Genome of wild mandarin and domestication history of mandarin. *Molecular Plant*. 11: 1024–1037.

Wang, L., Huang, Y., Liu, Z., He, J., Jiang, X., He, F., et al. (2021) Somatic variations led to the selection of acidic and acidless orange cultivars. *Nat Plants*. 7: 954–965.

Wu, G.A., Prochnik, S., Jenkins, J., Salse, J., Hellsten, U., Murat, F., et al. (2014) Sequencing of diverse mandarin, pummelo and orange genomes reveals complex history of admixture during citrus domestication. *Nat Biotechnol*. 32: 656–662.

Zhu, C., Zheng, X., Huang, Y., Ye, J., Chen, P., Zhang, C., et al. (2019) Genome sequencing and CRISPR/Cas9 gene editing of an early flowering Mini-Citrus (*Fortunella hindsii*). *Plant Biotechnology Journal*. 17: 2199–2210.

```
                ....|....| ....|....| ....|....| ....|....| ....|....| ....|....|
                        1330       1340       1350       1360       1370       1380
Pummelo_HWB     ATCTGACCTT GAAATAGACA AGGTAACAGT TTATATATAT ATATATATAT ATAAAGCATT
c22985_g1_i1    ---------T GAAATAGACA AG-------- ---------- ---------- ----------
CpPT4           ATCTGACCTT GAAATAGACA AG-------- ---------- ---------- ----------

                ....|....| ....|....| ....|....| ....|....| ....|....| ....|....|
                        1570       1580       1590       1600       1610       1620
Pummelo_HWB     CTAATAATTC TGCTGAACAT TGTTAAAGAT TAACAAGCCC GATCTTCCAC TTGCTTCTGG
c22985_g1_i1    ---------- ---------- --------AT TAACAAGCCC GATCTTCCAC TTGCTTCTGG
CpPT4           ---------- ---------- --------AT TAACAAGCCC GATCTTCCAC TTGCTTCTGG

                ....|....| ....|....| ....|....| ....|....| ....|....| ....|....|
                        1630       1640       1650       1660       1670       1680
Pummelo_HWB     TGACTTCTCA ACGGGAACTG GAGTTGCAGT TTATTTAGTA TCTTCATTCA CGGTAAGAAA
c22985_g1_i1    TGACTTCTCA ACGGGAACTG GAGTTGCAGT TTATTTAGTA TCTTCATTCA CG--------
CpPT4           TGACTTCTCA ACGGGAACTG GAGTTGCAGT TTATTTAGTA TCTTCATTCA CG--------

                ....|....| ....|....| ....|....| ....|....| ....|....| ....|....|
                        2230       2240       2250       2260       2270       2280
Pummelo_HWB     TGTTTTTCAA TTTTAAGAGT --GGCTATGG GAGTTATATC TCGATCTCCA CCTCTTCTTC
c22985_g1_i1    ---------- -------AGT CTGGCTATGG GAGTTATGTC TCGATCTCCA CCTCTTCTCC
CpPT4           ---------- -------AGT CTGGCTATGG GAGTTATGTC TCGATCTCCA CCTCTTCTCC

                ....|....| ....|....| ....|....| ....|....| ....|....| ....|....|
                        2290       2300       2310       2320       2330       2340
Pummelo_HWB     TGGCCTTCAT CATCTGGTTT CTACTCGGAA CCGCTTACTC CATCCAAGTA AAGTGCCCAA
c22985_g1_i1    TGGCCCTCAT CATATGGTTT TTACTCGGAA CCGCTTACTC CATCCAA--- ----------
CpPT4           TGGCCCTCAT CATATGGTTT TTACTCGGAA CCGCTTACTC CATCCAA--- ----------

                ....|....| ....|....| ....|....| ....|....| ....|....| ....|....|
                        2590       2600       2610       2620       2630       2640
Pummelo_HWB     CTCAATATTG CAGCTTCCCT TTCTAAGATG GAAAGGACAT CCATTTCTTG CACCAACCTC
c22985_g1_i1    ---------- ---CTTCCCT TTCTAAGATG GAAAGGACAT CCATTTCTTG CACCAACCTC
CpPT4           ---------- ---CTTCCCT TTCTAAGATG GAAAGGACAT CCATTTCTTG CACCAACCTC

                ....|....| ....|....| ....|....| ....|....| ....|....| ....|....|
                        2650       2660       2670       2680       2690       2700
Pummelo_HWB     CATGATGATT TTGATGGGAC TTGTATATCA ATTGCTATTC TACATACACA TCCAGGTATA
c22985_g1_i1    CATGATGATT TTGATGGGAC TTGTATATCA ATTGCTATTC CACATACACA TACAG-----
CpPT4           CATGATGATT TTGATGGGAC TTGTATATCA ATTGCTATTC CACATACACA TACAG-----

                ....|....| ....|....| ....|....| ....|....| ....|....| ....|....|
                        3430       3440       3450       3460       3470       3480
Pummelo_HWB     AAATACGTGC TTGCAAGACC TGTAATAATT ACAAAACCAC TGGTGTTTGC TACCACTTTC
c22985_g1_i1    ---------- ---------- ---------- ---------- ---------- ----------
CpPT4           AAATACGTGC TTGCAAGACC TGTAATAATT ACAAAACCAC TGGTGTTTGC TACCACTTTC

                ....|....| ....|....| ....|....| ....|....| ....|....| ....|....|
                        3490       3500       3510       3520       3530       3540
Pummelo_HWB     ATGTCCGTAT TCTCCTTTGC AAATGGCCTT CTCAAGGTAC GGATAATTCA AAAATTGATG
c22985_g1_i1    ---------- ---------- ---------- ---------- ---------- ----------
CpPT4           ATGTCTGTAT TCTCCTTTGC AAATGGCCTT CTCAAG---- ---------- ----------
                          2nd Asp-rich motif     K

                ....|....| ....|....| ....|....| ....|....| ....|....| ....|....|
                        3730       3740       3750       3760       3770       3780
Pummelo_HWB     CATTAATTTA CTTGTTTCGA TTTTATAAAA GGATGTACCT GATGTGGATG GCGACAGAGA
c22985_g1_i1    ---------- ---------- ---------- ---------- ---------- ----------
CpPT4           ---------- ---------- ---------- -GATGTACCT GATGTGGATG GCGACAGAGA
                      2nd Asp-rich motif     D   V   P   D   V   D   G   D
```

**Figure S1.** Multiple alignment of the genome of pummelo, c22985_g1_i1, and *CpPT4*.

```
               ....|....| ....|....| ....|....| ....|....| ....|....| ....|....|
                      3790       3800       3810       3820       3830       3840
Pummelo_HWB    ATTTGGCATT CAGACGCTCA GTGTTATCAT AGGAAAAGAA AAAGTAAGTA AATTAAGAAT
c22985_g1_i1   ---------- ---------- ---------- ---------- ---------- ----------
CpPT4          ATTTGGCATT CAGACGCTCA GTGTTATCAT AGGAAAAGAA AAA------- ----------

               ....|....| ....|....| ....|....| ....|....| ....|....| ....|....|
                      4150       4160       4170       4180       4190       4200
Pummelo_HWB    ATATTCTTCT CAAAAGGTAT TTTGGCTTGG TGTTTATATG ATGTTAACTG CCTATGGAGC
c22985_g1_i1   ---------- ------GTAT TTTGGCTTGG TGTTTATATG ATGCTAACTG CCTATGGAGC
CpPT4          ---------- ------GTAT TTTGGCTTGG TGTTTATATG ATGCTAACTG CCTATGGAGC

               ....|....| ....|....| ....|....| ....|....| ....|....| ....|....|
                      4210       4220       4230       4240       4250       4260
Pummelo_HWB    TGCTATATTA GTTGGAACTT CTTCATCTTT TCTGCTAAGC AAGCTTGTCA CTGTAAGAAT
c22985_g1_i1   TGCTATATTA GTTGGAGCTT CTTCATCTTT TCTGCTAAGC AAGCTTGTCA CT--------
CpPT4          TGCTATATTA GTTGGAGCTT CTTCATCTTT TCTGCTAAGC AAGCTTGTCA CT--------

               ....|....| ....|....| ....|....| ....|....| ....|....| ....|....|
                      4270       4280       4290       4300       4310       4320
Pummelo_HWB    GGCATCATTT TGTAAGAAAT TTAGCTTGTG CTTGAATTTG ATTCAGTTAA TCATGAAGAC
c22985_g1_i1   ---------- ---------- ---------- ---------- ---------- ----------
CpPT4          ---------- ---------- ---------- ---------- ---------- ----------

               ....|....| ....|....| ....|....| ....|....| ....|....| ....|....|
                      4330       4340       4350       4360       4370       4380
Pummelo_HWB    CCTTTTTTTT TTTTTTTGGA AGATAATTTC TCACAGTATA CTTGCTTTCA TTCTGTGGCT
c22985_g1_i1   ---------- ---------- --ATAATTTC TCACAGTATA CTTGCTTTCA TTCTGTGGCT
CpPT4          ---------- ---------- --ATAATTTC TCACAGTATA CTTGCTTTCA TTCTGTGGCT

               ....|....| ....|....| ....|....| ....|....| ....|....| ....|....|
                      4390       4400       4410       4420       4430       4440
Pummelo_HWB    TCGAGCCCAA ACTGTTGATC TCTCCAATAA AGCATCAACA CTTTCTTTTT ACATGTTCAT
c22985_g1_i1   TCGAGCCCAA ACTGTTGATC TCTCCAATAA AGCGTCAACA CTTTCTTTTT ACATGTTCAT
CpPT4          TCGAGCCCAA ACTGTTGATC TCTCCAATAA AGCGTCAACA CTTTCTTTTT ACATGTTCAT

               ....|....| ....|....| ....|....| ....|....| ....|....| ....|....|
                      4450       4460       4470       4480       4490       4500
Pummelo_HWB    TTGGAAGGCA AGTGATTGGT CTTTAATATG AGAATATTGA ATCAAATCCA ATCATGTTCC
c22985_g1_i1   TTGGAAG--- ---------- ---------- ---------- ---------- ----------
CpPT4          TTGGAAG--- ---------- ---------- ---------- ---------- ----------

               ....|....| ....|....| ....|....| ....|....| ....|....| ....|....|
                      4510       4520       4530       4540       4550       4560
Pummelo_HWB    AATTACATAG CTTGAAACTG ACCTGTTATG TTATTGCATC TGCAGTTACA CTACGTTGAA
c22985_g1_i1   ---------- ---------- ---------- ---------- -----TTACA CTACGTTGAA
CpPT4          ---------- ---------- ---------- ---------- -----TTACA CTACGTTGAA

               ....|....| ....|....| ....|..
                      4570       4580
Pummelo_HWB    TACCTCCTTA TTCATTTTGT ACGTTGA
c22985_g1_i1   TACCTCCTTA TTCATTTTGT ACGTTGA
CpPT4          TACCTCCTTA TTCATTTTGT ACGTTGA
```

**Figure S1.** Multiple alignment of the genome of pummelo, c22985_g1_i1, and *CpPT4*. *–continued* The second aspartate-rich motif is highlighted by a blue square. The upper ruler indicates the base number from the start codon on the pummelo (cv. Wanbaiyou).

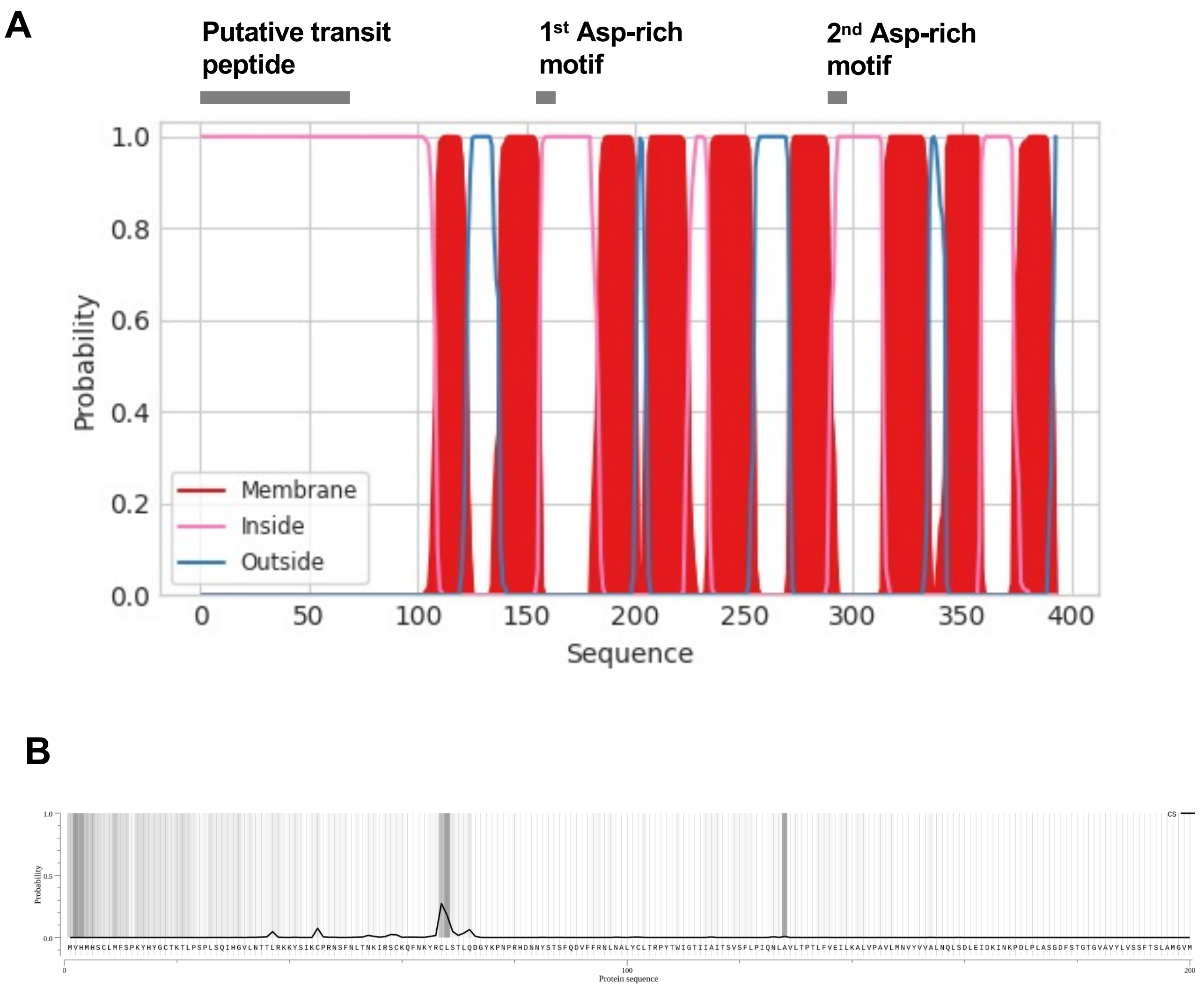


**Figure S2.** *In silico* analysis of the CpPT4 polypeptide.

# C

**CpPT4_prot**

**Predicted localizations:** Plastid

**Predicted signals:** Chloroplast transit peptide

| Localization | Cytoplasm | Nucleus | Extracellular | Cell membrane | Mitochondrion | Plastid | Endoplasmic reticulum | Lysosome/Vacuole | Golgi apparatus | Peroxisome |
|---|---|---|---|---|---|---|---|---|---|---|
| **Probability** | 0.0879 | 0.0984 | 0.0106 | 0.0619 | 0.2095 | 0.9330 | 0.0564 | 0.0559 | 0.0720 | 0.1160 |

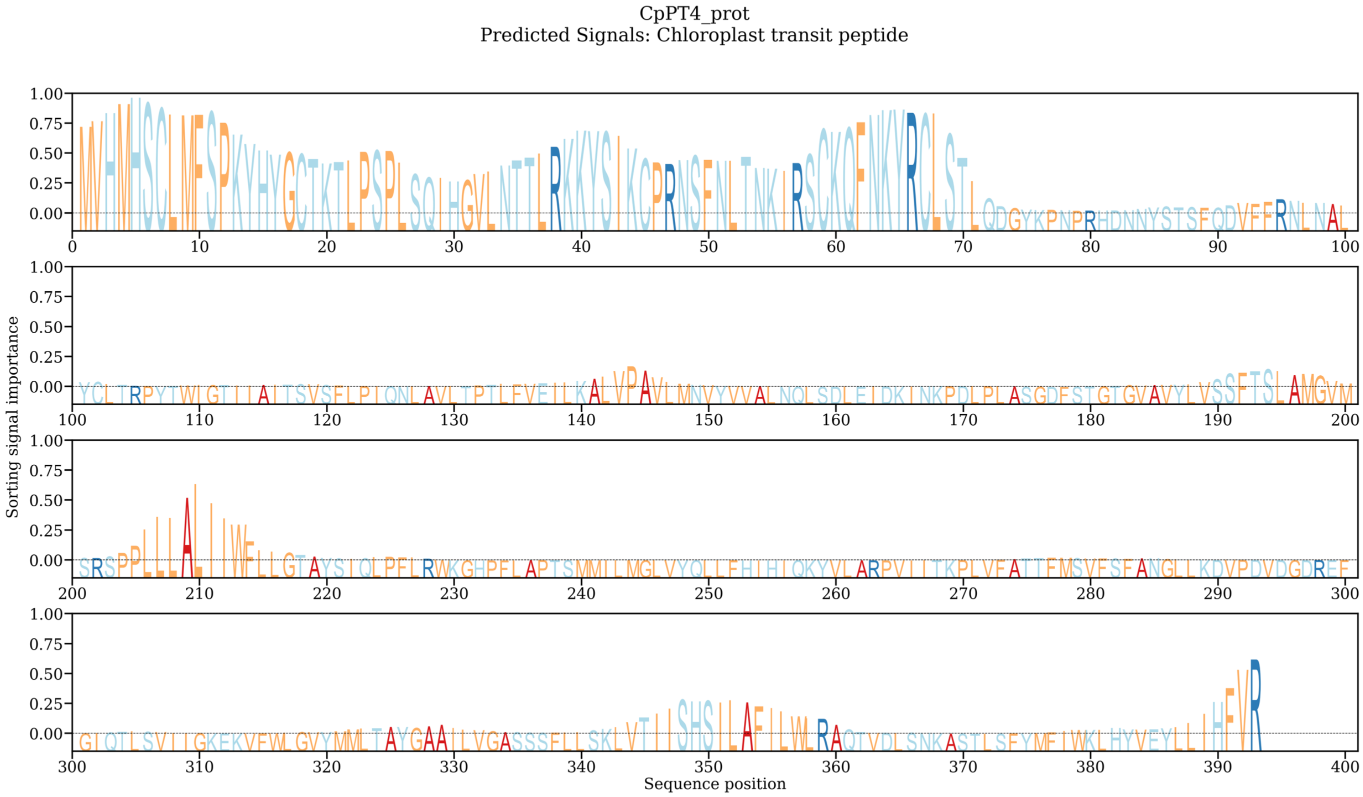


**Figure S2.** *In silico* analysis of the CpPT4 polypeptide. *–continued*

## D

```
CpPT4     -MVHMHSCLMFSPKYHY-----GCTKTLPSPLSQIHGVLNTTLRKKYSIKC-----PR-NSFNLT-NKI--------RSCKQFNKYRCLSTLQ
ClPT1     -MLQMHSNSSFSPKCYYPLQHAGCVKTLQLPLTKVHGGLNRSESKNYAIKC-----TQSDSFYST-NKIRNNENSSSRNCKPFNKYRVAVTLQ
CpPT1     MLLQMNLCSSFSLKYHP-LQQNGTVKTFQSPLTQIYGLANRRESNKYSVKG-----SIQSSFCLTNNKIGNNEDMMNRYHKPLKKSTVPMALQ
CpPT2     -MLQMQ-CLNLAPKFNP-LQSPGCSRKFASPLV--------TQRHKSSIKC-----SSQSSFSFP-NQNKITHNNDKSPYKPL----VPLALQ
CpPT3     MMLQMHSSSSFSPKYYYPLQHAGCDKTLQLPLTKVHGGLNRSESKNYAIKC-----TQSDSFYST-NKIKNNENTSSRNCKPFNKYRVAVTLQ
CmiPT1a   MLLQMNLCSSFSLKYHP-LQQNGTVKTFHSPLTQIYGLANRRESNKYSVKG-----STQSSFCLTNNKIGNNEDMMNRYHKPL-KSTVPRALQ
CmiPT1b   MLLQMNLCSSFSLKYHP-LQQNGTVKTFQSPLTQIYGLANRRESNKYSVKG-----STQSSFCLTNNKIGNNEDMMNRYHKPL-KSTVPRALQ
MePT1     -MIQMHSSSSFSPKYYYPLQQAGCVKTLQLPVTKVHGGLNRIESKNYAIKC-----TQSDSFYST-NKIRINENTSSRNCKPFNKYRVAVALQ
MePT2     MLLQMNSSASFPVKFNPLHRRPDSIKTFRSLAIQRHDLVNGRESKKYPIKCSSQRCPSQSSLYLT-NKTRTSEDVINRNHKTLNKSTVPLASQ

                                                                                1st Asp-rich motif
CpPT4     RHDNNYST---SFQDVFFRNLNALYCLTRPYTWIGTIIAITSVSFLPIQNLAVLTPTLFVEILKALVPAVLMNVYVVALNQLSDLEIDKINKP
ClPT1     NEDDINST---SFRDVLLKKLHALYVFTRPFAMIGTIVGITSIAILPLQSFADLTPKYFMEFLKALLSAVLMNNYVGTVNQVADVEIDKVNKP
CpPT1     QNENIVST---SFLDFLTKKLDAFYRLSRPYAWTSIIVGILSSSLLPIQSLADLTPTFLIEVLKPIVPSIMMNIFVVAINQLSDIAIDKVNKP
CpPT2     SEDDNKPAAAPSFLEVVKRKLNAISHVTRYYAQINIIVSVTSVCFLPVQSLSQVTPAFLMGVLKAVVAQIFMNISLCSLNQICDVEIDKINKP
CpPT3     NEDDINST---SFWDVLLKKLHALYVFTRPFAMIGTIVGITSIAILPLQSFADLTPKYFMEFLKALLSAVLMNNYVGTVNQVADVEIDKVNKP
CmiPT1a   QNENIVWT---NFLDFLTKKLDAFYRLSRPYAWTSIIVGILSSSLLPIQSLADLTPTFLIEVLKPIVPTIMMNIFVVAINQLSDIAIDKVNKP
CmiPT1b   QNENIVWT---NFLDFLTKKLDAFYRLSRPYAWTSIIVGILSSSLLPIQSLADLTPTFLIEVLKPIVPTIMMNIFVVAINQLSDIAIDKVNKP
MePT1     NEDDINST---SFQDALMKKLNALYRFTRPFAVIGTIVGITSISVLPLQSFADLTPRYFMGLLKTLLSASLMNTYVGTLNQLTDVEIDKVNKP
MePT2     ---KIFST---SFLENLNKKLDALYRLTRPYTWIGIIVGILSVSLLPVQSLADLTPTFFMEVLKSTMPAILMNVFVTAINQLSDVEIDKVNKP

CpPT4     DFSTGTGVAVYLVSSFTSLAMGVMSRSPPLLLALIIWFLLGTAYSIQLPFLRWKGHPFLAPTSMMILMGLVYQLLFHIHIQKYVLARPVIITK
ClPT1     DLSVGTGLAITLILSLTSLAIALSLQSPPLIFGLIVWFLLGTAYSVDLPFLRWKTNPFLAGMCMVIVFGLVYQFSFFIHFQKYVLGRPVVITR
CpPT1     DISIGEAIAIAVISTLTSLAMGVMLRSPPLVIALILRCILGAAYSIDLPLLRWKASPLMAAVAIVIGNGINNVLPYFLHVQKYVLGRPVVFTK
CpPT2     ELSMGTGIAICAGSALLSLALAFLSGSPAVLCAVIAWGLTGAAYSVPLPFLRWKSHTFMAPFTLVILMGLILQIPYFIHSQTYLLGKPFEVTG
CpPT3     DLSMGTGLAITLTLSLTSLAIALSLQSPPLIFGLIVWFLLGTAYSVDLPFLRWKTKPFLAGMCMVTVFGLVYQFSFFIHFQKYVLGRPVVITR
CmiPT1a   DISIGEAIAIAVISTLTSLAMGVMLRSPPLVIALILRCILGAAYSIDLPLLRWKASPLMAAVAIVIGNGINNVLPYFLHVQKYVLGRPVVFTK
CmiPT1b   DISIGEAIAIAVISTLTSLAMGVMLRSPPLVIALILRCILGAAYSIDLPLLRWKASPLMAAVAIVIGNGINNVLPYFLHVQKYVLGRPVVFTK
MePT1     DLSMGTGLAISLIMSLMSLAIALSLQSPPLVFGLIVWFLLGTAYSVDLPFLRWKSNSFLAGMCMVIVFGLVYQFPFFIHFQKYALGRPIIITR
MePT2     DFSIGEGVAIAVLSTLTSLAMGVMLRSPPLVIGLITWWIVGAAYSIDLPLLRWKGNPLMAAVTIMILNGLLLQFPYFVHVQKYVLGRRLEFTK

             2nd Asp-rich motif
CpPT4     FMSVFSFANGLLKDVPDVDGDREFGIQTLSVIIGKEKVFWLGVYMMLTAYGAAILVGASSSFLLSKLVTIISHSILAFILWLRAQTVDLSNKA
ClPT1     IISTISAVMSLLKDIPDEDGDKQFGYQSISSKLGKENVLRLCVYALFFAYGVAVIVGASSSFQLGKLVSIIGHSTLAFLLWLRAQTVDLSNNA
CpPT1     FSAIFSIVLSFLKDIPDVEGDKKSGIRTLPVILGKERVLSMSTGILLMAYASAALAGVFSPILLCKLVTMIGHSVLGFILWSKAQTVDLSNAK
CpPT2     IMSIYAFVNGLLKDLPDVEGDKAFGMQTLCVLLGKEKVLPLCVNMMLLGFGGAVLAGATSTLMISKLVTIIGHIILALMMWLRSRKVDLDNFD
CpPT3     IISTISAVMSLLKDIPDEDGDKQFGFQSISSKLGKENVLRLCVYALFFAYGVSVIVGASSSFQLVKLVSIIGHSTLAFLLWLRAQTVDLSNNA
CmiPT1a   FSAIFSIVLSFLKDIPDVEGDKKSGIRTLPVILGKERVLSMSTGILLMAYASAALAGVFSPILLCKLVTMIGHSVLGFILWSKAQTVDLSNAK
CmiPT1b   FSAIFSIVLSFLKDIPDVEGDKKSGIRTLPVILGKERVLSMSTGILLMAYASAALAGVFSPILLCKLVTMIGHSVLGFILWSKAQTVDLSNAK
MePT1     IICTISAVMSLLKDIPDEEGDKKYGIQSMSSKLGKEKVLWLCVYTLSFAFGAAVIVGASSSFLLGKLVGVIGHSTLALVLWLRAQTVDLSNNA
MePT2     FMGIFNIAIAFVKDLPDVEGDKEFGLRTLPVILGKEKVFSISVNMLLMAYGAAVLTGASSPFLLCKLVSMIGHSALGFILWRQAQTIDLSDTK

CpPT4     FIWKLHYVEYLLIHFVR
ClPT1     FVWKLFYGEYLLIHFLR
CpPT1     FIFQLYYTEFFLMHFVR
CpPT2     FLWQLNYVEYILIHFLR
CpPT3     FIWKLFYAEYLLIHFLR
CmiPT1a   FIFQLYYTEFFLMHFVR
CmiPT1b   FIFQLYYTEFFLMHFVR
MePT1     FVWKLFYAEYLLIHFLR
MePT2     FIFKLYYAEFFLMHFIR
```

## E

| PT | ClPT1 | CpPT1 | CpPT2 | CpPT3 | CmiPT1a | CmiPT1b | MePT1 | MePT2 |
|---|---|---|---|---|---|---|---|---|
| **CpPT4 (%)** | 57.9 | 50.2 | 47.4 | 58.3 | 49.7 | 49.7 | 57 | 52.5 |

**Figure S2.** *In silico* analysis of the CpPT4 polypeptide. *–continued*
**(A)** Multiple transmembrane regions of CpPT4 predicted by DeepTMHMM. **(B)** Probability of cleavage site of CpPT4 predicted by TargetP-2.0. **(C)** Transit peptide of CpPT4 predicted by DeepLoc. **(D)** MUCSLE multiple alignment of CpPT4 and Rutaceae PTs. The first and second aspartate-rich motifs are highlighted by the boxes. **(E)** Amino acid sequence identity among the PTs of Rutaceae.

**Coumarins (simple coumarins and furanocoumarins)**

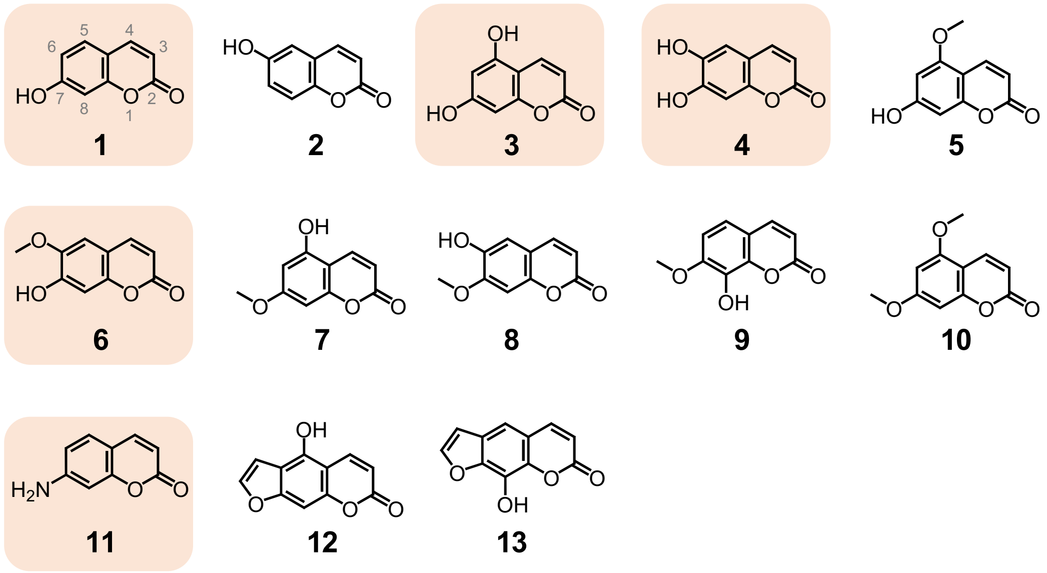


**Non-coumarin molecules**

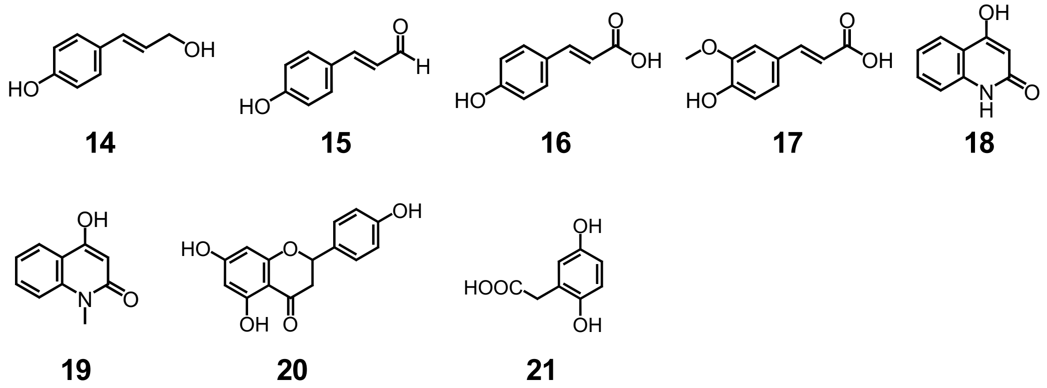


**Figure S3.** The chemical structure of prenyl acceptor substrates tested in functional screening of CpPT4.
Substrates that accept prenyl moieties are highlighted in orange. The number of carbons in umbelliferone (compound 1) is shown in grey.

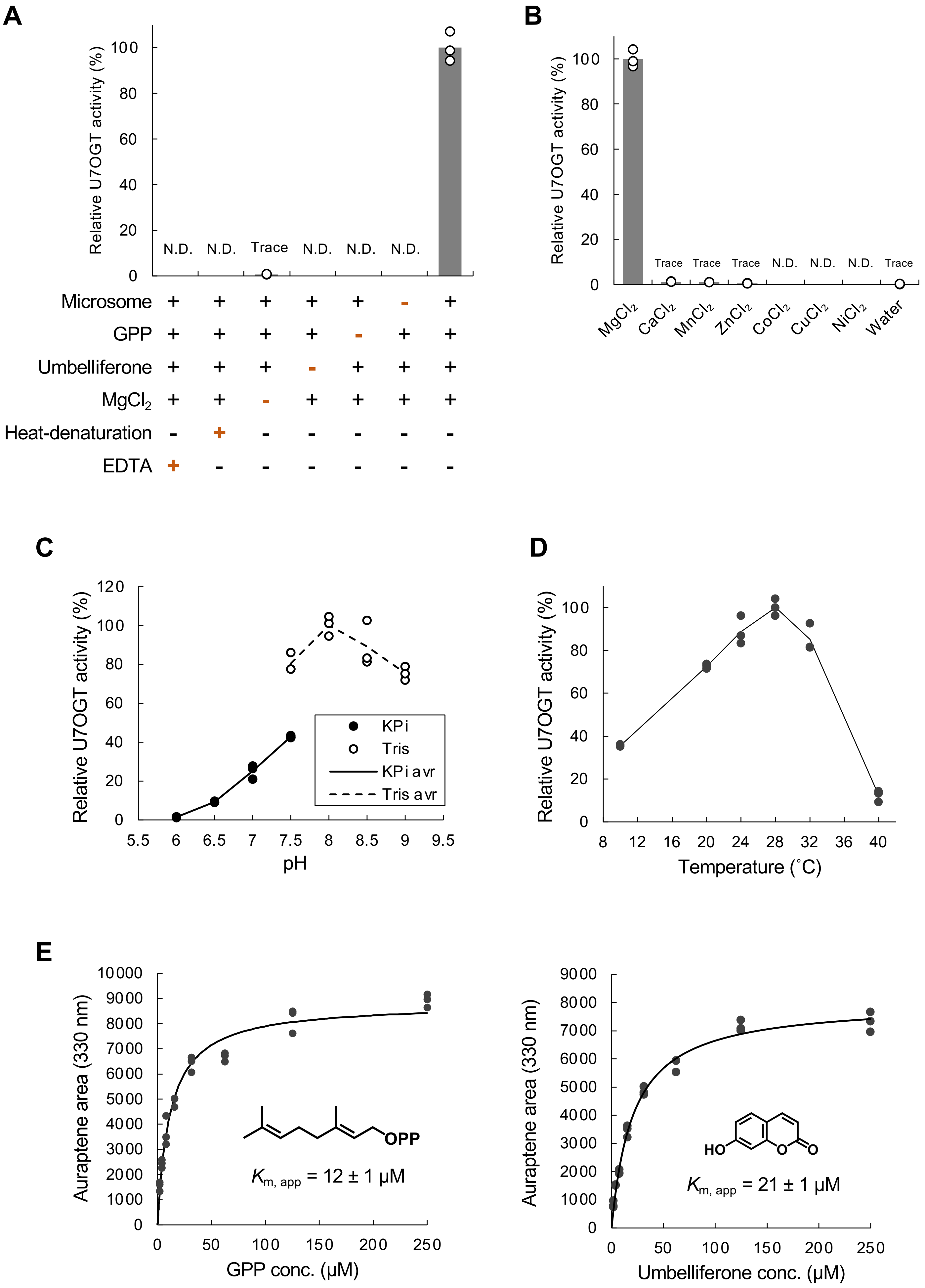


**Figure S4.** Characterization of the U7OGT activity of CpPT4.

**(A)** Negative control assays for U7OGT activity. From left to right: incubations performed in the presence of EDTA, with microsomes after heat denaturation (95 °C for 25 min), in the absence of $MgCl_2$, umbelliferone, GPP, or microsomes, and the full assay. U7OGT activities are shown relative to the mean U7OGT activity in the full assay. The bars represent the means of three independent experiments (n = 3). N.D., not detected. **(B)** Divalent cation dependence of U7OGT activity. Incubation was performed in the presence of $MgCl_2$, $MnCl_2$, $CaCl_2$, $CoCl_2$, $ZnCl_2$, or water. U7OGT activities are shown relative to the mean U7OGT activity with $MgCl_2$. The bars represent the means of three independent experiments (n = 3). N.D., not detected. **(C)** pH dependency. U7OGT activity was measured at pH 6.0 to 9.0 using KPi buffer (pH 6.0 to 7.5) and Tris-HCl buffer (pH 7.5 to 9.0). The results of three independent experiments (n = 3) are shown relative to the mean U7OGT activity at pH 8.0. The lines are drawn with the means. **(D)** Temperature dependency of the data. Incubations were performed at temperatures ranging from 10 to 40 °C. The results of three independent experiments are shown relative to the mean U7OGT activity at 28 °C. The line is drawn with the means. **(E)** Kinetic analysis. The apparent $K_m$ values for GPP (left) and umbelliferone (right) were calculated using the nonlinear least squares method. The concentrations of substrates ranged from 2–250 μM for GPP and 2–250 μM for umbelliferone in the presence of 250 μM co-substrate. For each condition, three independent experiments were performed (n = 3).

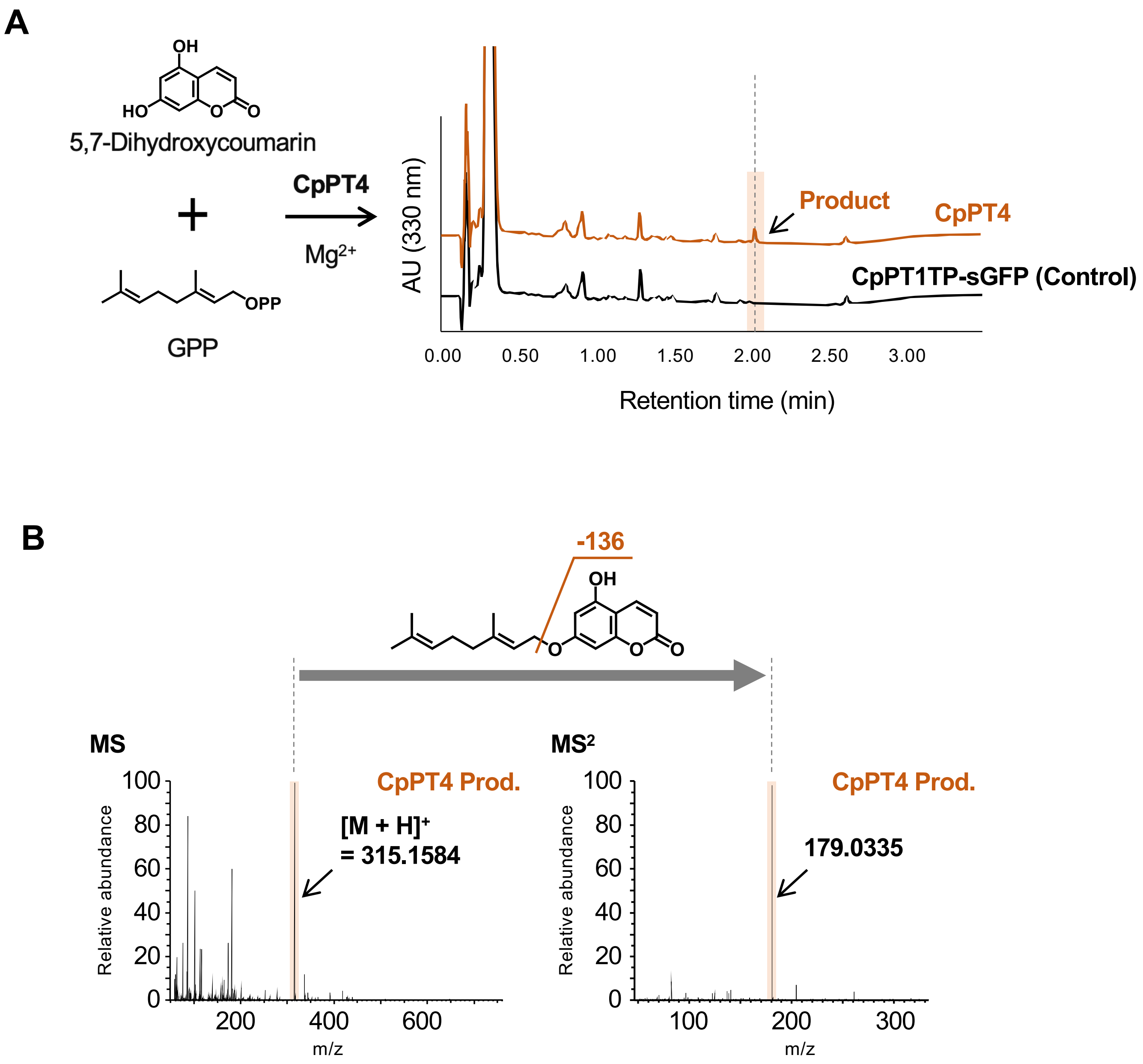


**Figure S5.** Enzymatic reactions catalyzed by recombinant CpPT4.

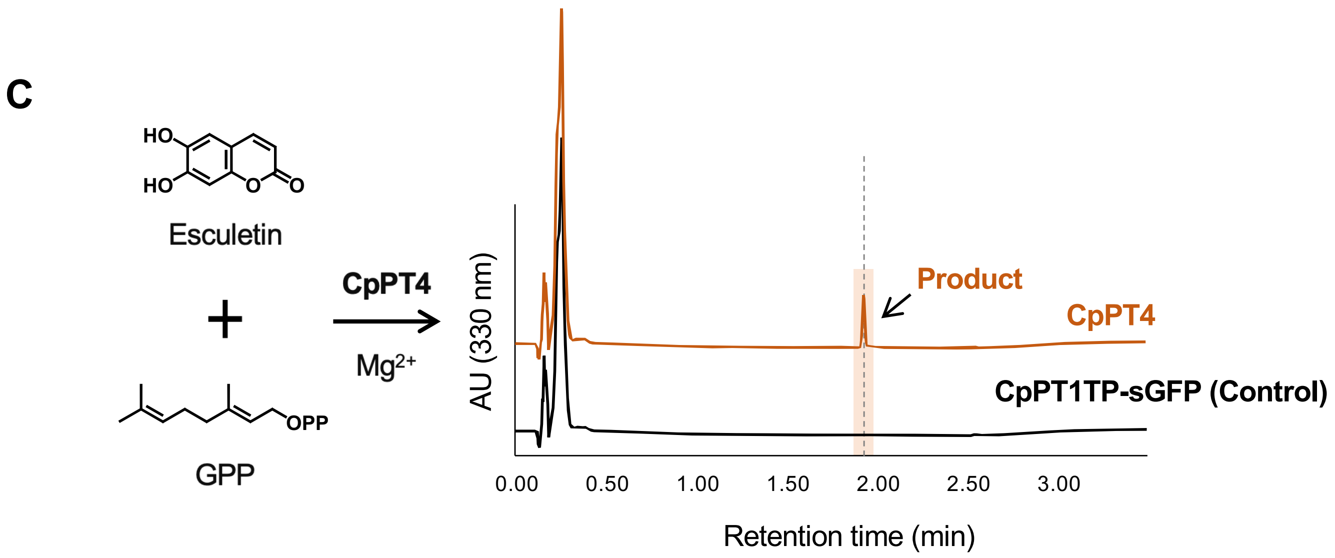


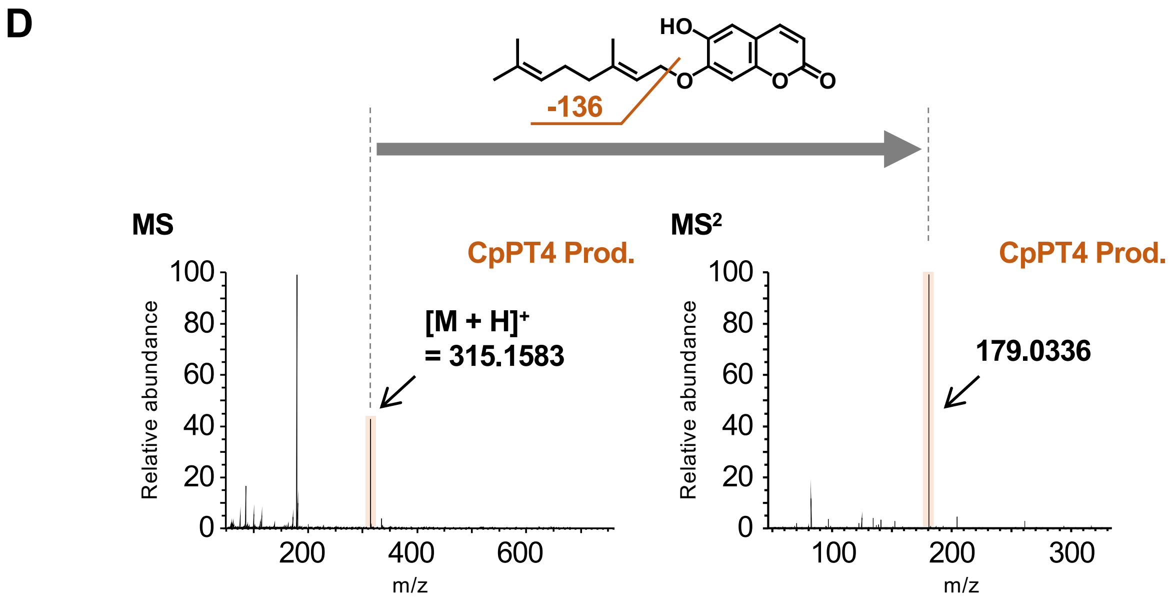


**Figure S5.** Enzymatic reactions catalyzed by recombinant CpPT4. *-continued*

E

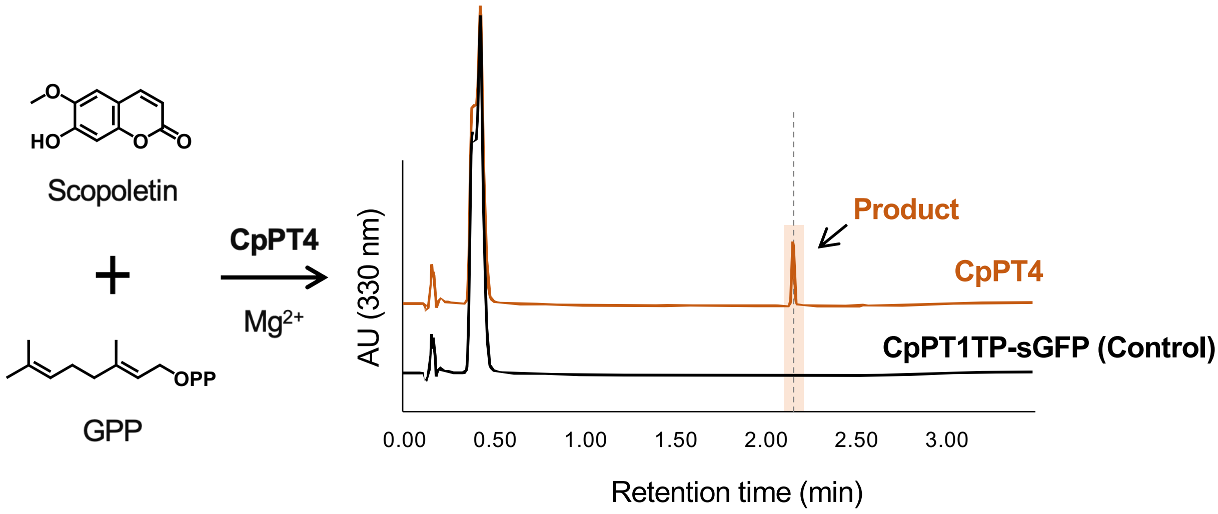


F

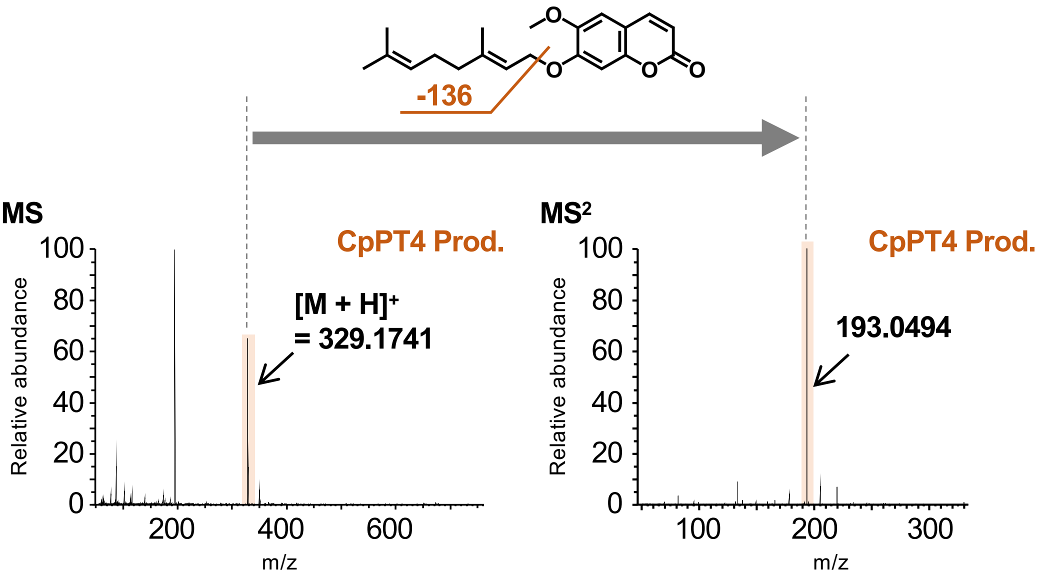


**Figure S5.** Enzymatic reactions catalyzed by recombinant CpPT4. *-continued*

G

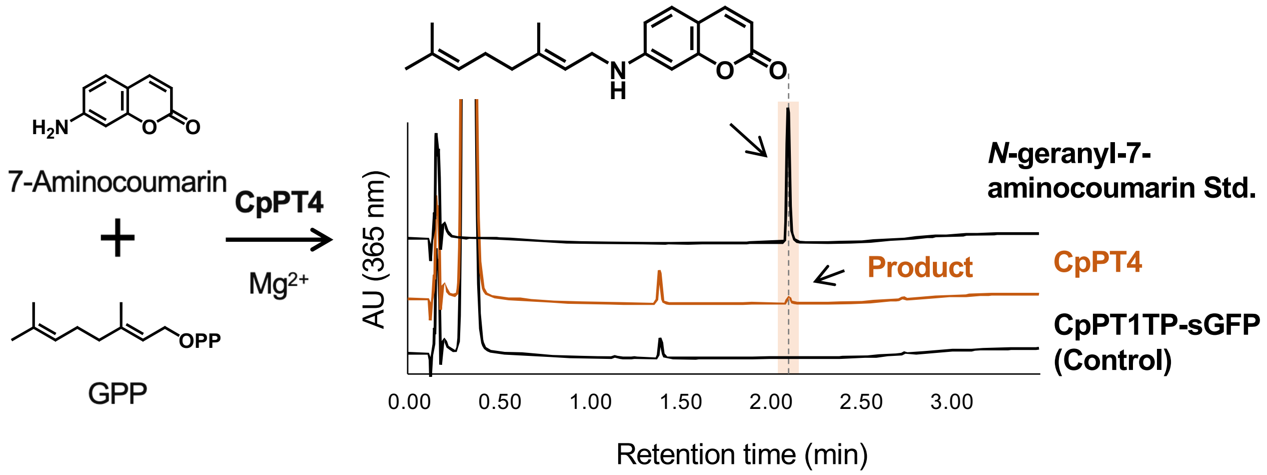


H

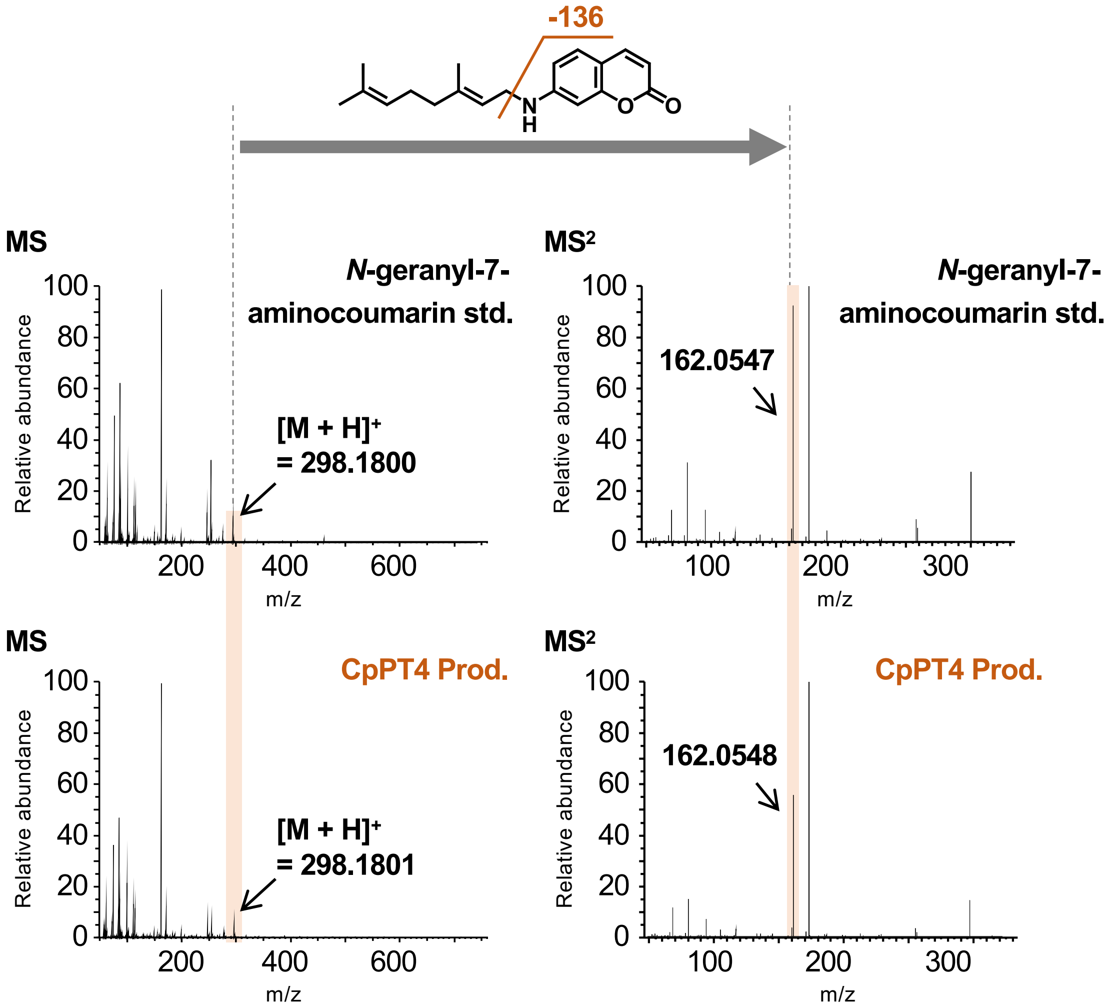


**Figure S5**. Enzymatic reactions catalyzed by recombinant CpPT4. –*continued*
**(A-H)** *O/N*-GT activities of CpPT4 toward coumarin derivatives, excluding umbelliferone.
**(A, C, E, G)** UV chromatograms of reaction mixtures containing CpPT4 and individual coumarins—**(A)** 5,7-dihydroxycoumarin (no. 3), **(C)** esculetin (no. 4), **(E)** scopoletin (no. 6), and **(G)** 7-aminocoumarin (no. 11) —in the presence of GPP as the prenyl donor. CpPT1TP-sGFP served as the negative control, and its chromatograms are shown at scales comparable to those of the corresponding CpPT4 reactions. **(B, D, F, H)** MS and MS$^2$ spectra of the products formed in **(A, C, E, G)**, respectively, acquired in the positive ion mode. Based on the MS$^2$ fragmentation patterns and substrate specificity of CpPT4 (Table S2), the products of 5,7-dihydroxycoumarin (no. 3), esculetin (no. 4), and scopoletin (no. 6) were predicted to be 7-*O*-geranylated.

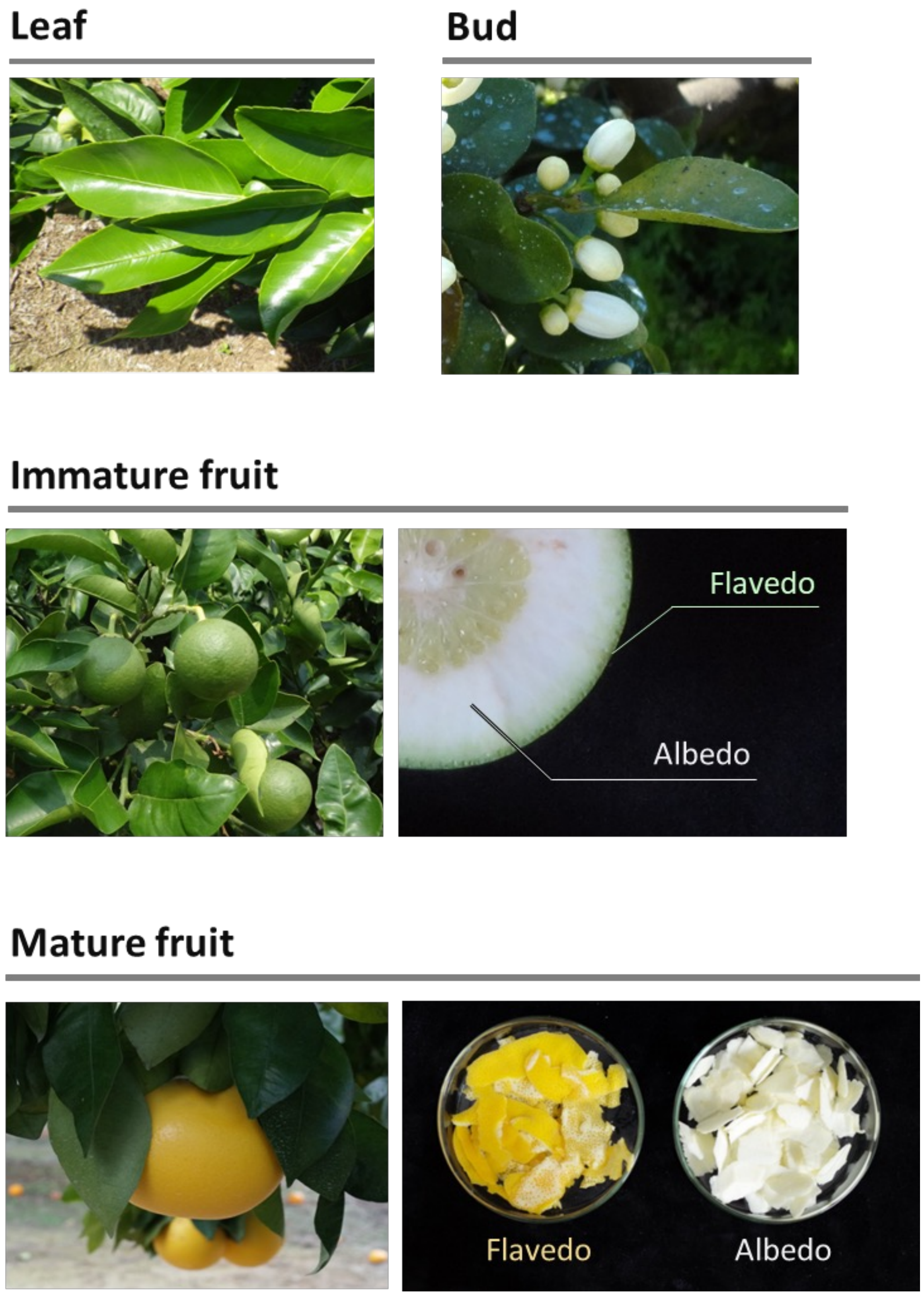


**Figure S6.** Grapefruit organs used in this study.
Leaves, buds, and immature and mature fruits of grapefruit (cv. Marsh) were harvested from the Regional Revitalization and Agricultural Research Institute farm of Kindai University. The outer peel flavedo and inner white peel albedo were collected from each fruit.

**A**

```
                           1                                                                                                     103
CpPT4                      ATGGTTCATATGCATTCATGTTTAATGTTCTCTCCAAAATATCATTATGGTTGCACTAAAACGCTGCCATCGCCTTTGTCACAGATTCATGGCGTCCTAAACA
Pummelo_Cupi_Majiayou      ATGGTTCATATGCATTCATGTTTAATGTTCTCTCCAAAATATCATTATGGTTGCACTAAAACGCTGCCATCGCCTTTGTCACAGATTCATGGCGTCCTAAACA
Pummelo_Wanbaiyou          ATGGTTCATATGCATTCATGTTTAATGTTCTCTCCAAAATATCATTATGGTTGCACTAAAACGCTGACATCGCCTTTGTCACAGATTCATGGCGTCCTAAACA
Sweet_orange               ATGGTTCATATGCATTCATGTTTAATGTTCTCTCCAAAATATCATTATGGTTGCACTAAAACGCTGCCATCGCCTTTGTCACAGATTCATGGCGTCCTAAACA
Pure_mandarin              ATGGTTCATATGCATTCATGTTTAATGTTCTCTCCAAAATATCATTATGGTTGCACTAAAACGCTGCCATCGCCTTTGTCACAGATTCATGGCGTCCTAAACA
Clementine                 ATGGTTCATATGCATTCATGTTTAATGTTCTCTCCAAAATATCATTATG------------------------------------------------------

                           104                                                                                                   206
CpPT4                      CAACTCTAAGGAAAAAGTACTCCATCAAATGCCCCCGAAATTCATTTAATTTAACCAATAAAATTAGAAGCTGTAAACAGTTCAACAAGTATCGTTGTCTTTC
Pummelo_Cupi_Majiayou      CAACTCTAAGGAAAAAGTACTCCATCAAATGCCCCCGAAATTCATTTAATTTAACCAATAAAATTAGAAGCTGTAAACAGTTCAACAAGTATCGTTGTCTTTC
Pummelo_Wanbaiyou          CAACTCTAAGGAAAAAGTACTCCATCAAATGCCCCCGAAATTCATTTAATTTAACCAATAAAATTAGAAGCTGTAAACAGTTCAACAAGTATCGTTGTCTTAC
Sweet_orange               CAACTCTAAGGAAAAAGTACTCCATCAAATGCCCCCGAAATTCATTTAATTTAACCAATAAAATTAGAAGCTGTAAACAGTTCAACAAGTATCGTTGTCTTTC
Pure_mandarin              CAACTCTAAGGAAAAAGTACTCCATCAAATGCCCCCGAAATTCATTTAATTTAACCAATAAAATTAGAAGCTGTAAACAGTTCAACAAGTATCGTTGTCTTTC
Clementine                 -------------------------------------------------------------------------------------------------------

                           207                                                                                                   309
CpPT4                      TACCTTACAAGATGGCTACAAACCAAACCCCCGACATGATAATAATTATTCGACAAGCTTTCAAGATGTTTTCTTCAGGAACTTAAATGCACTCTACTGTCTT
Pummelo_Cupi_Majiayou      TACCTTACAAGATGGCTACAAACCAAACCCCCGACATGATAATAATTATTCGACAAGCTTTCAAGATGTTTTCTTCAGGAACTTAAATGCACTCTACTGTCTT
Pummelo_Wanbaiyou          TACCTTACAAGATGGCTACAAACCAAACCCCCTACATGATAATAATTATTCGACAAGCTTTCAAGATGTTTTCTTCAGGAACTTAAATGCACTCTACTGTCTT
Sweet_orange               CACCTTACAAGATGGTTACAAACCAAACACCCTACGTGATAATAGTAATTCGACAAGCTTTCAAGATGTTTCCTTCAGGAACTTAAATGCACTCTACTGTCTT
Pure_mandarin              CACCTTACAAGATGGTTACAAACCAAACACCCTACGTGATAATAGTAATTCGACAAGCTTTCAAGATGTTTCCTTCAGGAACTTAAATGCACTCTACTGTCTT
Clementine                 -------------------------------------------------------------------------------------------------------

                           310                                                                                                   412
CpPT4                      ACTCGTCCCTACACTTGGATTGGAACTATTATAGCGATAACATCGGTTTCTTTTCTTCCAATACAAAATCTTGCTGTTTTGACTCCCACATTATTTGTTGAAA
Pummelo_Cupi_Majiayou      ACTCGTCCCTACACTTGGATTGGAACTATTATAGCGATAACATCGGTTTCTTTTCTTCCAATACAAAATCTTGCTGTTTTGACTCCCACATTATTTGTTGAAA
Pummelo_Wanbaiyou          ACTCGTCCCTACACTTGGATTGGAACTATTATAGCGATAACATCGGTTTCTTTTCTTCCAATACAAAATCTTTCTGTTTTGACTCCCACATTATTTGTTGAAA
Sweet_orange               ACTCGTCCCTACACTTGGATTGGAACTATTATAGCGATAACATCGGTTTCTTTTCTTCCAATACAAAATCTTGCTGTTTTAACTCCCACATTATTTGTTGAAA
Pure_mandarin              ACTCGTCCCTACACTTGGATTGGAACTATTATAGCGATAACATCGGTTTCTTTTCTTCCAATACAAAATCTTGCTGTTTTGACTCCCACATTATTTGTTGAAA
Clementine                 -------------------------------------------------------------------------------------------------------

                           413                                                                                                   515
CpPT4                      TCTTGAAGGCTTTGGTGCCCGCTGTACTAATGAACGTTTACGTGGTGGCCCTAAACCAGTTATCTGACCTTGAAATAGACAAGATTAACAAGCCCGATCTTCC
Pummelo_Cupi_Majiayou      TCTTGAAGGCTTTGGTGCCCGCTGTACTAATGAACGTTTACGTGGTGGCCCTAAACCAGTTATCTGACCTTGAAATAGACAAGATTAACAAGCCCGATCTTCC
Pummelo_Wanbaiyou          TCTTGAAGGCTTTGGTGCCCGCTGTACTAATGAACGTTTACGTGGTGGCCCTAAACCAGTTATCTGACCTTGAAATAGACAAGATTAACAAGCCCGATCTTCC
Sweet_orange               TCTTGAAGGCTTTGGTGCCCGCTGTACTAATGAACGTTTACGTGGTGGCCCTAAACCAGTTATCTGACCTTGAAATAGACGAGATTAACAAGCCCGATCTTCC
Pure_mandarin              TCTTGAAGGCTTTGGTGCCCGCTGTACTAATGAACGTTTACGTGGTGGCCCTAAACCAGTTATCTGACCTTGAAATAGACAAGATTAACAAGCCCGATCTTCC
Clementine                 -------------------------------------------------------------------------------------------------------
```

**Figure S7.** The nucleotide sequences of *CpPT4* orthologs in *Citrus*.

**A**

```
                          516                                                                                                   618
CpPT4                     ACTTGCTTCTGGTGACTTCTCAACGGGAACTGGAGTTGCAGTTTATTTAGTATCTTCATTCACGAGTCTGGCTATGGGAGTTATGTCTCGATCTCCACCTCTT
Pummelo_Cupi_Majiayou     ACTTGCTTCTGGTGACTTCTCAACGGGAACTGGAGTTGCAGTTTATTTAGTATCTTCATTCACGAGTCTGGCTATGGGAGTTATGTCTCGATCTCCACCTCTT
Pummelo_Wanbaiyou         ACTTGCTTCTGGTGACTTCTCAACGGGAACTGGAGTTGCAGTTTATTTAGTATCTTCATTCACGAG--TGGCTATGGGAGTTATATCTCGATCTCCACCTCTT
Sweet_orange              ACTTGCTTCTGGTGACTTCTCAACGGGAACTGGATTTGCAGTTTATTTAGTATCTTCATTCACGAGTCTGGCTATGGGAGTTATGTCTCGATCTCCACCTCTT
Pure_mandarin             ACTTGCTTCTGGTGACTTCTCAACGGGAACTGGAGTTGCAGTTTATTTAGGATCTTCATTCACGAGTCTGGCTATGGGAGTTATGTCTCGATCTCCACCTCTT
Clementine                -------------------------------------------------------------------------------------------------------

                          619                                                                                                   721
CpPT4                     CTCCTGGCCCTCATCATATGGTTTTTACTCGGAACCGCTTACTCCATCCAACTTCCCTTTCTAAGATGGAAAGGACATCCATTTCTTGCACCAACCTCCATGA
Pummelo_Cupi_Majiayou     CTCCTGGCCCTCATCATATGGTTTTTACTCGGAACCGCTTACTCCATCCAACTTCCCTTTCTAAGATGGAAAGGACATCCATTTCTTGCACCAACCTCCATGA
Pummelo_Wanbaiyou         CTTCTGGCCTTCATCATCTGGTTTCTACTCGGAACCGCTTACTCCATCCAACTTCCCTTTCTAAGATGGAAAGGACATCCATTTCTTGCACCAACCTCCATGA
Sweet_orange              CTCCTGGCCCTCATCATATGGTTTTTACTCGGAACCGCTTACTCCATCCAACTTCCCTTTCTAAGATGGAAAGGACATCCATTTCTTGCACCAACCTCCATGA
Pure_mandarin             CTCCTGGCCCTCATCATATGGTTTTTACTCGGAACCGCTTACTCCATCCAACTTCCCTTTCTAAGATGGAAAGGACATCCATTTCTTGCACCAACCTCCATGA
Clementine                -------------------------------------------------------------------------------------------------------

                          722                                                                                                   824
CpPT4                     TGATTTTGATGGGACTTGTATATCAATTGCTATTCCACATACACATACAGAAATACGTGCTTGCAAGACCTGTAATAATTACAAAACCACTGGTGTTTGCTAC
Pummelo_Cupi_Majiayou     TGATTTTGATGGGACTTGTATATCAATTGCTATTCCACATACACATACAGAAATACGTGCTTGCAAGACCTGTAATAATTACAAAACCACTGGTGTTTGCTAC
Pummelo_Wanbaiyou         TGATTTTGATGGGACTTGTATATCAATTGCTATTCTACATACACATCCAGAAATACGTGCTTGCAAGACCTGTAATAATTACAAAACCACTGGTGTTTGCTAC
Sweet_orange              TGATTTTGATGGGACTTGTATATCAATTGCTATTCTACATACACATCCAGAAATACGTGCTTGCAAGACCTGTAATAATTACAAAACCACTGGTGTTTGCTAC
Pure_mandarin             TGATTTTGATGGGACTTGTATATCAATTGCTATTCTACATACACATCCAGAAATACGTGCTTGCAAGACCTGTAATAATTACAAAACCACTGGTGTTTGCTAC
Clementine                -------------------------------------------------------------------------------------------------------

                          825                                                                                                   927
CpPT4                     CACTTTCATGTCTGTATTCTCC--TTTGCAAATGGCCTTCTCAAGGATGTACCTGATGTGGATGGCGACAGAGAATTTGGCATTCAGACGCTCAGTGTTATCA
Pummelo_Cupi_Majiayou     CACTTTCATGTCTGTATTCTCC--TTTGCAAATGGCCTTCTCAAGGATGTACCTGATGTGGATGGCGACAGAGAATTTGGCATTCAGACGCTCAGTGTTATCA
Pummelo_Wanbaiyou         CACTTTCATGTCCGTATTCTCC--TTTGCAAATGGCCTTCTCAAGGATGTACCTGATGTGGATGGCGACAGAGAATTTGGCATTCAGACGCTCAGTGTTATCA
Sweet_orange              CACTTTCATATCTGTATTCTCTCGTTTGCAAATGGCCTTCTCAAGGATGTACCTGATGTGGATGGCGACAGAGAATTTGGCATTCAGACGCTCAGTGTTATCA
Pure_mandarin             CACTTTCATATCTGTATTCTCTCGTTTGCAAATGGCCTTCTCAAGGATGTACCTGATGTGGATGGCGACAGAGAATTTGGCATTCAGACGCTCAGTGTTATCA
Clementine                -------------------------------------------------------------------------------------------------------

                          928                                                                                                   1030
CpPT4                     TAGGAAAAGAAAAAGTATTTTGGCTTGGTGTTTATATGATGCTAACTGCCTATGGAGCTGCTATATTAGTTGGAGCTTCTTCATCTTTTCTGCTAAGCAAGCT
Pummelo_Cupi_Majiayou     TAGGAAAAGAAAAAGTATTTTGGCTTGGTGTTTATATGATGCTAACTGCCTATGGAGCTGCTATATTAGTTGGAGCTTCTTCATCTTTTCTGCTAAGCAAGCT
Pummelo_Wanbaiyou         TAGGAAAAGAAAAAGTATTTTGGCTTGGTGTTTATATGATGTTAACTGCCTATGGAGCTGCTATATTAGTTGGAACTTCTTCATCTTTTCTGCTAAGCAAGCT
Sweet_orange              TAGGAAAAGAAAAAGTATTTTGGCTTGGTGTTTATATGATGTTAACTGCCTATGGAGCTGCTATATTAGTTGGAGCTTCTTCATCTTTTCTGCTAAGCAAGCT
Pure_mandarin             TAGGAAAAGAAAAAGTATTTTGGCTTGGTGTTTATATGATGTTAACTGCCTATGGAGCTGCTATATTAGTTGGAGCTTCTTCATCTTTTCTGCTAAGCAAGCT
Clementine                -------------------------------------------------------------------------------------------------------

                          1031                                                                                                  1133
CpPT4                     TGTCACTATAATTTCTCACAGTATACTTGCTTTCATTCTGTGGCTTCGAGCCCAAACTGTTGATCTCTCCAATAAAGCGTCAACACTTTCTTTTTACATGTTC
Pummelo_Cupi_Majiayou     TGTCACTATAATTTCTCACAGTATACTTGCTTTCATTCTGTGGCTTCGAGCCCAAACTGTTGATCTCTCCAATAAAGCGTCAACACTTTCTTTTTACATGTTC
Pummelo_Wanbaiyou         TGTCACTATAATTTCTCACAGTATACTTGCTTTCATTCTGTGGCTTCGAGCCCAAACTGTTGATCTCTCCAATAAAGCATCAACACTTTCTTTTTACATGTTC
Sweet_orange              TGTCACTATAATTTCTCACAGTATACTTGCTTTCATTCTGTGGCGTCGAGCCCAAACTGTTGATCTCTCCAATAAAGCATCAACACTTTCTTTTTACATGTTC
Pure_mandarin             TGTCACTATAATTTCTCACAGTATACTTGCTTTCATTCTGTGGCGTCGAGCCCAAACTGTTGATCTCTCCAATAAAGCATCAACACTTTCTTTTTACATGTTC
Clementine                -------------------------------------------------------------------------------------------------------

                          1134                                            1184
CpPT4                     ATTTGGAAGTTACACTACGTTGAATACCTCCTTATTCATTTTGTACGTTGA
Pummelo_Cupi_Majiayou     ATTTGGAAGTTACACTACGTTGAATACCTCCTTATTCATTTTGTACGTTGA
Pummelo_Wanbaiyou         ATTTGGAAGTTACACTACGTTGAATACCTCCTTATTCATTTTGTACGTTGA
Sweet_orange              ATTTGGAAGTTACACTACGTTGAATACCTCCTTATTCATTTTGTGCGTTGA
Pure_mandarin             ATTTGGAAGTTACACTGCGTTGAATACCTCCTTATTCATTTTGTGCGTTGA
Clementine                ---------------------------------------------------
```

**Figure S7.** The nucleotide sequences of *CpPT4* orthologs in *Citrus*. *-continued*

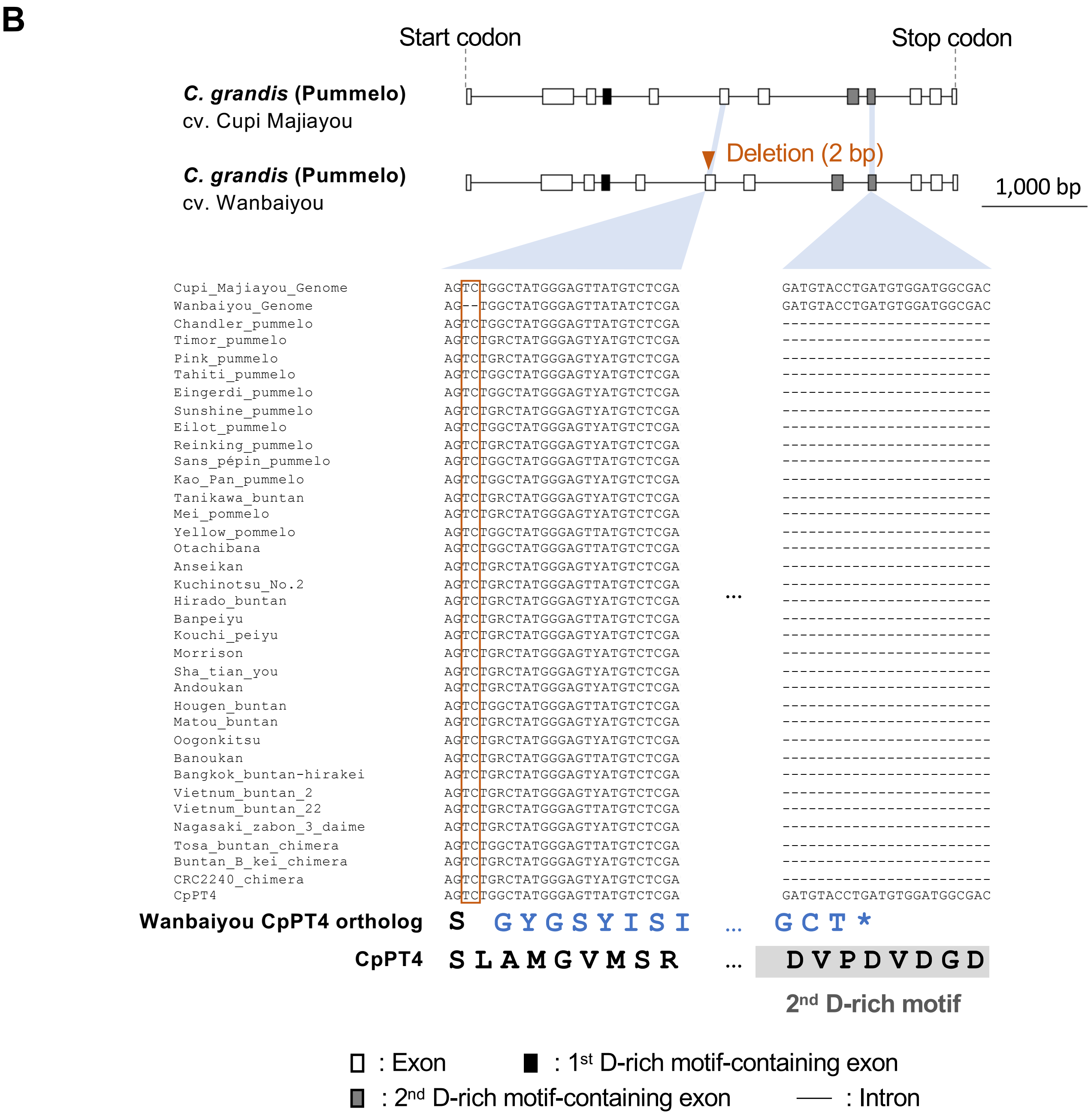


**Figure S7.** The nucleotide sequences of *CpPT4* orthologs in *Citrus*. *-continued*

**C**

```
                    1                                                                                                     104
CpPT4               ATGGTTCATATGCATTCATGTTTAATGTTCTCTCCAAAATATCATTATGGTTGCACTAAAACGCTGCCATCGCCTTTGTCACAGATTCATGGCGTCCTAAACAC
Atalantia_buxifolia ATGGTTCACATGCATTCATGTTTAACGTTCTCTCCAAAATATCATTATGGTTGCATTAAAACGCTGCAATCGCCTTCGTCACAGATTCATGGCGTCCTAAACAC
Fortunella_hindsii  ATGGTTCATATGCATTCATGTTTAATGTTCTCTCCAAAATATCATTATGGTTGCACTAAAACGCTGCAATCGCCTTTGTCACAGATTCATGGCGTCCTAAACAC
Poncirus_trifoliata ATGGTTCTTATGCATTCATGTTTAATGTTCTCTCCAAAATATCATTATGGTTGCACTAAAACGCTGCCATCGCCTTTGTCACAGATTCATGGCGTCCTAAACAC
C_ichangensis       ATGGTTCATATGCATTCATGTTTAATGTTCTCTCCAAAATATCATTATGGTTGCACTAAAACGCTGCAATCGCCTTTGTCACAGATTCATGGCGTCCTAAACAC
C_medica            ATGGTTCATATGCACTCATGTTTAATGTTCTCTCCAAAATATCATTATGGTTGCACTAAAACGCTGCAATCGCCTTTGTCACAGATTCATGGCGTCCTAAACAC

                    105                                                                                                   208
CpPT4               AACTCTAAGGAAAAAGTACTCCATCAAATGCCCCCGAAATTCATTTAATTTAACCAATAAAATTAGAAGCTGTAAACAGTTCAACAAGTATCGTTGTCTTTCTA
Atalantia_buxifolia AACTCTAAGCAAAAAGTACTCCATCAAATGCCCCCGAAATTCATTTAATTTAACCTATAAGATTAGAAGCTGTAAACAGTTCAAAAAGTATCGTTGTCTTCCCA
Fortunella_hindsii  AACTCTAAGCAAAAAGTACTCCATCAAATGCCCCCGAAATTCATTTAATTTAACCAATAAAATTAGAAGCTGTAAACAGTTCAACAAGTATCGTTGTCTTTCCA
Poncirus_trifoliata AACTCTAAGGAAAAAGTACTCCATCAAATACCCCCGAAATTCATTTAATTTAACCAATAAAATTAGAAGCTGTAAACAGTTCAACAAGTATCGTTGTCTTTCTA
C_ichangensis       AACTCTAAGCAAAAAGTACTCCATCAAATGCCCCCGAAATTCATTTAATTCAACCAATAAAATTAGAAGCTGTAAACAGTTCAACAAGTATCGTTGTCTTTCCA
C_medica            AACTCTAAGCAAAAGGTACTCCATCAAATGCCCCCGAAATTCATTTAATTTAACCAATAAAATTAGAAGCTGTAAGCAGTTCAACAAGTATCGTTGCCTTTCCA

                    209                                                                                                   312
CpPT4               CCTTACAAGATGGCTACAAACCAAACCCCCGACATGATAATAATTATTCGACAAGCTTTCAAGATGTTTTCTTCAGGAACTTAAATGCACTCTACTGTCTTACT
Atalantia_buxifolia CCTTACAAGATGGTTACAAACCAAACCCCCTACATGATAATAATTATTCGACAAGCTTTCAAGATGTTTTCTTCAGGAACTTAAATGCACTCTACTGTCTTACT
Fortunella_hindsii  CCTTACAAGATGGTTACAAACCAAACACCCTACGTGATAATAATAATTCGACAAGCTTTCAAGATGTTTCCTTCAGGAACTTAAATGCACTCTACTGTCTTACT
Poncirus_trifoliata CCTTACAAGATGGCTACAAACCAAACCCCCTTCATGATAATAATTATTCAACAAGCTTTCAAGATGTTTTCTTCAGAAACTTAAATGCACTCTACTGTCTTACT
C_ichangensis       CCTTACAAGATGGTTACAAACCAAACACCCTACATGATAATAATAATTCGACAAGCTTTCAAGATGTTTCCTTCAGGAACTTAAATGCACTCTACTGTCTTACT
C_medica            CCTTACAAGATGGTTACAAACCAAACACCCTACGTGATAATAATAATTCGACAAGCTTTCAAGATGTTTCCTTCAGGAACTTAAATGCACTCTTCTGTCTTACT

                    313                                                                                                   416
CpPT4               CGTCCCTACACTTGGATTGGAACTATTATAGCGATAACATCGGTTTCTTTTCTTCCAATACAAAATCTTGCTGTTTTGACTCCCACATTATTTGTTGAAATCTT
Atalantia_buxifolia CGTCCCTACACTTGGATTGGAACTATTATAGCGATAACATCGGTTTCTTTTCTTCCAATACAAAATCTTGCTGTTTTGACTCCCACGTTGTTTGTTGAAATCTT
Fortunella_hindsii  CGTCCCTACACTTGGATTGGAACTATTATAGCGATAACATCGGTTTCTTTTCTTCCAATACAAAATCTTGCTGTTTTGACTCCCACATTATTTGTTGAAATCTT
Poncirus_trifoliata CGTCCCTACACTTGGATTGGAACTATTATAGCGATAACATCGGTTTCTTTTCTTCCAATACAAAATCTCGCTGTTTTGACTCCCACATTATTTGTTGAAATCTT
C_ichangensis       CGTCCCTACACTTGGATTGGAACTATTATAGCGATAACATCGGTTTCTTTTCTCCCAATACAAAATCTTGCTGTTTTGACTCCCACATTATTTGTTGAAATCTT
C_medica            CGTCCCTACACTTGGATTGGAACTATTATAGCGATAACATCGGTTTCTTTTTTTCCAATACAAAATCTTGCTGTTTTGACTCCCACATTATTTGTTGAAATCTT

                    417                                                                                                   520
CpPT4               GAAGGCTTTGGTGCCCGCTGTACTAATGAACGTTTACGTGGTGGCCCTAAACCAGTTATCTGACCTTGAAATAGACAAGATTAACAAGCCCGATCTTCCACTTG
Atalantia_buxifolia GAAGGCTTTGGTGCCCTCTGTACTAATGAACATTTACGTGGTGGCCCTAAACCAGTTATCTGACCTTGAAATAGACAAGATTAACAAGCCCGATCTTCCACTTG
Fortunella_hindsii  GAAGGCTTTGGTGCCCGCTGTACTAATGAACGTTTACGTGGTGGCCCTAAACCAGTTATCTGACCTTGAAATAGACAAGATTAACAAGCCCGATCTTCCACTTG
Poncirus_trifoliata GAAGGCTTTGGTGCCCGCTGTACTAATGAACGTTTACGTGGTGGCCCTAAACCAGTTATCTGACCTTGAAATAGACAAGATTAACAAGCCCGATCTTCCACTTG
C_ichangensis       GAAGGCTTTGGTGCCCGCTGTACTAATGAACATTTACGTGGTGGCCCTAAACCAGTTATCTGACCTTGAAATAGACAAGATTAACAAGCCCGATCTTCCACTTG
C_medica            GAAGGCTTTGGTGCCCGCTGTACTAATGAACGTTTACGTGGTGGCCCTAAACCAGTTATCTGACCTTGAAATAGACAAGATTAACAAGCCCGATCTTCCACTTG

                    521                                                                                                   624
CpPT4               CTTCTGGTGACTTCTCAACGGGAACTGGAGTTGCAGTTTATTTAGTATCTTCATTCACGAGTCTGGCTATGGGAGTTATGTCTCGATCTCCACCTCTTCTCCTG
Atalantia_buxifolia CTTCTGGTGACTTCTCAACGGGAACTGGAGTCGCAGTTTATTTAGTATCTTCATTCACGAGTCTGGCAATGGGAGTTATGTCTCGATCTCTGCCTCTTCTCCTG
Fortunella_hindsii  CTTCTGGTGACTTCTCAACGGGAACTGGAGTTGCAGTTTATTTAGTATCTTCATTCACGAGTCTGGCTATGGGAGTTATGTCTCGATCTCCACCTCTTCTCCTG
Poncirus_trifoliata CTTCTGGTGAATTCTCAACGGGAACTGGAGTTGCAGTTTATTTAGTATCTTCATTCACGAGTCTGGTTATGGGAGTTATATCTCGATCTCCACCTCTTCTTCTG
C_ichangensis       CTTCTGGTGACTTCTCAACGGGAACTGGAGTTGCAGTTTATTTAGTATCTTCATTCACGAGTCTGGCTATGGGAGTTATGTCTCGATCTCCACCTCTTCTCCTG
C_medica            CTTCTGGTGACTTCTCAACGGGAACTGGAGTTGCAGTTTATTTAGTATCTTCATTCACGAGTCTGGCTATGGGAGTTTTATCTCGATCTCCACCTCTTCTTCTG
```

**Figure S7.** The nucleotide sequences of *CpPT4* orthologs in *Citrus*. *-continued*

**C**

```
                    625                                                                                                  728
CpPT4               GCCCTCATCATATGGTTTTTACTCGGAACCGCTTACTCCATCCAACTTCCCTTTCTAAGATGGAAAGGACATCCATTTCTTGCACCAACCTCCATGATGATTTT
Atalantia_buxifolia GCCCTCATCATATGGTTTTTACTCGGAACCACTTACTCCATCAAACTTCCCTTTCTCAGATGGAAAGGACATCCATTTCTTGCACCAACCTCCATGATGATTTT
Fortunella_hindsii  GCCCTCATCATATGGTTTTTACTCGGAACCGCTTACTCCATCCAACTTCCCTTTCTAAGATGGAAAGGACATCCATTTCTTGCACCAACCTCCATGATGATTTT
Poncirus_trifoliata GCCCTCATCATCTGGTTTCTACTCGGAACTGCTTACTCCATTCAACTTCCCTTTCTAAGATGGAAAGGACATCCATTTCTTGCACCAACCTCCATGATGATTTT
C_ichangensis       GCCCTCATCATATGGTTTTTACTCGGAACCGCTTACTCCATCCAACTTCCCTTTCTAAGATGGAAAGGACATCCATTTCTTGCACCAACCTCCATGATGATTTT
C_medica            GCCCTCATCATCTGGTTTTTACTCGGAACCGCTTACTCCATCCAACTTCCCTTTCTAAGATGGAAAGGACATCCATTTCTTGCACCAACCTCCATGATGATTTT

                    729                                                                                                  832
CpPT4               GATGGGACTTGTATATCAATTGCTATTCCACATACACATACAGAAATACGTGCTTGCAAGACCTGTAATAATTACAAAACCACTGGTGTTTGCTACCACTTTCA
Atalantia_buxifolia GATGGGACTTGTATATCAATTGCTATTCTACATACACATCCAGAAATACGTGCTTGGAAGACCTGTAATAATTACAAAACCACTGGTGTTTGCTACCACTTTCA
Fortunella_hindsii  GATGGGACTTGTATATCAATTGCTATTCTACATACACATCCAGAAATACGTGCTTGCAAGACCTGTAATAATTACAAAACCACTGGTGTTTGCTACCACTTTCA
Poncirus_trifoliata GATGGGACTTGTATATCAATTGCTATTCTACATACACATCCAGAAATACGTGCTTGCAAGACCTGTAATAATTACAAAACCACTGGTGTTTGCTACCACTTTCA
C_ichangensis       GATGGGACTTGTATATCAATTGCTATTCTACATACACATCCAGAAATACGTGCTTGCAAGACCTGTAATAATTACAAAACCACTGGTGTTTGCTACCACTTTCA
C_medica            GATGGGACTTGTATATCAATTGCTATTCTACATACACATCCAGAAATACGTGCTTGCAAGACCTGTAATAATTACAAAACCACTGGTGTTTGCTACCGCTTTCA

                    833                                                                                                  936
CpPT4               TGTCTGTATTCTCCTTTGCAAATGGCCTTCTCAAGGATGTACCTGATGTGGATGGCGACAGAGAATTTGGCATTCAGACGCTCAGTGTTATCATAGGAAAAGAA
Atalantia_buxifolia TATCTGTATTCTCCTTTGCAAATGGCCTTCTCAAGGATGTACCTGATGTCGAGGGCGACAAGGAATTTGGCATTCAGACGCTCAGTGTTATCATAGGAAAAGAA
Fortunella_hindsii  TATCTGTATTCTCGTTTGCAAATGGCCTTCTCAAGGATGTACCTGATGTGGATGGCGACAGAGAATTTGGCATTCAGACGCTCAGTGTTATCATAGGAAAAGAA
Poncirus_trifoliata TGTCTGTATTCTCCTTTGCAAATGGCCTTCTCAAGGATGTACCTGATGTGGATGGCGACAGAGAATTTGGCATTCAGACGCTCACTGTTATCATAGGAAAAGAA
C_ichangensis       TATCTGTATTCTCGTTTGCAAATGGCCTTCTCAAGGATGTACCTGATGTGGATGGCGACAGAGAATTTGGCATTCA-ACGCTCAGTGTTATCATAGGAAAAGAA
C_medica            TGTCCGTATTCTCCTTTGCAAATGGCCTTCTCAAGGATGTACCTGATGTGGATGGCGACAGAGAATTTGGCATTCAGACGCTCAGTGTTATCATAGGAAAAGAA

                    937                                                                                                 1040
CpPT4               AAAGTATTTTGGCTTGGTGTTTATATGATGCTAACTGCCTATGGAGCTGCTATATTAGTTGGAGCTTCTTCATCTTTTCTGCTAAGCAAGCTTGTCACTATAAT
Atalantia_buxifolia AAAGTATTTTGGCTTGGTGTTTCTATGATGTTAACTGCCTATGGAGCTGCTATATTAGTTGGAGCTTCTTCATCTTTACTGCTAAGCAAGCTTGTCACTATAAT
Fortunella_hindsii  AAAGTATTTTGGCTTGGTGTTTATATGATGTTAACTGCCTATGGAGCTGCTATATTAGTTGGAGCTTCTTCATCTTTTCTGCTAAGCAAGCTTGTCACTATAAT
Poncirus_trifoliata AAAGTATTTTGGCTTGGTGTTTATATGATGTTAACTGCCTATGGAGCTGCTATATTAGTTGGAGCTTCTTCATCTTTTCTGCTAAGCAAGCTTGTCACTATGAT
C_ichangensis       AAAGTATTTTGGCTTGGTGTTTATATGATGTTAACTGCCTATGGAGCTGCTATATTAGTTGGAGCTTCTTCATCTTTTCTGCTAAGCAAGCTTGTCACTATAAT
C_medica            AAAGTATTTTGGCTTGGTGTTTATATGATGTTAGCTGCCTATGGAGCTGCTATATTGGTTGGAGCTTCTTCATCTTTTCTGCTAAGCAAGCTTGTCACTATAAT

                    1041                                                                                                1144
CpPT4               TTCTCACAGTATACTTGCTTTCATTCTGTGGCTTCGAGCCCAAACTGTTGATCTCTCCAATAAAGCGTCAACACTTTCTTTTTACATGTTCATTTGGAAGTTAC
Atalantia_buxifolia TTCTCACAGTATACTTGCTTTCATTCTGTGGCTTCGAGCCCAAACTGTTGATCTCTCCAGTAAAGCATCAACACTTTCTTTTTACATGTTCATTTGGAAGTTAC
Fortunella_hindsii  TTCTCACAGTATACTTGCTTTCATTCTGTGGCGTCGAGCCCAAACTGTTGATCTCTCCAATAAAGCATCAACACTTTCTTTTTACATGTTCATTTGGAAGTTAC
Poncirus_trifoliata TTCTCACAGTGTACTTGCTTTCATTCTGTGGCTTCGAGCCCAAACTGTTGATCTCTCCAATAAAGCATCAACACTTTCTTTTTACATGTTCATTTGGAAGTTAC
C_ichangensis       TTCTCACAGTATACTTGCTTTCATTCTGTGGCGTCGAGCCCAAACTGTTGATCTCTCCAATAAAGCATCAACACTTTCTTTTTACATGTTCATTTGGAAGTTAC
C_medica            TTCTCACAGTATACTTGCTTGCATTCTGTGGCTTCGAGCCCAAACTGTTGATCTCTCCAATAAAGCATCAACACTTTCTTTTTACATGTTCATTTGGAAGTTAC

                    1145                              1182
CpPT4               ACTACGTTGAATACCTCCTTATTCATTTTGTACGTTGA
Atalantia_buxifolia ACTATGTTGAATACCTCCTTATTCATTTTGTACGTTGA
Fortunella_hindsii  ACTACGTTGAATACCTCCTTATTCATTTTGTACGTTGA
Poncirus_trifoliata ACTACGTTGAATACCTCCTTATTCATTTTGTACGTTGA
C_ichangensis       ACTACGTTGAATACCTCCTTATTCATTTTGTACGTTGA
C_medica            ACTACGTTGAATACCTCCTTATTCATTTTGTACGTTGA
```

**Figure S7.** The nucleotide sequences of *CpPT4* orthologs in *Citrus*. *-continued*

**D**

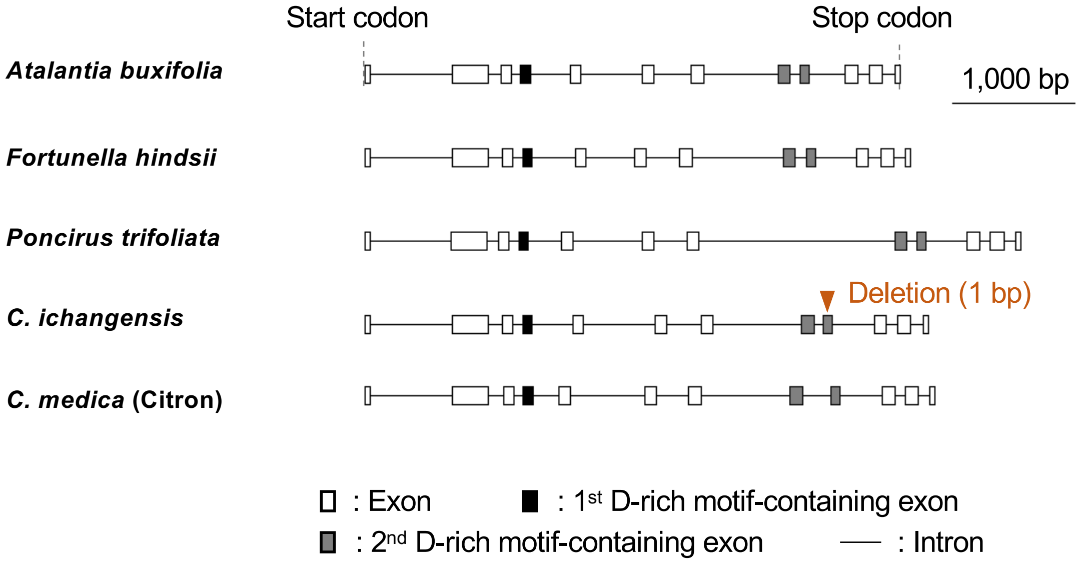


**Figure S7.** The nucleotide sequences of *CpPT4* orthologs in *Citrus*. *–continued*
**(A)** MAFFT multiple alignment of *CpPT4* CDS and its orthologs from pummelo (cv. Cupi Majiyaou and Wanbaiyou), sweet orange, pure mandarin, and clementine, respectively. Mismatches with *CpPT4* are highlighted in red. The deletion of the *CpPT4* ortholog in Wanbaiyou is shown by the red box, and the insertion in sweet orange and pure mandarin is shown by the blue box. **(B)** Gene structure model of *CpPT4* ortholog in pummelo based on (A). Alignment of the related pummelo varieties Cupi Majiayou and Wanbaiyou genomic sequences (Cupi_Majiayou_Genome and Wanbaiyou_Genome) indicated an exon-intron structure proximate to the deletion. **(C)** MAFFT multiple alignment of *CpPT4* CDS and its orthologs from *Atalantia buxifolia*, *Fortunella hindsii*, *Poncirus trifoliata*, *C. ichangensis*, and *C. medica*. Mismatches with *CpPT4* are highlighted in red. The deletion of the *CpPT4* ortholog in *C. ichangensis* is shown in the red box. **(D)** Gene structures of *CpPT4* orthologs of *Atalantia buxifolia*, *Fortunella hindsii, Poncirus trifoliata, C. ichangensis* and *C. medica* (Citron) based on their genome sequences.
